# Bayesian inference of global land plant diversification dynamics across the Phanerozoic

**DOI:** 10.64898/2026.09.29.755219

**Authors:** Xiaoya Ma, Fabien L. Condamine, Yuxuan Zhao, Yutong Zhu, Xing Zhang, Yuanhao Li, Bojian Zhong

**Affiliations:** College of Life Sciences, Ministry of Education Key Laboratory of NSLSCS, Nanjing Normal University, Nanjing, China; CNRS, Institut des Sciences de l’Évolution de Montpellier, Université de Montpellier, 475 rue du Truel, 34090 Montpellier, France

**Author notes:** Corresponding author:* Bojian Zhong.

**Keywords:** co-diversification, competitive, macroevolution, palaeoenvironment, land plants

## Abstract

Underlying diversification patterns and potential drivers of land plants remain puzzling, particularly at deep-time and global scales. Here, we reconstruct the diversification dynamics of land plants using an updated time-calibrated phylogeny comprising approximately 30,000 extant species and nearly 40,000 fossil occurrences within the Bayesian birth-death frameworks. Our analyses indicate that angiosperms exhibit the highest diversification rates, whereas ferns underwent a marked diversification increase during the Cretaceous, coincident with the rapid radiation of angiosperms. Among the major Phanerozoic mass extinctions, the Cretaceous-Paleogene event has the most significant impact on land plant diversity, with estimated genus-level losses of 13%, 14%, and 4% in bryophytes, pteridophytes, and gymnosperms. We identify a pattern consistent with competitive displacement scenario, in which pteridophyte origination rates declined as gymnosperm diversity increased, accompanying the transition from pteridophyte- to gymnosperm-dominated floras. We find that plant-insect interactions have originated as early as the emergence of pteridophytes, potentially promoting the diversification of pteridophytes and gymnosperms. During the Cretaceous, these associations shifted toward angiosperms, as indicated by a positive relationship between insect diversity and angiosperm origination rates. Finally, continental fragmentation, fluctuations in aridity, atmospheric carbon, and sedimentary dynamics showed stronger associations with the successive rise and decline of major land plant groups.

## Introduction

The emergence of land plants reshaped global ecosystems, marking the beginning of the explosion of terrestrial diversity on Earth. Land plants set the stage for the increased oxygenation of the atmosphere, greening of the continent, maintaining ecological balance, providing habitats and sustaining food chains [1,2]. Estimating a reliable pattern of land plant diversification is important for elucidating the origins and evolutionary mechanisms of biodiversity, assessing the impact of climate change on plant communities, and providing scientific foundations for climate change adaptation and management [3,4]. The interplay between origination and extinction processes determines how and when clades thrive, decline, and replace one another. Some key abiotic and biotic factors play a significant role in driving changes in diversification rates, which ultimately shape biodiversity [5,6]. Estimating origination and extinction rates and deciphering how these factors contribute to variations in rates over time are crucial for elucidating land plant diversification patterns.

The early land plant lineages had already undergone a massive radiation in biochemical, molecular, and signaling functions [7], and in morphological forms [8], providing the necessary pre-adaptations for terrestrialization. During the long evolutionary trajectory of land plants, several key events occurred, such as the colonization of terrestrial environments by plants, the flourishing of Carboniferous tropical wetlands [9], the reduction of arborescent lycophyte forests in the Pennsylvanian [10,11], and the Cretaceous explosion of angiosperms. These transitions profoundly reshaped terrestrial ecosystems and contributed to the assembly of modern floras. Consistent with this complex history, land plant diversification has been characterized by repeated shifts in origination and extinction rates, some of which coincided with major geological boundaries [5]. Recent advances in paleobotany and phylogenetics have greatly improved our understanding of land plant diversification, particularly research on the radiation of angiosperms [12–15]. A synthesis of the mesofossil floras indicated that the diversity of angiosperms and representation in fossil assemblage increased significantly around the Early-Late Cretaceous boundary [13]. Following the Cretaceous-Paleogene (K-Pg) boundary, angiosperms increasingly contributed to the restructuring of terrestrial ecosystems toward their modern configurations, and many crown-group radiations underlying present-day angiosperm diversity were established by the Neogene [16]. Global phylogenetic analyses also demonstrated that angiosperm diversity increased during the Early Cretaceous and remained stable through the remainder of the Mesozoic, until the end of the K-Pg mass extinction event [12,15]. In addition, there is so far no evidence for a mass extinction of angiosperms at the K-Pg mass extinction event. The survival of angiosperms through their higher lineages demonstrates their macroevolutionary resilience to the K-Pg mass extinction event [14]. However, despite substantial progress in resolving the evolutionary history of major plant lineages, a comprehensive understanding of diversification dynamics across the entire land plant tree of life is lacking. It remains unclear how origination and extinction rates varied through geological time, whether major evolutionary transitions coincided with shifts in diversification, and how large-scale biotic and environmental changes shaped the modern terrestrial floras.

Major land plant groups differ markedly in species richness. Angiosperms comprise approximately 90% of extant land plant species, whereas gymnosperms, their extant sister group, account for only a small fraction of this diversity. Explaining such heterogeneity in diversity is a central challenge in evolutionary biology. The land plant diversity throughout the Phanerozoic was likely shaped by a complex interplay of biotic interactions, environmental change, and intrinsic evolutionary innovations [17]. Among these potential drivers, abiotic factors are widely regarded as fundamental to long-term diversification dynamics and the wax-and-wane of major clades [18]. Land plants evolved against a backdrop of profound tectonic and climatic transformations throughout the Phanerozoic [6,19–21], including continental rearrangements and fragmentation [22,23], major fluctuations in global temperature [24], and mass extinction events [25]. These geological changes may affect diversification by altering habitat availability, geographic connectivity, and ecological opportunity [6,19–21]. Accordingly, the “geological events” scenario predicts that diversification rates are more strongly associated with abiotic changes than with biotic variables (Fig. 1A).

**Fig. 1.**
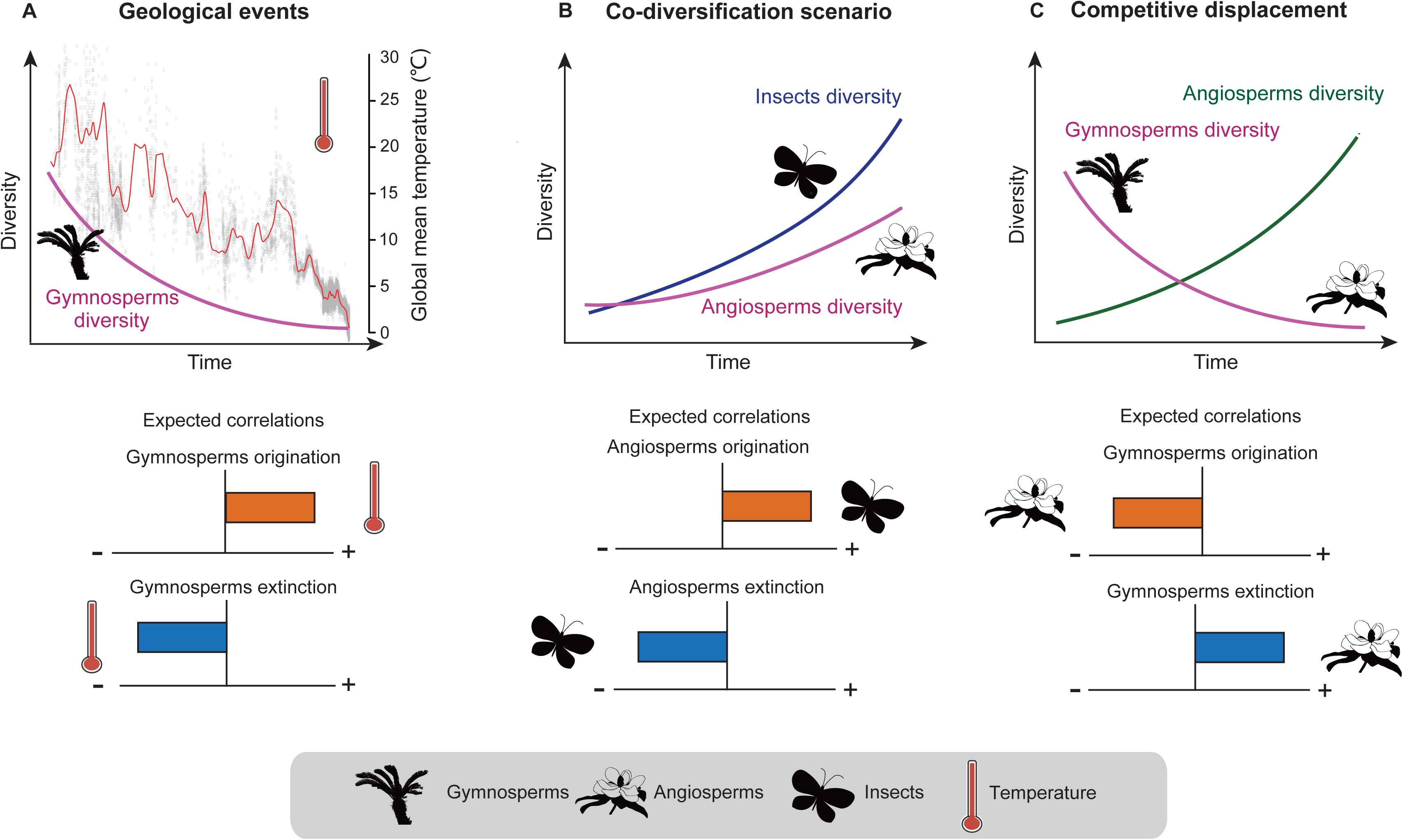
Main scenarios to explain the past diversity dynamics of major groups of land plants (exemplified by angiosperms or gymnosperms). The horizontal bars below each scenario summarize the predicted associations between the explanatory variables and plant origination (blue) or extinction (orange) rates. Bars extending to the right and left indicate positive and negative associations, respectively. A filled bar indicates a significant correlation parameter for a given variable [shrinkage weights (ω) > 0.5]. **(A)** Co-diversification scenario. The diversity and origination of angiosperms increasing synchronously to insect diversification. The expected result for this hypothesis is a significant positive correlation between angiosperms origination and the rise of insects. **(B)** Competitive scenario. A significant positive correlation is expected between the extinction of gymnosperms and the rise of angiosperm diversity, and a significant negative correlation between the origination of gymnosperms and the increase of angiosperm diversity. **(C)** Geological events scenario: the wax-and-wane of gymnosperms depends on the environmental forcing such as global temperature variations or continental fragmentation. The expectation for this hypothesis is that there no correlation between the gymnosperm origination and extinction and the rise of angiosperm diversity and insects. Silhouettes from http://phylopic.org/ (license at https://creativecommons.org/publicdomain/zero/1.0/).

In addition to environmental forcing, biotic interactions have also been proposed as important drivers of diversification. Under relatively stable environmental conditions, interactions among species may play an increasingly important role in shaping evolutionary trajectories [26,27]. One scenario is “co-diversification”, whereby reciprocal interactions between ecologically associated lineages promote diversification [28]. The extraordinary diversity of angiosperms and pollinating insects has often been interpreted as a classic example of co-diversification mediated by mutualistic interactions [29,30]. Under this co-diversification scenario, temporal changes in plant and insect diversification are expected to be positively associated. Alternatively, an increase in insect diversity could reduce the extinction rate of plants (Fig. 1B). However, fossil evidence suggested that the early diversification of angiosperms and insect lineages were temporally decoupled, because several insect groups that later became important angiosperm pollinators originated well before crown-group angiosperms [31]. These ancient insect lineages were initially associated with gymnosperms, while broader fossil evidence also documents diverse plant-insect interactions involving ferns and other early vascular plants [31–35]. Whether such interactions promoted the diversification of early land plant groups and subsequently shifted toward angiosperm-dominated associations remains unclear.

Diversification may also be shaped by ecological replacement among competing lineages. The “competitive displacement” scenario proposes that radiation of an emerging clade may be accompanied by diversity declines in previously dominant groups [36–38]. Under the competitive displacement scenario, increasing diversity in an emerging plant group is expected to be associated with reduced origination or increased extinction in previously dominant groups (Fig. 1C). Consistent with this scenario, the rise of major vascular plant groups has often coincided with the stabilization or decline of formerly dominant lineages [38]. Fossil assemblages from South China further document a gradual turnover of plant diversity across the Permian-Triassic transition [39], illustrating the potential importance of long-term ecological restructuring in shaping diversification patterns.

Despite extensive research into abiotic and biotic influences on diversification, their relative importance in shaping land plant diversity remains poorly understood. Resolving this question requires complementary evidence from the fossil record and molecular data. Fossils provide a direct, although incomplete, archive of past biodiversity, documenting the origins, radiations, declines, and extinctions of taxa through geological time [40,41]. Time-calibrated phylogenies of extant species provide a complementary perspective on lineage diversification, particularly where the fossil record is sparse. Concordant patterns recovered from these independent sources can strengthen interpretations of land plant evolutionary history [13,37]. Nevertheless, most previous studies have focused on particular clades, specific time intervals, or individual diversification correlates. Few have quantitatively assessed diversification across all major land plant groups using both fossil occurrences and molecular phylogenies. The continued expansion of paleontological and molecular datasets, together with recent developments in Bayesian birth-death modelling, now provides an opportunity to reconstruct land plant diversification and evaluate its potential drivers.

Here, we estimate the diversification dynamics of land plants through deep time using fossil record and molecular data. First, we analyze an updated time-calibrated phylogeny within a Bayesian birth-death framework to infer diversification patterns among extant land plant lineages. Second, we use PyRate to estimate temporal variation in origination, extinction, and net diversification rates from a comprehensive compilation of fossil occurrences. We then examine whether origination and extinction dynamics of land plants changed across major mass extinction events. Finally, we use a multivariate birth–death model to evaluate associations predicted by three non-mutually exclusive scenarios (geological events, co-diversification, competitive displacement). Together, these analyses provide new insights on the processes associated with the assembly of land plant diversity over more than 450 million years.

## Results and Discussion

### Diversification dynamics of land plants based on the time-calibrated phylogeny

Using an updated time-calibrated phylogeny comprising approximately 30,000 extant species, we estimated the diversification pattern across land plants with BAMM [42]. The mean net diversification rate across the phylogeny was estimated to be 0.061 events per lineage per million years (events/Myr). However, rates varied markedly among lineages, with multiple supported rate shifts occurring within species-rich clades (Fig. 2). Consistent with recent phylogenetic studies [12,15], our results recovered a pronounced increase in angiosperm diversification during the Cretaceous, accompanied by numerous rate shifts (Fig. 2). This time interval also contained extensive gene tree conflicts across the angiosperm phylogeny [15], which may represent an additional characteristic of the rapid succession of lineage divergences during the early angiosperm radiation.

**Fig. 2.**
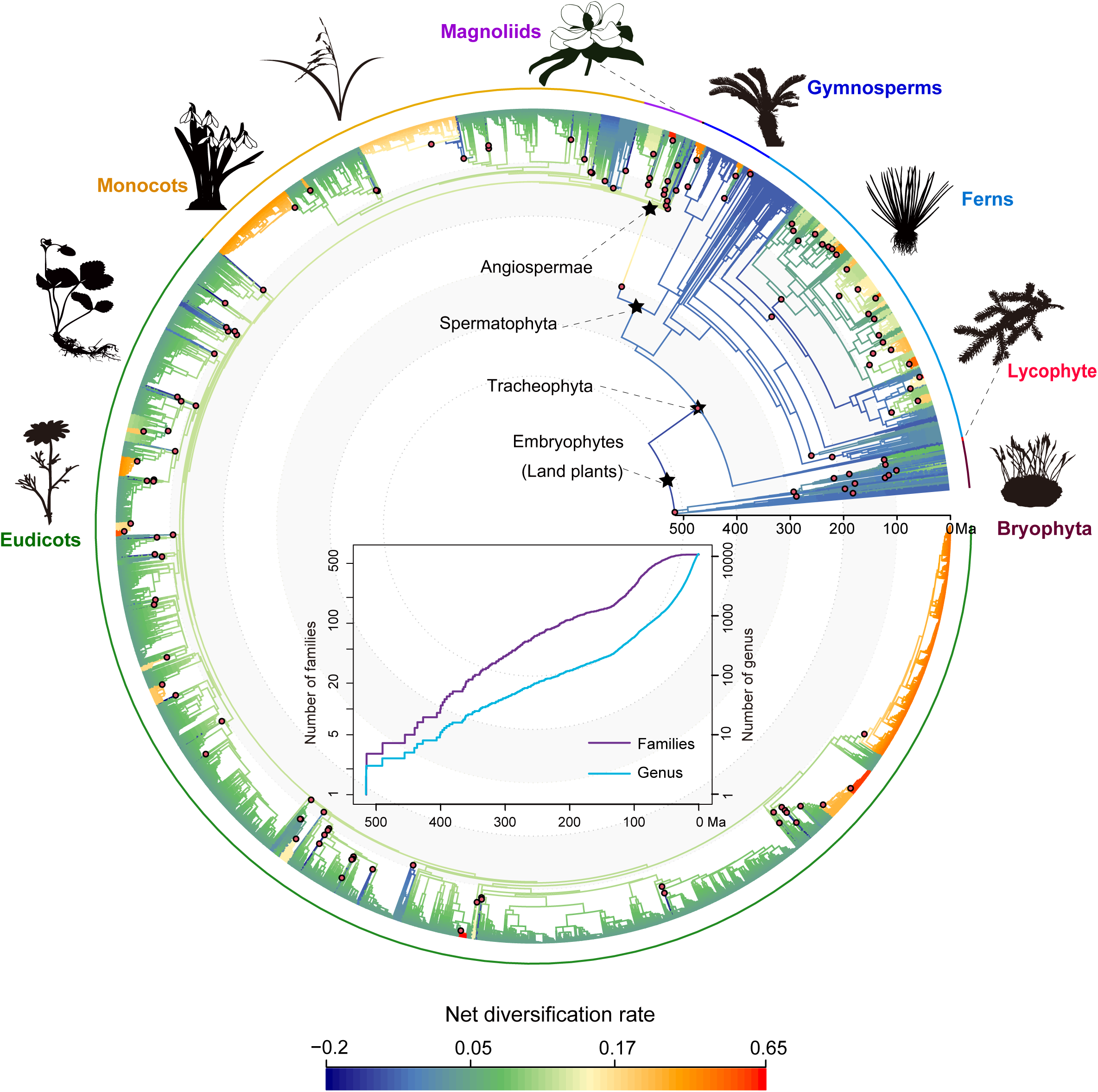
Temporal dynamics of land plant diversification. Maximum a posteriori probability shift configuration showed variation in net diversification rates across the phylogeny of land plants. The diversification shifts marked with circles and the origin of land plants marked with asterisk.

Elevated diversification rates were also recovered in several fern lineages, broadly coinciding with the rapid radiation of angiosperms during the Cretaceous (Fig. 2). Previous studies have proposed that the expansion structural complexity of angiosperm-dominated forests created new ecological opportunities for ferns, including shaded understory environments and epiphytic niches [43–46]. In particular, shade-dwelling ferns appear to have diversified more rapidly than sun-dwelling lineages following the expansion of angiosperm-dominated forests [45]. Genus and family level lineage-through-time plots likewise showed an increase in lineage accumulation beginning in the Cretaceous (Fig. 2). The temporal correspondence between angiosperm and fern diversification was therefore consistent with the ecological-opportunity hypothesis. By contrast, lycophytes and gymnosperms displayed lower and more temporally stable diversification rates. Bryophytes also exhibited relatively low diversification rates despite their substantial extant species richness (Fig. 2A). However, bryophytes were sparsely represented in our phylogeny, with considerably lower proportional coverage than most vascular plant groups. Their diversification-rate estimates should therefore be interpreted cautiously, because sparse and uneven sampling may reduce the power to detect rate variation even when incomplete sampling is incorporated into the model.

### Temporal dynamics of land plant diversification inferred from the fossil record

We further investigated the temporal variation in the rates of origination and extinction using the genus-level fossil occurrence and a Bayesian birth-death model implemented in PyRate accounting for taxonomic and temporal heterogeneous preservation rates. Model comparisons indicated that the fossil datasets for land plants as a whole and for each of the major groups best fit the time-variable preservation model.

We identified four major episodes of elevated diversification in land plants: the Middle Devonian (Eifelian) (∼393–388 Ma), the Carboniferous (Bashkirian) (∼323–315 Ma), the Lower Triassic (∼252–247 Ma), and the Paleocene (Thanetian)–Eocene (Ypresian) (∼59–48 Ma) (Fig. 3A). Origination rates of land plants exhibited four peaks at approximately 393 Ma, 323 Ma, 252 Ma, and 56 Ma, corresponding to the Early–Middle Devonian, Mississippian–Pennsylvanian, Permian–Triassic, and Paleocene–Eocene boundaries, respectively. Extinction rates reached their highest estimated values near 393 Ma and the Permian–Triassic boundary at about 252 Ma. Estimated extinction rates were generally lower during the Mesozoic than during the Paleozoic (Fig. 3A). Consistent with previously reported family-level patterns in vascular plants [47], genus-level richness declined markedly across the Permian–Triassic boundary, recovered rapidly during the Triassic, and remained relatively stable until the Early Cretaceous (Fig. 4A).

**Fig. 3.**
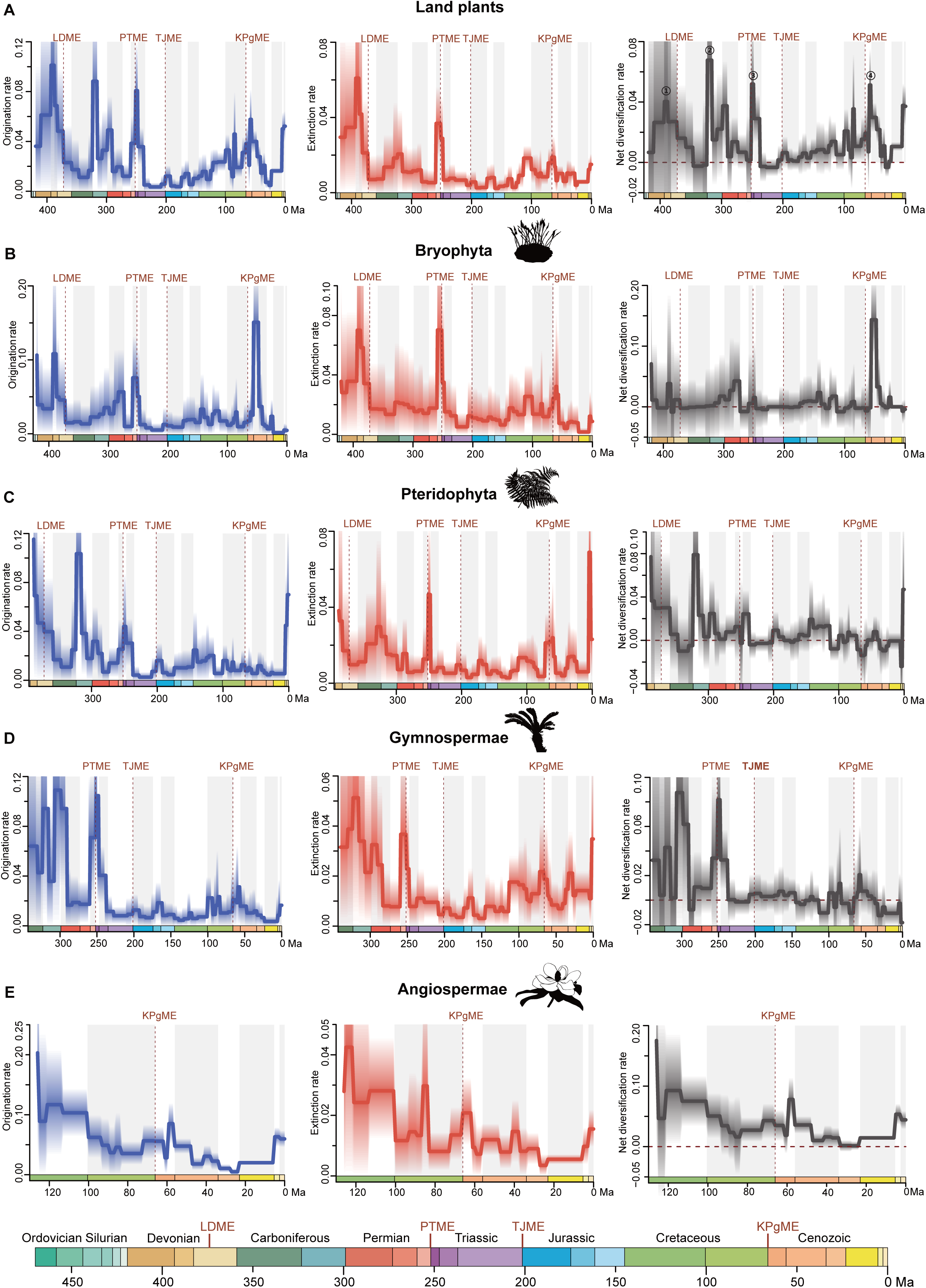
Genus-level diversification analysis of all land plants. Solid lines indicate mean posterior rates, whereas the shaded areas show 95% CI. Net diversification rates (grey) are defined as origination minus extinction. The dashed lines indicate the major mass extinction events widely recognized: the Late Devonian mass extinction (LDME), Permian–Triassic mass extinction (PTME), Triassic–Jurassic mass extinction (TJME) and Cretaceous–Paleogene mass extinction (KPgME). Ord: Ordovician, Sil: Silurian, Car: Carboniferous. Silhouettes from http://phylopic.org/ (license at https://creativecommons.org/publicdomain/zero/1.0/).

**Fig. 4.**
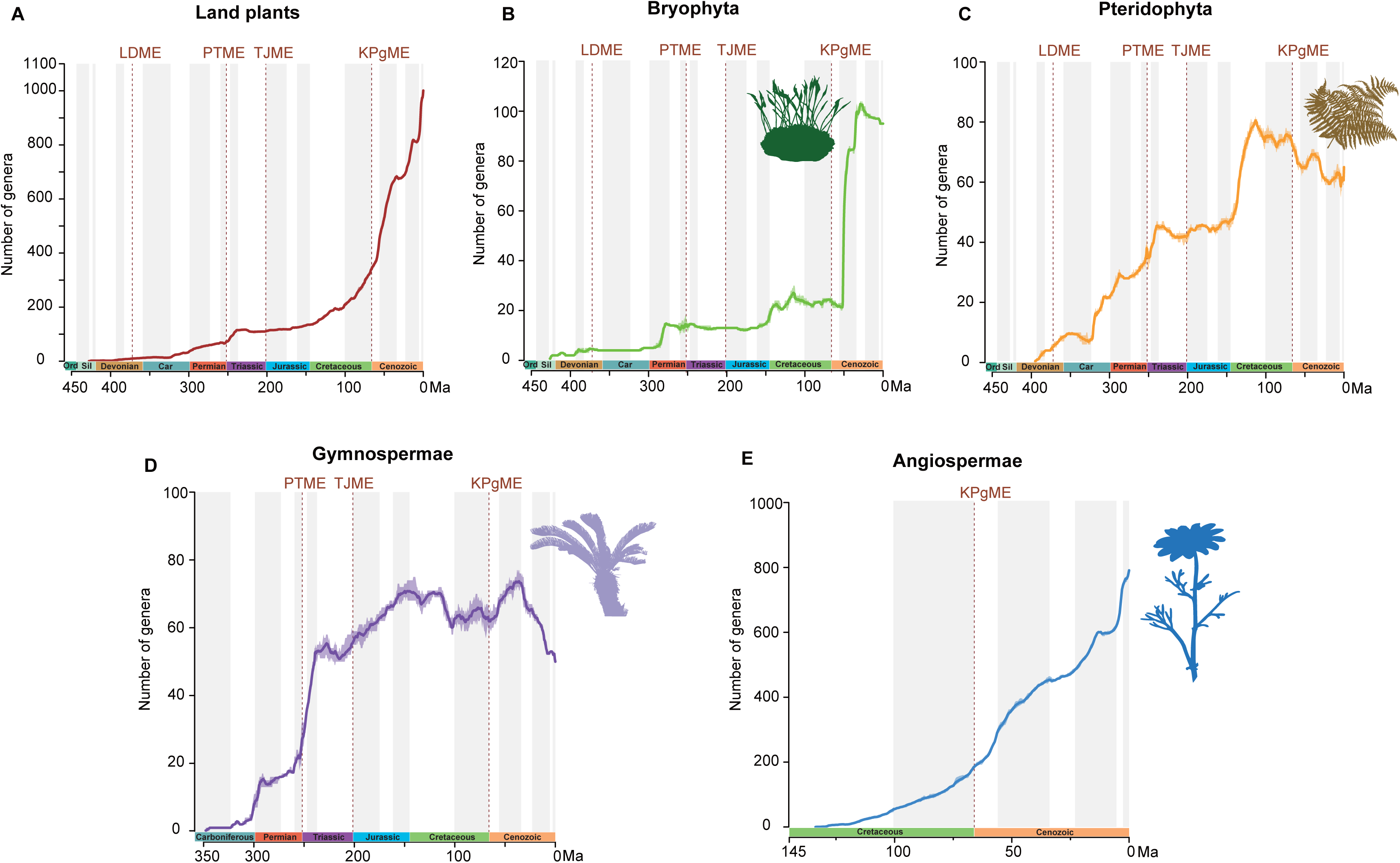
Accumulated diversification of land-plant major groups’ genera through geological time estimated using the software PyRate. Red dotted lines indicate the four mass extinctions. Ord: Ordovician, Sil: Silurian, Car: Carboniferous.

For bryophytes, PyRate analyses inferred a marked increase in net diversification rates following the K–Pg boundary (Fig. 3). Estimated genus-level richness began to increase rapidly at approximately 50 Ma and reached its maximum during the Oligocene (Fig. 4B), aligning with diversification patterns inferred from fossil-calibrated bryophyte phylogeny [48]. This post K–Pg increase of bryophytes coincided with substantial ecological and climatic restructuring. The development of structurally complex, angiosperm-dominated forests during the Early Eocene may have expanded shaded and humid habitats suitable for epiphytic bryophytes [49,50]. In addition, contemporary warming and reduced aridity may also promote both forest expansion and bryophyte diversification [51,52].

The fossil-based results differed partly from our phylogenetic estimates, which indicated relatively low overall diversification rates in bryophytes, although several lineages showed elevated rates during the Cretaceous (Fig. 2). These two approaches exhibited differences in both taxonomic scale and observation process. PyRate estimated genus-level rates while accounting for fossil preservation, whereas BAMM inferred diversification exclusively among sampled extant lineages. The sparse fossil record and limited sampling of bryophytes in our analyses also introduced uncertainties. The discrepancy should therefore be interpreted cautiously rather than as direct disagreement between the fossil and phylogenetic records.

Pteridophytes and gymnosperms exhibited three broad phases of diversification (Fig. 3C,D and Fig. 4C,D). Before the Early Triassic, both groups pronounced fluctuations in net diversification, including several intervals of elevated rates. From the Late Triassic to Late Jurassic, their net diversification rates fluctuated within a narrower range. From the Cretaceous onward, reduced origination and elevated extinction produced negative net diversification during several intervals. Nevertheless, genus richness recovered from the Late Paleocene into the Eocene, when origination again exceeded extinction. Complementing these fossil-based patterns, our BAMM analyses indicated that fern lineages experienced rapid diversification in the Cenozoic. Previous studies have suggested that increasingly complex forest understories and the expansion of epiphytic niches provided ecological opportunities for fern diversification [43,44,46].

Angiosperms were characterized by high origination and relatively low extinction rates during the Cretaceous (Fig. 3E). Both rates declined during the Cenozoic, but net diversification remained relatively high and positive across most intervals (Fig. 3E). Compared with other land plant groups, angiosperms maintained higher net diversification rates and did not experience major diversity declines during their radiation, with the notable exception of the Eocene-Oligocene transition (Fig. 3E). Persistently positive net diversification resulted in a substantial accumulation in angiosperm genus richness throughout the Cretaceous and Cenozoic (Fig. 4E), a trajectory broadly consistent with molecular phylogenetic evidence for elevated angiosperm diversification [15].

Moreover, we evaluated the sensitivity of these inferences by excluding singleton taxa, randomly subsampling 50% of the occurrence data, and combining 50% subsampling with singleton exclusion. All these sensitivity analyses recovered overall diversification trajectories largely consistent with those inferred from the complete dataset (Supplementary Figs. 1-4).

### Changes in land plant diversity during mass extinction events

We examined changes in origination, extinction, net diversification, and genus richness across four major mass extinction boundaries represented in the land plant fossil record. Following the Late Devonian mass extinction (LDME), net diversification declined in bryophytes but remained relatively stable in pteridophytes (Fig. 3B, C). More pronounced changes occurred across the Permian–Triassic mass extinction (PTME), the most severe biotic crisis of the Phanerozoic [53,54]. Extinction rates increased sharply in bryophytes, pteridophytes, and gymnosperms, but only pteridophytes underwent a marked transition to negative net diversification (Fig. 3B–D), consistent with previous fossil-based analyses [5]. Pteridophyte genus richness declined by approximately 11% across the boundary (Fig. 4C). Shortly thereafter, pteridophyte net diversification rebounded and reached a peak, resembling the rapid post-extinction recovery described as “flash recovery” in some marine groups, including ammonoids [55]. Bryophytes and gymnosperms had both high origination and extinction during the PTME. Gymnosperms subsequently experienced another increase in net diversification during the Early Triassic, followed by a diversity decline during the Late Triassic. Across the Triassic–Jurassic mass extinction (TJME), we detected no significant changes in extinction rates for bryophytes or gymnosperms, while pteridophytes experienced a small peak in net diversification.

The most widespread lineage response was associated with the Cretaceous– Paleogene mass extinction (KPgME), during which extinction rates increased to varying degrees across most major land plant groups (Fig. 3B– E). Genus-level richness declined by approximately 13% in bryophytes, 14% in pteridophytes, and 4% in gymnosperms, with gymnosperms reaching their lowest estimated richness during this interval (Fig. 4B–D). Angiosperm extinction rates also increased and their net diversification declined across the K– Pg boundary. Nevertheless, net diversification remained positive and rebounded to a peak within approximately 6 Ma (Fig. 3E). Consistent with Thompson and Ramírez-Barahona [14], our results suggested that angiosperms were less affected by the K–Pg mass extinction than the other major groups. This comparatively rapid recovery may partly reflect differences in life-history and functional traits among plant lineages [56]. Fossil leaf assemblages indicated that extinction across the K– Pg boundary was ecologically selective, disproportionately affecting taxa with slow-return leaf strategies while favoring deciduous angiosperms with faster-return strategies [56]. More generally, many angiosperms exhibited faster growth and resource acquisition than gymnosperms [57,58]. Although these mechanisms were not tested directly here and vary substantially among lineages, they may have increased the capacity of some angiosperms to tolerate post-impact environmental stress and recolonize disturbed habitats. The K–Pg mass extinction may have acted as an ecological filter, creating opportunities for the subsequent expansion and ecological reorganization of surviving angiosperm lineages [59].

Following the KPgME, major land plant lineages exhibited distinct pulses of diversification. Whole-genome duplication (WGD) events represented one possible contributor to post-extinction recovery. Phylogenomic studies have identified multiple WGD events across vascular plants whose estimated ages broadly overlap with major mass extinctions [60–63]. Polyploidy may enhance genetic redundancy and provide genetic material for functional innovation, potentially increasing evolutionary flexibility under rapidly changing environmental conditions [64]. This expanded genomic toolkit may have enabled surviving lineages to evolve novel morphological and physiological traits, thereby promoting post-extinction recovery and subsequent diversification.

### Biotic and abiotic correlates of pre-Cretaceous land plant diversification

We used a multivariate birth–death (MBD) model [65] to examine associations between land plant origination and extinction rates and a suite of biotic and abiotic predictors, which has been instrumental for testing hypotheses on clades’ diversification [37,66,67]. Biotic predictors included the reconstructed genus-level richness of the four major land plant groups and the diversity of insects and pollinators. Abiotic predictors included global temperature, arid regions, continental fragmentation, atmospheric carbon, marine sedimentary selenium (Se), and sediment flux. Notably, the MBD results represented macroevolutionary associations rather than evidence of causal effects.

During the Permian–Early Cretaceous interval, the MBD analyses indicated that the rise of gymnosperms was consistent with the decline extinction of bryophytes (ωµ = 0.97 and *G*µ = −9.53) (Fig. 5A, Supplementary Fig. 5 and Supplementary Table 1). This association could reflect ecological facilitation, differentiation in habitat use, or shared responses to environmental conditions. For example, to adapt to the environment dominated by gymnosperms, bryophytes may diverge into some particular ecological niches, such as the rock crevices, bases of tree trunks where gymnosperms have difficulty growing, and become the pioneer plant and trailblazer in ecological restoration of rocky desertification areas [68].

**Fig. 5.**
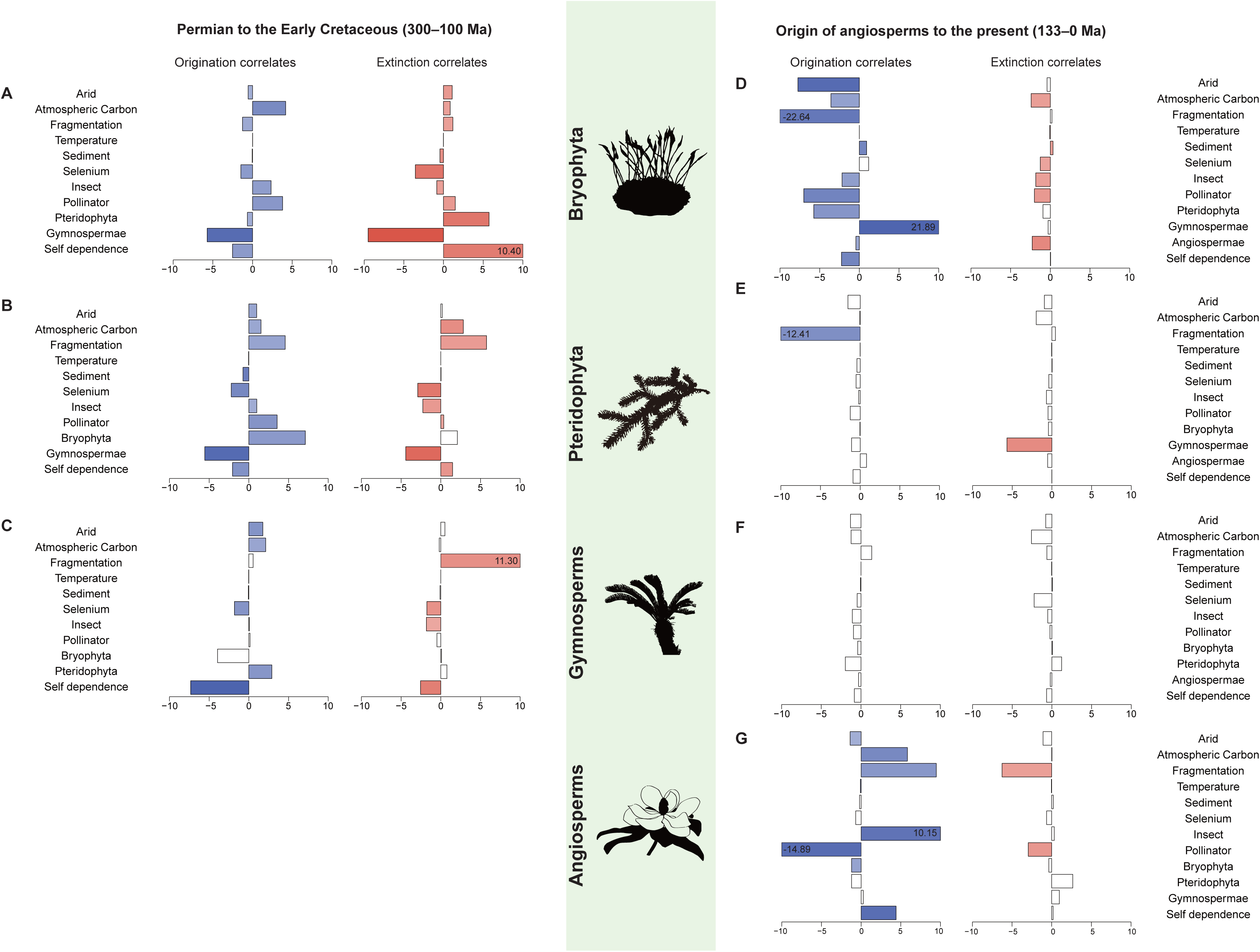
Estimated correlation parameters on origination and extinction rates of land plants divided by four major groups for two periods of time: Permian to the Early Cretaceous (300–100 Ma) and from the origin of angiosperms to the present (133–0 Ma). The drivers are global mean temperature (Temperature), Bryophyta relative diversity (Bryophyta), Pteridophyta relative diversity (Pteridophyta), Gymnosperm relative diversity (Gymnosperms), Angiosperm relative diversity (Angiosperms), Continental fragmentation (Fragmentation), Pollinator family richness (Pollinator), Insect family richness (Insect), and arid regions (Arid), Atmospheric Carbon, Selenium and Sediment. Bars extending to the right and left indicate positive and negative associations, respectively. A filled bar indicates a significant correlation parameter for a given variable [shrinkage weights (ω) > 0.5]. The x-axes have no unit scale for correlation parameters in the MBD analyses. If any of the dates is out of their corresponding scale, its value is displayed inside a box. In the MBD results, this temporal association should not be interpreted as evidence of direct causation.

For pteridophytes, we detected a significant negative correlation between gymnosperm diversity and origination rates (ωλ = 0.92 and *G*λ = −5.56) (Fig. 5B, Supplementary Fig. 6 and Supplementary Table 2). This pattern was consistent with competition displacement scenario, in which the expansion of an emerging group coincides with reduced establishment of new lineages in a formerly dominant group.

In the absence of ecological, physiological, or functional trait data, we cannot identify the mechanisms underlying this association or distinguish direct competition from shared responses to environmental change, which may also have independently constrained pteridophyte origination. Future integration of traits related to resource acquisition, water-use efficiency, reproductive strategy, and habitat preference could help evaluate whether functional advantages of gymnosperms contributed to this competition displacement pattern. Gymnosperm richness was also negatively associated with its own origination rate (ωλ = 0.96 and *G*λ = −7.34; Fig. 5C and Supplementary Table 3), a pattern consistent with intra-clade diversity dependence or ecological limits to diversification [69,70].

Insect diversity was associated with the diversification dynamics of both pteridophytes and gymnosperms during the Permian-Early Cretaceous. For pteridophytes, insect diversity was positively associated with origination and negatively associated with extinction rates (Fig. 5B and Supplementary Table 2). A similar pattern was recovered for gymnosperms, in which insect diversity was negatively associated with extinction rates (Fig. 5C and Supplementary Table 3). Fossil evidence provided support for specialized interactions between Mesozoic insects and gymnosperms [71,72]. For example, long-proboscid insects, used their elongated mouthparts to access pollination droplets exuded through narrow ovular micropyles, an association interpreted as potential pollination mutualism [73,74].

Several abiotic predictors, including atmospheric CO_2_, aridity, continental fragmentation, sediment flux and Se, were associated with diversification rates (Fig. 5A–C and Supplementary Tables 1–3). These predictors represented interconnected components of broader Earth-system change. Widespread aridity and variation in atmospheric CO_2_ during the Permian and Mesozoic may have altered the relative ecological performance of major plant groups, potentially favoring lineages with traits associated with water conservation [38,75,76]. During the Permian and Triassic, the extensive connected terrestrial with Pangaea and the development of large endorheic basins increased sediment-covered surface areas, potentially favoring the diversification of deep-rooting plants [6]. Furthermore, the MBD analyses identified negative associations between Se and both origination and extinction rates in bryophytes, pteridophytes, and gymnosperms (Fig. 5A–C). Se was derived from the weathering and erosion of continental rocks, reflecting changes in nutrient availability within continental systems [77]. However, its abundance was also influenced by transport processes, marine redox conditions, and sedimentary burial [78–80]. Because Se record covaries with other environmental variables, its associations with diversification should be interpreted as a composite Earth-system signal rather than as evidence for a direct biological effect of selenium.

### Transition toward angiosperm-dominated terrestrial flora from the Cretaceous

From the Cretaceous onward, major land plant groups underwent contrasting diversification trajectories. Bryophyte diversity has accumulated rapidly, especially from approximately 50 Ma to the present (Fig. 4B). In the MBD analyses, aridity was negatively associated with bryophyte origination rates (ωλ = 0.92 and *G*λ = −7.63), suggesting that reduced aridity may have facilitated bryophyte diversification during the Cenozoic (Fig. 5D, Supplementary Fig. 8 and Supplementary Table 1). For pteridophytes, we detected that the diversity of gymnosperms was negatively correlated with pteridophyte extinction rates (ωµ = 0.66 and *G*µ = −5.67; Fig. 5E, Supplementary Fig. 9 and Supplementary Table 2). This pattern indicated that declines in gymnosperm diversity coincided with elevated pteridophyte extinction. From the origin of angiosperms to the present, several factors were correlated with origination rates of angiosperms. Of the 12 tested variables, we found that the origination rate of angiosperms has strong positive association with self-diversity dependence (ωλ = 0.90 and *G*λ = 4.38; Fig. 5G, Supplementary Fig. 11 and Supplementary Table 4). This pattern was consistent with positive diversity dependence, potentially reflecting ecological opportunity, geographic expansion, or self-reinforcing increases in habitat and niche diversity [69,70].

The Late Cretaceous marked a major transition in terrestrial floras, accompanied by substantial climatic and tectonic restructurings. One important environmental shift was the demise of extensive latitudinal desert belts, which commenced in the Early Cretaceous and was largely complete by the Cenomanian [81]. This reduction in aridity may have increased connectivity among humid tropical and warm-temperate regions, creating opportunities for plant migration and diversification. Consistent with this scenario, angiosperm origination rates were positively associated with atmospheric CO_2_ and continental fragmentation, and negatively associated with aridity (Fig. 5G). These associations suggested that greenhouse conditions, changing paleogeography, and reduced arid barriers together may have contributed to the expansion of angiosperms [76,82,83]. Nevertheless, because continental fragmentation, atmospheric CO_2_, aridity, and biotic interactions covaried during this interval, these predictors should be interpreted as components of a broader Earth-system transition rather than as independent causal drivers. This environmental context may have helped angiosperms expand into newly available ecological space and contributed to the establishment of modern angiosperm-dominated terrestrial ecosystems [37]. Additionally, many globally distributed angiosperms have been inferred to diversify through niche shifts, land-bridge migration, and long-distance dispersal during the Cretaceous and Cenozoic [16,84,85].

We also detected that the transformation of insect communities since the Early Cretaceous coincided with the increasing ecological dominance of angiosperms (Fig. 5G). A positive correlation was identified between insect diversity and angiosperm origination (ωλ = 0.86 and *G*λ = 10.15), whereas corresponding associations with gymnosperms and pteridophytes were no longer supported (Supplementary Figs. 9–11 and Supplementary Tables 2–4). This shift was consistent with a reorganization of plant-insect associations toward angiosperm-dominated ecosystems. Fossil evidence suggested that some insect lineages later associated with flowering plants may previously have interacted with gymnosperms [81,86]. For example, mid-Cretaceous amber inclusions preserve beetles associated with abundant gymnosperm and early angiosperm pollen grains, supporting the possibility of transitional plant-insect associations during this interval [81,86]. A recent study further showed that angiosperms played a dual role that changed through time, mitigating insect extinction in the Cretaceous and promoting insect origination in the Cenozoic [87]. Together, our results suggested that the ecological expansion of angiosperms was accompanied by a shift from earlier insect associations with gymnosperms and pteridophytes toward increasingly angiosperm-centered patterns of co-diversification.

The recent study revealed episodic expansions of morphospace occupation during land plant evolution [88]. These episodes did not consistently coincide with periods of elevated origination rates or rapid increases in diversity recovered in our results. This discrepancy suggested that taxonomic diversification and morphological innovation may have been partially decoupled, such that increases in lineage diversity did not necessarily be accompanied by proportional expansions in morphospace occupation. Integrating fossil occurrence-based diversification estimates with trait-based disparity analyses will be necessary to evaluate this relationship.

Overall, the MBD results revealed patterns consistent with ecological interactions among major plant groups and responses to long-term environmental change. These results identified plausible correlates of land plant diversification rather than demonstrating the independent causal effects of individual drivers.

## Limitations

By conducting Bayesian analyses of a time-calibrated phylogeny and fossil occurrence data, we reconstructed the temporal diversification dynamics of land plants. Several limitations related to the datasets and analytical approaches should be considered. Simulation studies have demonstrated that PyRate can correctly estimate the past diversification dynamics even using incomplete taxon sampling and poor fossil record preservation [40,89]. When the incompleteness of the data reduced the power to recover the correct model of diversification, PyRate tends to be conservative, yielding near-uniform rates with wide credibility intervals rather than spurious rate shifts [89]. However, these results do not guarantee accurate inference in the presence of systematic taxonomic, temporal, or geographic biases.

It is notably that although we have assembled a large fossil occurrence dataset and analyzed an expanded time-calibrated phylogeny, the underrepresentation persists in some clades and geological intervals. Fossil occurrences of land plants are highly uneven across clades and geological intervals, and several lineages remain poorly represented. For example, despite being one of the largest and most rapidly diversifying angiosperm families, Orchidaceae have an exceptionally sparse fossil record. Similarly, gymnosperm macrofossils are substantially undersampled in parts of the Early Triassic, particularly the Induan, potentially producing an artifactual concentration of inferred extinction in the latest Permian [90]. In addition, bryophytes are the second-most diverse group of land plants, but their fossil record is considerably less complete than that of vascular plants. We supplemented fossil information from Tomescu et al. [91] into the bryophyte dataset, and conducted sensitivity analyses involving singleton exclusion and random subsampling. These analyses assessed the effects of singleton taxa and reduced sampling intensity but cannot fully correct systematic gaps in the fossil record. Consequently, diversification patterns inferred from fossil occurrence data alone may underestimate the major radiations within such groups.

Furthermore, our MBD results depended on the choice and availability of environmental and biological variables used as putative diversification drivers. We focused on 12 variables representing major biotic and environmental changes, but these predictors did not encompass all processes that may have influenced land plant diversification. Phylogenomic studies have associated major plant radiations, particularly in angiosperms, with whole-genome duplication and hybridization, which may generate genetic variation and facilitate functional innovation [92,93]. Moreover, many of the included predictors varied through time, making it difficult to partition their independent impacts. The estimated relationships should therefore be interpreted as associations with broad biotic and Earth-system changes rather than as causal effects of individual predictors.

A comprehensive understanding of land plant diversification requires complementary evidence from molecular phylogenies and the fossil record. Time-calibrated phylogenies reconstruct diversification among extant lineages and may obscure historical extinction and the evolutionary contributions of lineages without living descendants. Fossils directly document the occurrence of extinct taxa, allowing past origination and extinction dynamics to be estimated, but are affected by incomplete preservation, uneven sampling and variation in taxonomic practice. Therefore, these two sources of evidence cannot be regarded as mutually exclusive or entirely identical. Concordant patterns can strengthen macroevolutionary interpretations, whereas discrepancies may highlight the effects of extinction, preservation, taxonomic sampling, temporal resolution, or modeling assumptions. Therefore, combining these complementary sources can provide a comprehensive understanding of land plant diversification through deep time.

## Conclusion

Using an updated time-calibrated phylogeny and fossil occurrences, we reconstructed the deep-time diversification dynamics of land plants and evaluated the potential biotic and abiotic correlates. These complementary analyses revealed pronounced heterogeneity across lineages. Angiosperms exhibited the highest diversification rates, while several fern lineages underwent marked rate increases during the Cretaceous, broadly coinciding with the rapid radiation of angiosperms. Fossil-based analyses revealed four major episodes of elevated diversification in land plants. Of the four mass extinction events examined, the Cretaceous–Paleogene event was associated with the broadest cross-lineage response, whereas angiosperm net diversification remained positive. During the Permian–Early Cretaceous, increasing gymnosperm diversity was associated with reduced pteridophyte origination, a pattern consistent with competitive replacement. Insect diversity was associated with the diversification dynamics of pteridophytes and gymnosperms before the ecological expansion of angiosperms. From the origin time of angiosperms, the supported association shifted primarily toward a positive relationship between insect diversity and angiosperm origination, consistent with a reorganization of plant-insect associations toward angiosperm-dominated ecosystems. Diversification of major land plant groups were also associated with global environment changes, such as continental fragmentation, aridity, atmospheric CO_2_, sediment flux, and selenium. Our results suggested that modern land plant diversity was shaped by lineage-specific responses to the interacting effects of biotic turnover, mass extinction, ecological association, and long-term environmental change.

## Experimental Section

### Phylogenetic data

To construct a comprehensive land plant phylogeny, we used the crown node of Embryophyta from the One Thousand Plant Transcriptomes (1KP) phylogenetic framework [94] as a backbone constraint. We merged the bryophyte timetree including 531 species (533taxa) [95] and the vascular plant timetree comprising 76,225 species (“GBOTB_TS”) [96] by replacing these two clades in the 1KP tree. The final combined phylogenetic tree keeps the original divergence age estimates of the bryophyte and vascular plant tree, comprising 76,756 extant land plant species representing all major embryophyte lineages.

Bayesian diversification analyses on the complete land plant tree of 76,756 species were computationally challenging, we applied a genus-level subsampling strategy to the angiosperms, which accounted for the vast majority of species. For genera represented by more than five species, a maximum of five species was retained, whereas all species were retained for genera with five or fewer sampled species. The final phylogeny contained 30,278 species including 24,993 angiosperms, 825 gymnosperms, 3,893 ferns, 34 lycophytes, and 533 bryophytes (Supplementary Data 1), and was used for subsequent BAMM 2.5.0 [97] diversification analyses.

### Diversification analyses based on the molecular data

We analyzed the diversification dynamics of land plants using BAMM 2.5.0 [97]. To account for incomplete taxon sampling, we used a non-random incomplete taxon sampling correction and specified these sampling fractions by families. The family-specific sampling fractions were calculated by dividing the number of species represented in the time-calibrated phylogeny by the total number of accepted species recorded in World Flora Online (http://www.worldfloraonline.org/). The species richness and sampling fraction used for each family are provided in Supplementary Data 1.

We assigned appropriate prior values of the evolutionary rate parameters using the *setBAMMpriors* function in the R package BAMMtools [42]. The BAMM analysis was run for 10 million generations at a temperature increment parameter of 0.01, and event data were sampled every 5,000 generations. We discarded the first 25% of generations as burn-in and examined the effective sample size of the log-likelihood and the number of shift events for convergence with the R-package CODA [98]. The maximum *a posteriori* probability (MAP) shift configuration and rates through time analyses were inferred with BAMMtools.

### Fossil occurrences

To explore the long-term patterns of land plant diversification dynamics using fossil occurrences, we firstly classified the land plants into four main groups: Angiospermae, Gymnospermae, Pteridophyta, and Bryophyta, and retrieved genus-level fossil occurrences from the *Paleobiology Database* (PBDB; https://paleobiodb.org/#, last accessed in October 2024) for each group. To ensure consistency across datasets, all fossil occurrences were subjected to the same taxonomic cleaning and validation procedure regardless of plant group. Uncertain records (primary generic resolution noted by *cf., aff., ?, informal, ex. gr., “”*) with identical primary and accepted generic names were discarded, while those with different names (i.e., the species was reassigned to another genus) were retained.

Genus names were then standardized through manual taxonomic verification using multiple authoritative taxonomic resources, including *WFO Plant List* (https://wfoplantlist.org/), *Mindat*(https://zh.mindat.org/), *IRMNG* (https://www.irmng.org/) and *Taxonomicon* (http://taxonomicon.taxonomy.nl/), supplemented by relevant taxonomic literature, when necessary, supplemented by relevant taxonomic literature. Synonyms, outdated combinations, spelling variants, and other erroneous or doubtful records were cleaned and reconciled to a single accepted genus name. The same taxonomic framework and validation criteria were applied consistently across all fossil datasets. This procedure minimized inconsistencies arising from different taxonomic treatments among databases and ensured comparability across major land plant groups.

Absolute ages (maximum and minimum ages) of the occurrences were updated based on ages in the Geological Time Scale 2023/09 [99]. Occurrences with a high temporal uncertainty (>10 Ma), but not from an international stage (e.g., Aptian) were removed from the dataset. Additionally, we further supplemented the bryophyte dataset based on the fossil records provided by Tomescu et al. [91]. Our final dataset encompasses 39,857 fossil occurrences, which represent 1,629 land plant genera. The four-group fossil datasets contain: 1,036 fossil occurrences and 157 genera for bryophytes, 9,370 fossil occurrences and 190 genera for pteridophytes, 9,058 fossil occurrences and 194 genera for gymnosperms, and 20,393 fossil occurrences and 1,088 genera for angiosperms (Supplementary Data 2).

### Estimating origination and extinction rates based on the fossil data

We inferred the temporal dynamics of the times of origination (*Ts*) and extinction (*Te*) of each genus, the origination (λ) and extinction (*µ*) rates, and the preservation process (*q*) based on the fossil dataset using a birth–death model with constrained shifts (BDCS) [5,89] in a Bayesian framework implemented in PyRate 3 [40,41]. The robustness of PyRate has been thoroughly evaluated [5,40,89,100], overcoming a range of potential biases such as violations of sampling assumptions, variations in retention rates, and incomplete taxon sampling. It has been shown to correctly recover origination and extinction dynamics in the face of sudden rate changes and mass extinctions. We first used the *-PPmodeltest* function to assess three preservation models for land plants and each major lineage: 1) a homogeneous Poisson process (HPP), which assumes the preservation rate is constant through time; 2) a non-homogeneous Poisson process (NHPP) model, which assumes preservation rates change during the lifespan of each lineage following a bell-shaped distribution; and 3) a time-variable Poisson process (TPP) model that permits preservation rates are constant within a predefined time frame, but can vary across time frames. To further account for heterogeneity in preservation among taxa, we incorporated a Gamma model that allows preservation rates to differ across lineages. To avoid over-parameterization when fixing a prior on the number of rate shifts, we followed Silvestro et al. [100] and Condamine et al. [37] in assuming that the rates of origination and extinction are part of two families of parameters that follow a common prior distribution, with the parameters estimated from the data using hyper-priors. Specifically, we assigned a Gamma hyper-prior distribution with shape=1.5 and rate=0, allowing the MCMC to flexibly sample the origination and extinction rates while accounting for heterogeneity among lineages. To account for fossil ages, we generated 10 replicate datasets for each plant group. In each replicate, the age of fossil occurrence was randomly sampled from its documented minimum and maximum stratigraphic age bounds. We analyzed each replicate independently and combined the posterior samples of origination and extinction rates across replicates to generate rates-through-time plots. We inferred 10 posterior estimates of *Ts* and *Te* for all taxa and produced lineage through time plots for land plants and four major groups. All analyses were run for 500 million MCMC generations and sampled every 100,000 generations with time bins of geological epochs. After removing the first 10% of the MCMC sampling, we checked that the effective sample sizes (ESS) were greater than 200 for all parameters in each group by examining the MCMC log files in Tracer 1.7.1 [101].

To evaluate the robustness of our results to potential sampling bias in the fossil record, we conducted three sensitivity analyses. First, based on the main analyses, we reran all analyses using the *-singleton 1* option in PyRate to exclude genera represented by only a single fossil occurrence (singleton genera), thereby reducing the potential influence of sparsely sampled taxa. Second, we randomly subsampled 50% of the fossil occurrences within each geological time interval for the four major plant groups to assess the effect of reduced sampling intensity. The resulting dataset comprised 20,146 fossil occurrences representing 1,360 land plant genera, including 547 fossil occurrences and 113 genera for bryophytes, 4,796 fossil occurrences and 172 genera for pteridophytes, 4,582 fossil occurrences and 175 genera for gymnosperms, and 10,221 fossil occurrences and 900 genera for angiosperms. Using this subsampled dataset, we repeated the same analytical pipeline as in the main analyses. Specifically, we inferred 10 posterior estimates of origination (*Ts*) and extinction (*Te*) times for land plants and each of the four major groups based on these estimates. All analyses were run for 500 million MCMC generations, sampling every 100,000 generations with geological epochs used as time bins. After discarding the first 10% of samples as burn-in, convergence was assessed in Tracer v1.7.1 by confirming that the effective sample size (ESS) exceeded 200 for all estimated parameters [101]. Finally, to evaluate the combined effects of reduced sampling intensity and singleton taxa, we repeated the analyses on the 50% subsampled dataset after excluding singleton genera using the *-singleton 1* option in PyRate. The same analytical procedures as described above were applied to estimate *Ts* and *Te* for land plants and each major group.

### Analyzing the impact of abiotic and biotic factors on diversification

Since the origin of land plants, there have been four mass extinction events that occurred during geological periods. Based on the fossil occurrences, we mapped the four mass extinction events recorded in geological history onto the PyRate analysis of diversification rates to infer associations with origination and extinction rates. We further used the Multivariate Birth-Death model (MBD) implemented in PyRate to assess to what extent biotic and abiotic factors can explain temporal variation in origination and extinction rates.

Biotic predictors included the change in angiosperm, gymnosperm, spore-bearing plant and bryophyte diversities, insect and pollinator diversities. For each of the plant groups, we used the -*ltt* function of PyRate to estimate changes in diversity through time. The diversity estimates for each group were averaged across the retained MCMC samples, and the resulting mean trajectory was included as a separate predictor in the MBD analyses (see studies [5,20,65,67]). Relative diversity trajectories for insects and pollinators were obtained from previous family-level diversification estimates [87]. Abiotic predictors included temporal fluctuations of temperature [87], fragmentation of land masses [23], aridity [65], atmospheric CO_2_ [102], sediment flux [6], and selenium (Se) concentration in marine sediments [77]. The data source, temporal coverage and temporal resolution were summarized in Supplementary Tables 5-6.

The continental fragmentation index was obtained from [23] and was derived from palaeogeographic reconstructions for 1-million-year time intervals. The index approaches 0 when continental land masses are maximally aggregated and approaches 1 when all plates are disconnected. Marine Se was derived from the weathering and erosion of continental rocks [77], reflecting changes in nutrient availability within continental systems. Sediment flux was obtained from the Phanerozoic global paleo-landscape evolution dataset generated using the global Scalable Paleo Landscape Evolution (goSPL) model [6]. The dataset provides spatially explicit reconstructions of sediment flux across global terrestrial areas at a spatial resolution of approximately 5 km for each geological time slice. As the MBD model requires environmental variables to be provided as continuous time series, the spatial raster data were transformed into a single value for each time slice. Specifically, we calculated the global total sediment flux within each time slice, representing the overall intensity of global sedimentary activity. Enhanced sediment flux generally signifies intensified pedogenesis and elevated soil fertility, which could increase the carrying capacity of terrestrial ecosystems (Salles et al. 2023a). The resulting values were then used as an environmental predictor in the MBD analyses to evaluate the relationship between changes in global sedimentary dynamics and diversification rates.

In the MBD model, the estimation of origination and extinction rates is a function of multiple time-continuous variables. A MCMC algorithm jointly infers the baseline origination (λ0) and extinction (*µ*0) rates and correlation parameters (*G*λ and *Gµ*) with each variable. A positive *G* indicates that the factor is positively correlated with the rate and *vice versa*. We used exponential functions linking the temporal variation of birth-death rates with changes in these factors and employed a horseshoe prior algorithm to reduce the risk of over-parameterization [65]. We regarded an association as supported when the shrinkage weight (ω) of the corresponding correlation parameter exceeded 0.5. We ran the MBD model for 100 million generations and sampled every 5,000 generations. To consider the uncertainties in taxon origination and extinction, 10 replications were analysed separately for four groups of land plants (the *Ts* and *Te* for each group estimated by PyRate were used as input, avoiding re-modelling the complex preservation process). After excluding the first 10% of the samples, we combined the ten log files generated for each group. Convergence was evaluated using Tracer and parameters were considered convergent if their ESS were greater than 200. Notably, these parameters describe temporal associations and do not by themselves establish a causal effect.

### Statistical Analysis

Detailed procedures for data processing and model implementation are described in the preceding subsections. Briefly, the fossil-based analyses included 39,857 occurrences representing 1,629 genera, and the phylogenetic analyses included 30,278 extant species. Ten fossil-age replicate datasets were generated for each plant group. The numbers of occurrences and genera included in each group, together with the sources and processing of all predictor variables, are provided in Supplementary Tables 1-4.

Diversification-rate trajectories are presented as posterior summaries with 95% credible intervals where applicable, and BAMM rate shifts are summarized using the maximum a posteriori configuration. For MBD analyses, a predictor was considered statistically supported when its shrinkage weight (ω) exceeded 0.5. A positive G implies that the factor correlates positively with the rate and *vice versa*. Analyses were conducted using the R software, PyRate, BAMM, BAMMtools, CODA and Tracer.

## Acknowledgments

This work was supported by the National Natural Science Foundation of China (32370228 and W2511024) (B. Zhong); the Natural Science Foundation of Jiangsu Province (BK20250004) (B. Zhong); the Collaborative Innovation Center for Modern Crop Production co-sponsored by Province and Ministry (B. Z.); the Priority Academic Program Development of Jiangsu Higher Education Institutions (PAPD) (B. Z.); the European Research Council (ERC) under the European Union’s Horizon 2020 research and innovation programme (project GAIA, agreement no. 851188) (F.L.C); the “Investissements d’Avenir” program managed by the Agence Nationale de la Recherche (CEBA, ref. ANR-10-LABX-25-01) (F.L.C).

## Additional Information

Supplementary Information including Supplementary Figures 1–11, Supplementary Tables 1–6 and Supplementary Data 1–2 are available for this study. Time-calibrated phylogeny, fossil occurrences and diversification rate analyses have been deposited in Figshare: https://figshare.com/s/53a477d7cc5445888d4a.

