## Supplementary Figures 1-11 and Supplementary Tables 1-6 for "Bayesian inference of global land plant diversification dynamics across the Phanerozoic": Supplementary Information.pdf

This PDF file includes:

Supplementary Figures 1–11
Supplementary Tables 1–6

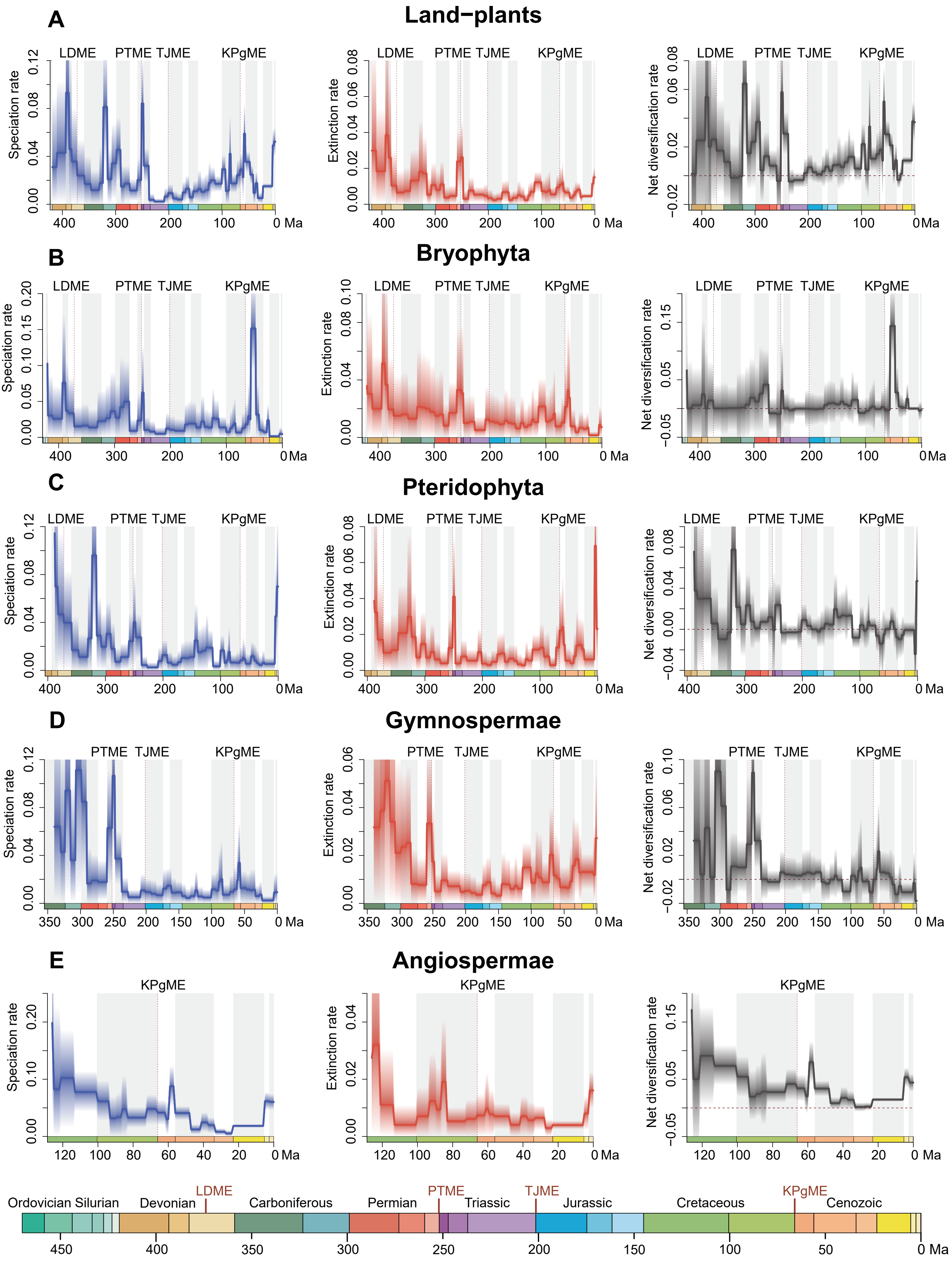

Supplementary Fig.1 Genus-level diversification analysis of all land plants without singletons. Solid lines indicate mean posterior rates, whereas the shaded areas show 95% CI. Net diversification rates (grey) are defined as origination minus extinction. The dashed lines indicate the major mass extinction events widely recognized: the Late Devonian mass extinction (LDME), Permian–Triassic mass extinction (PTME),Triassic–Jurassic mass extinction (TJME) and Cretaceous–Paleogene mass extinction (KPgME). Ord: Ordovician, Sil: Silurian, Car: Carboniferous.

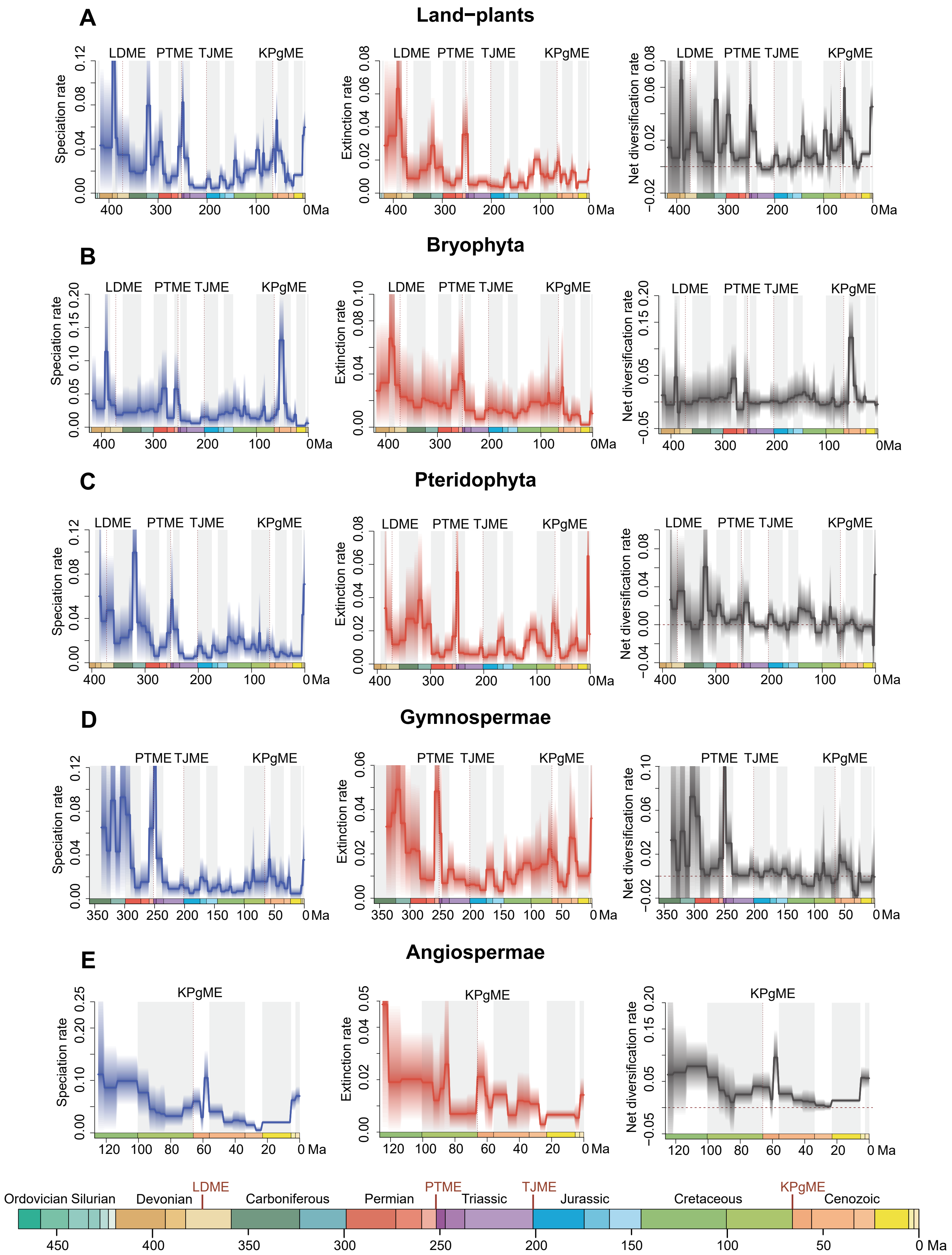

Supplementary Fig.2 Genus-level diversification analysis of a 50% sample of land plants. Solid lines indicate mean posterior rates, whereas the shaded areas show 95% CI. Net diversification rates (grey) are defined as origination minus extinction. The dashed lines indicate the major mass extinction events widely recognized: the Late Devonian mass extinction (LDME), Permian–Triassic mass extinction (PTME),Triassic–Jurassic mass extinction (TJME) and Cretaceous–Paleogene mass extinction (KPgME). Ord: Ordovician, Sil: Silurian, Car: Carboniferous.

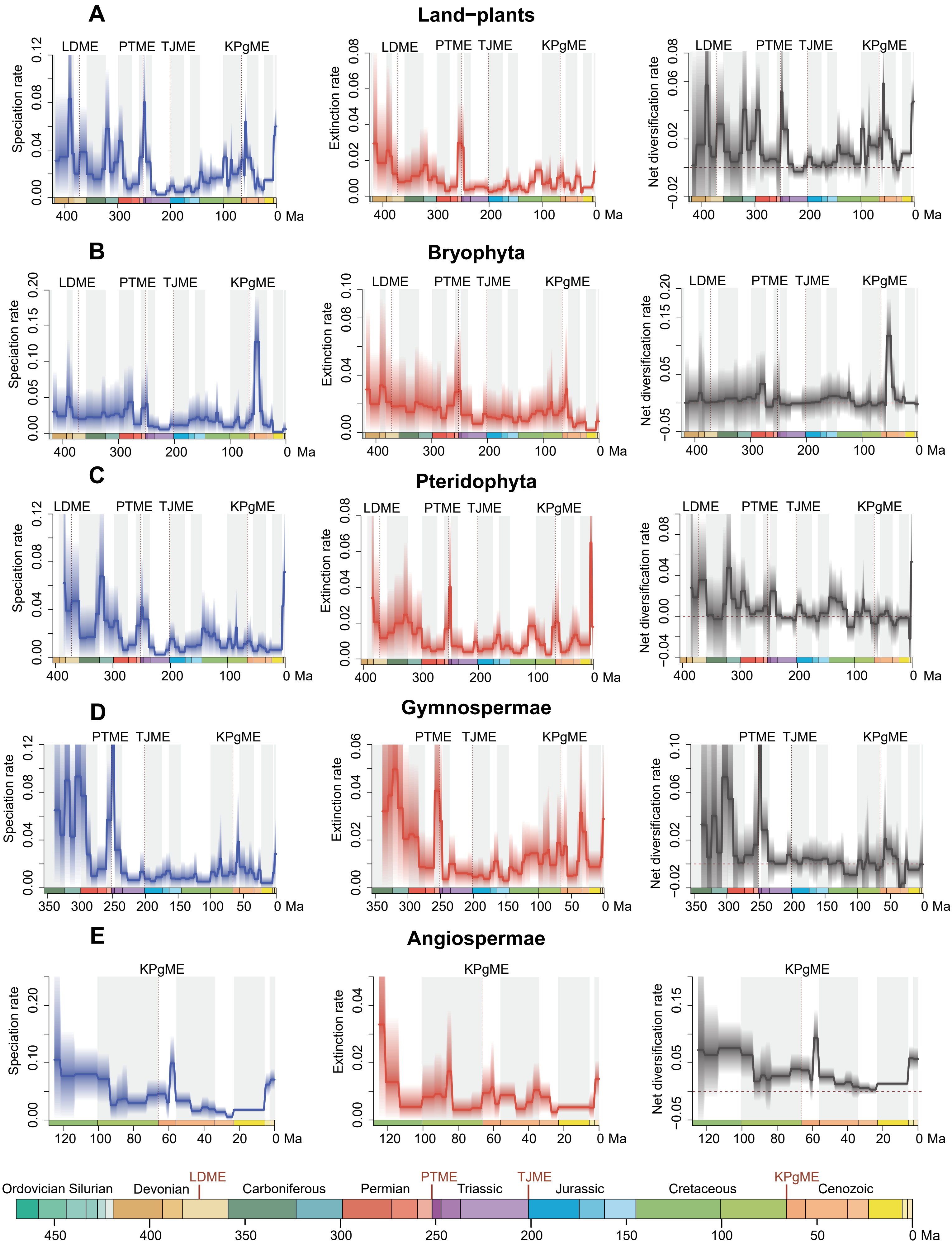

Supplementary Fig.3 Genus-level diversification analysis of a 50% sample of land plants without singletons. Solid lines indicate mean posterior rates, whereas the shaded areas show 95% CI. Net diversification rates (grey) are defined as origination minus extinction. The dashed lines indicate the major mass extinction events widely recognized: the Late Devonian mass extinction (LDME), Permian–Triassic mass extinction (PTME),Triassic–Jurassic mass extinction (TJME) and Cretaceous–Paleogene mass extinction (KPgME). Ord: Ordovician, Sil: Silurian, Car: Carboniferous.

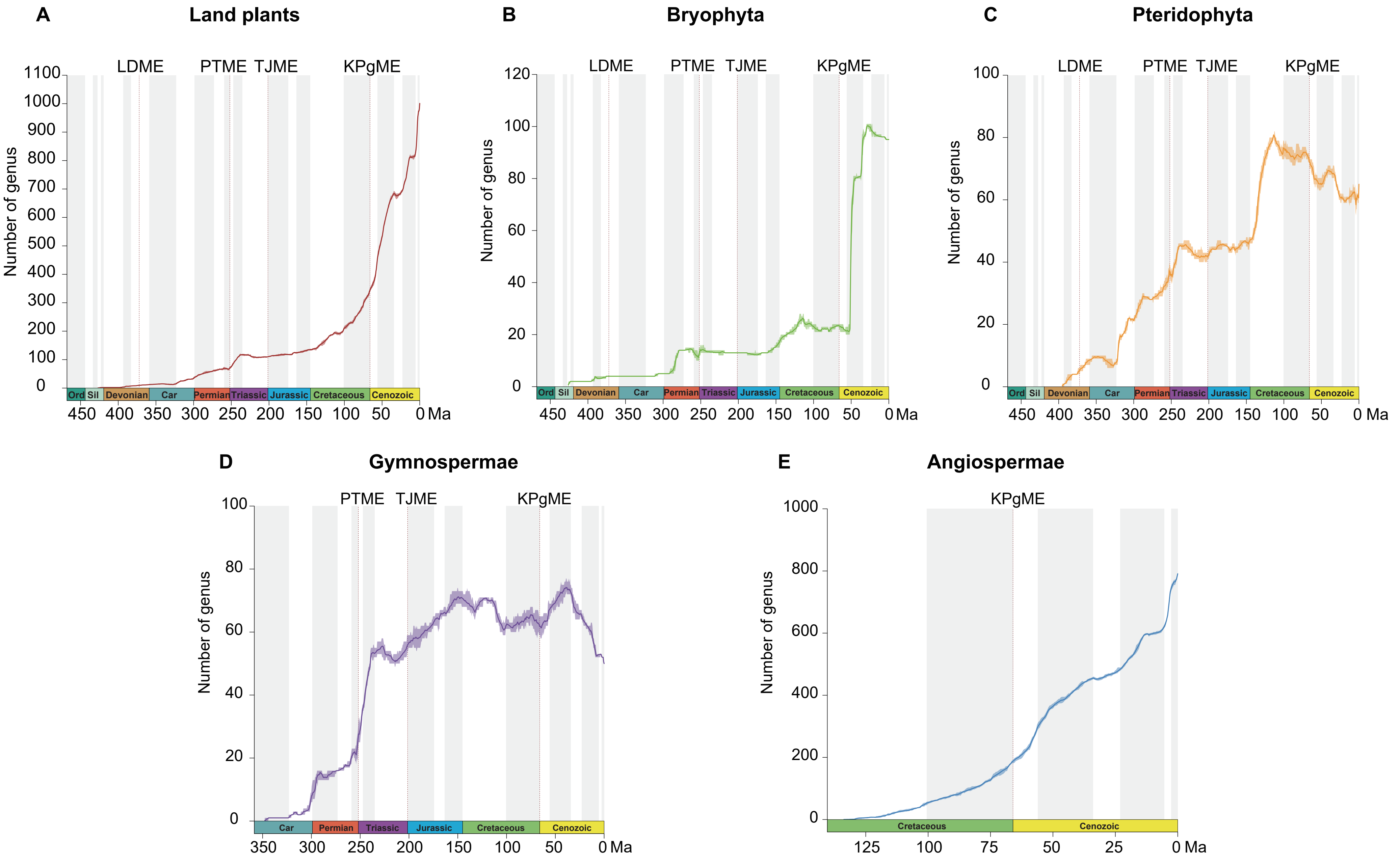

Supplementary Fig.4 Accumulated diversification of land-plant major groups genera without singletons through geological time estimated using the software PyRate. Red dotted lines indicate the four mass extinctions. Ord: Ordovician, Sil: Silurian, Car: Carboniferous.

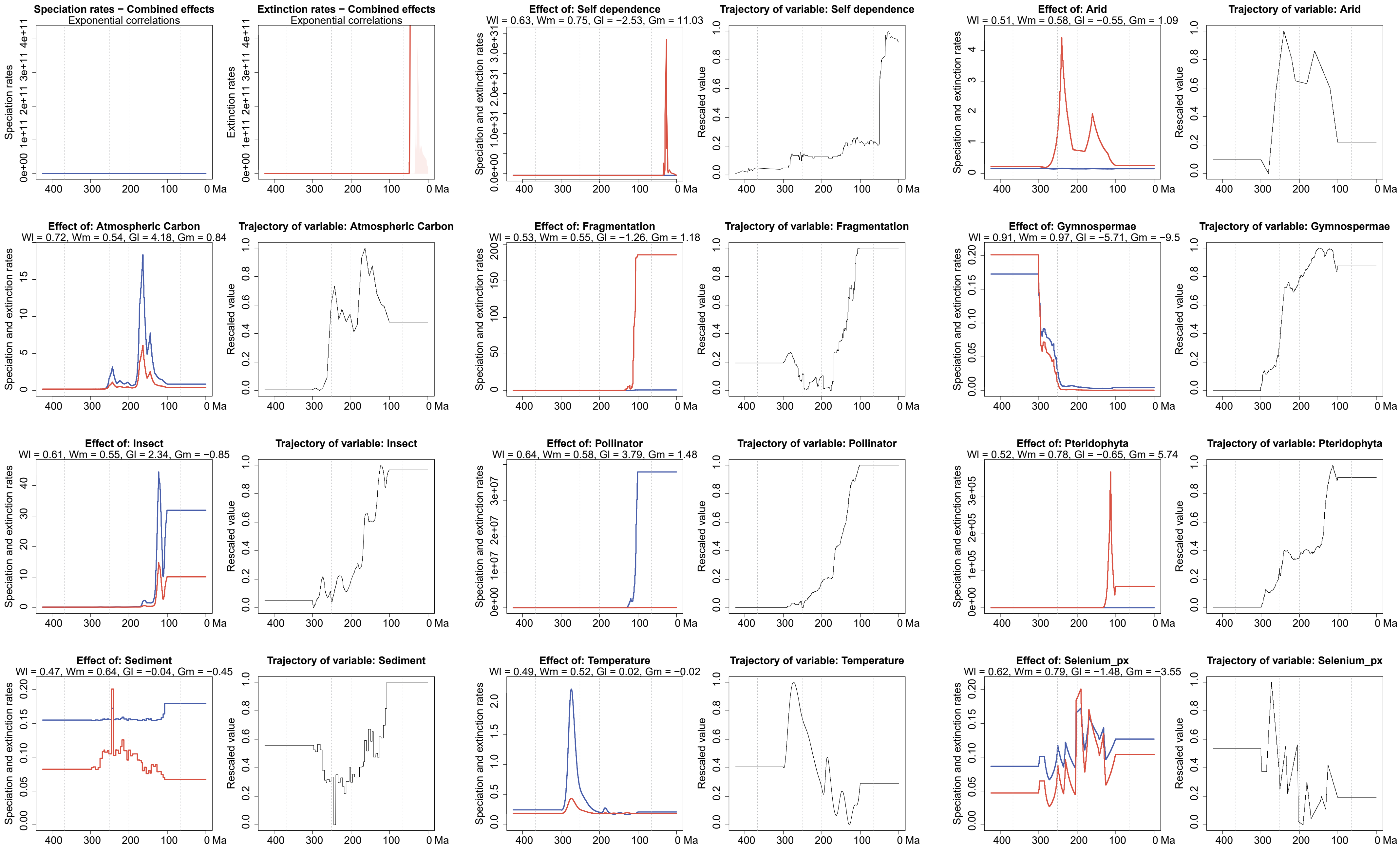

Supplementary Fig.5 The shrinkage weights (Wl and Wm for speciation and extinction, respectively) of Bryophyta during the time period from 300 to 100 Ma for all predictors are computed using MBD model. The shrinkage weights quantify the statistical support for each correlation factor. The correlation parameters are indicated by Gl and Gm.

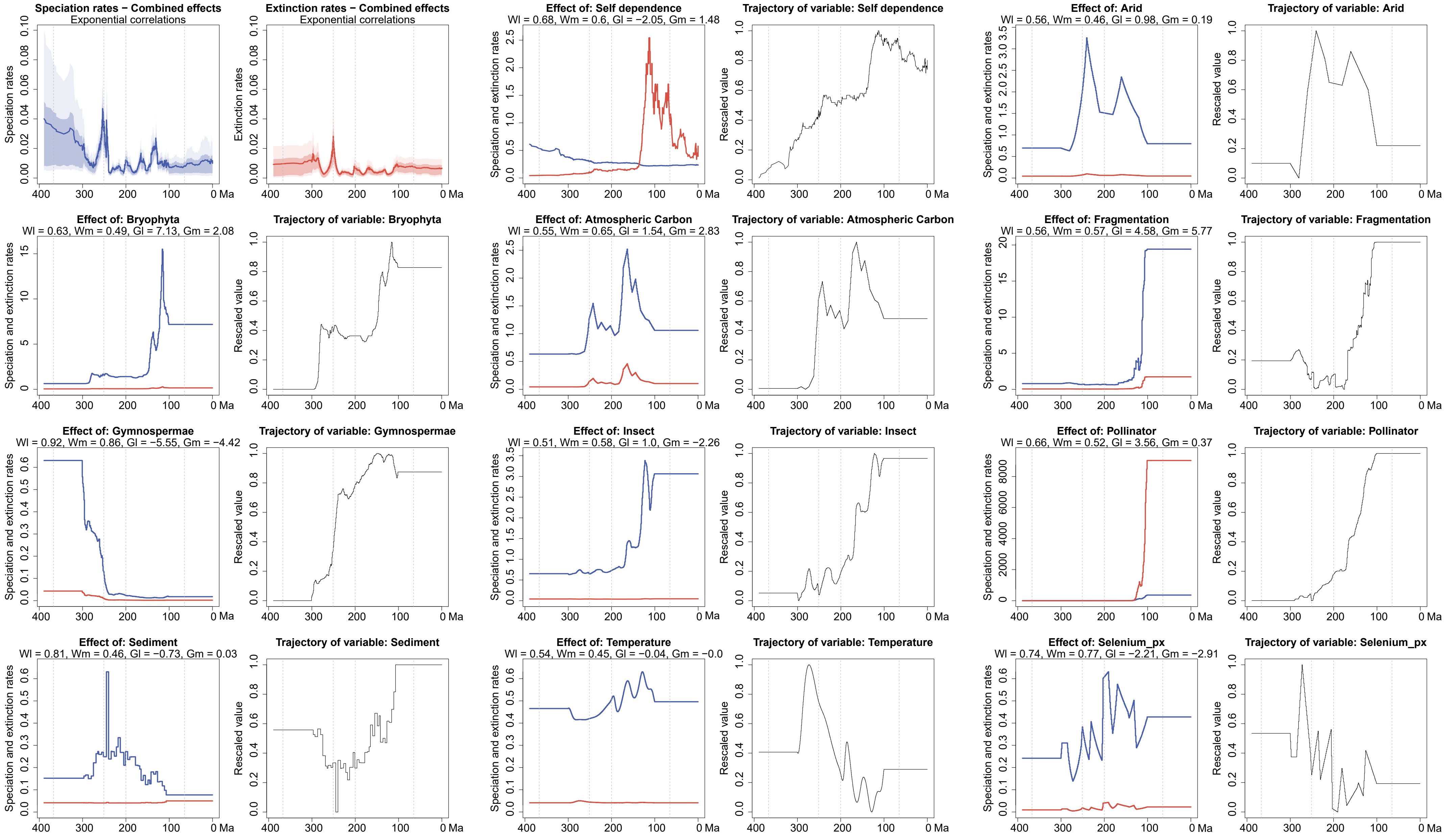

Supplementary Fig.6 The shrinkage weights (Wl and Wm for speciation and extinction, respectively) of Pteridophyta during the time period from 300 to 100 Ma for all predictors are computed using MBD model.The shrinkage weights quantify the statistical support for each correlation factor. The correlation parameters are indicated by Gl and Gm.

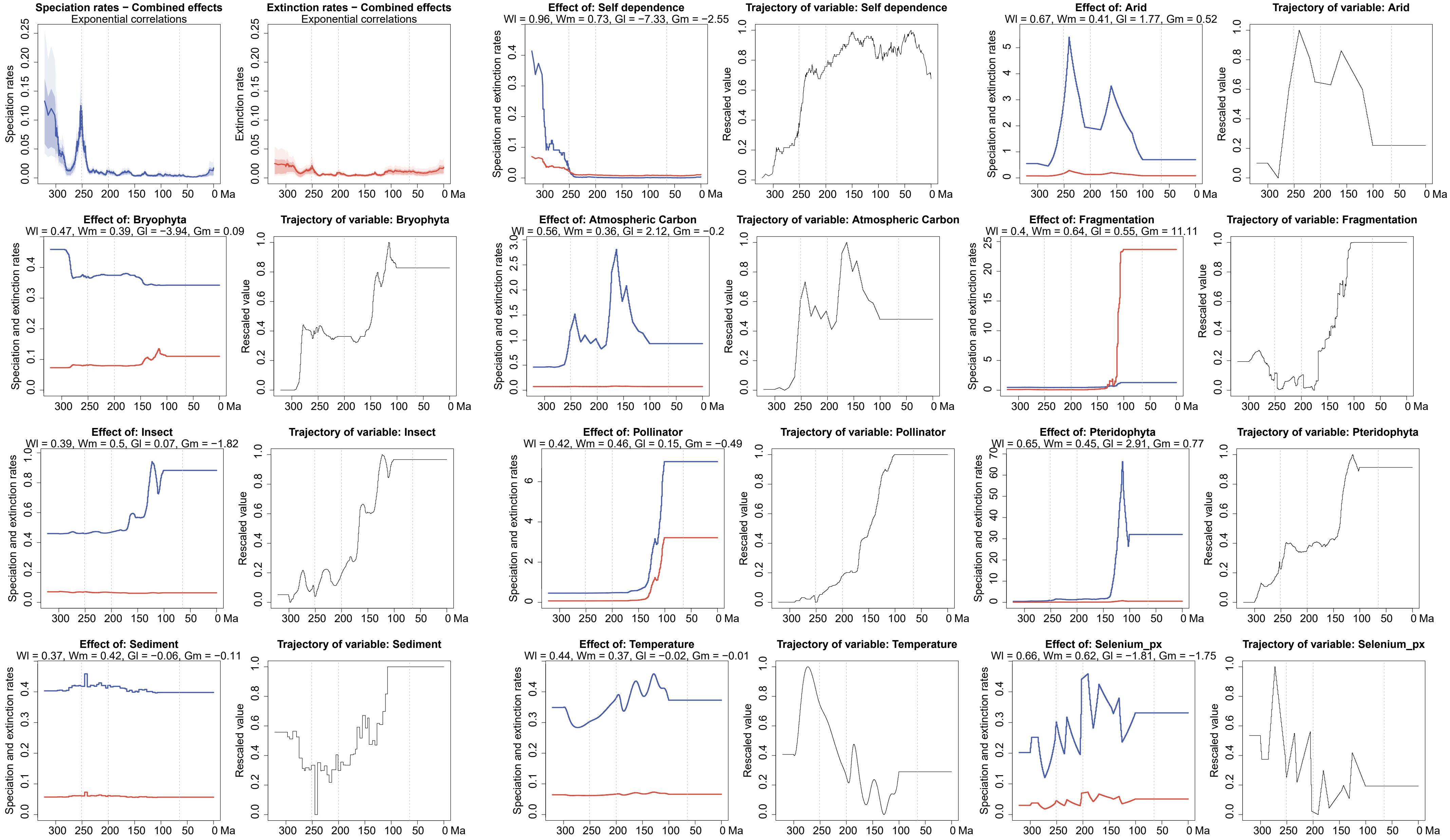

Supplementary Fig.7 The shrinkage weights (Wl and Wm for speciation and extinction, respectively) of Gymnospermae during the time period from 300 to 100 Ma for all predictors are computed using MBD model.The shrinkage weights quantify the statistical support for each correlation factor. The correlation parameters are indicated by Gl and Gm.

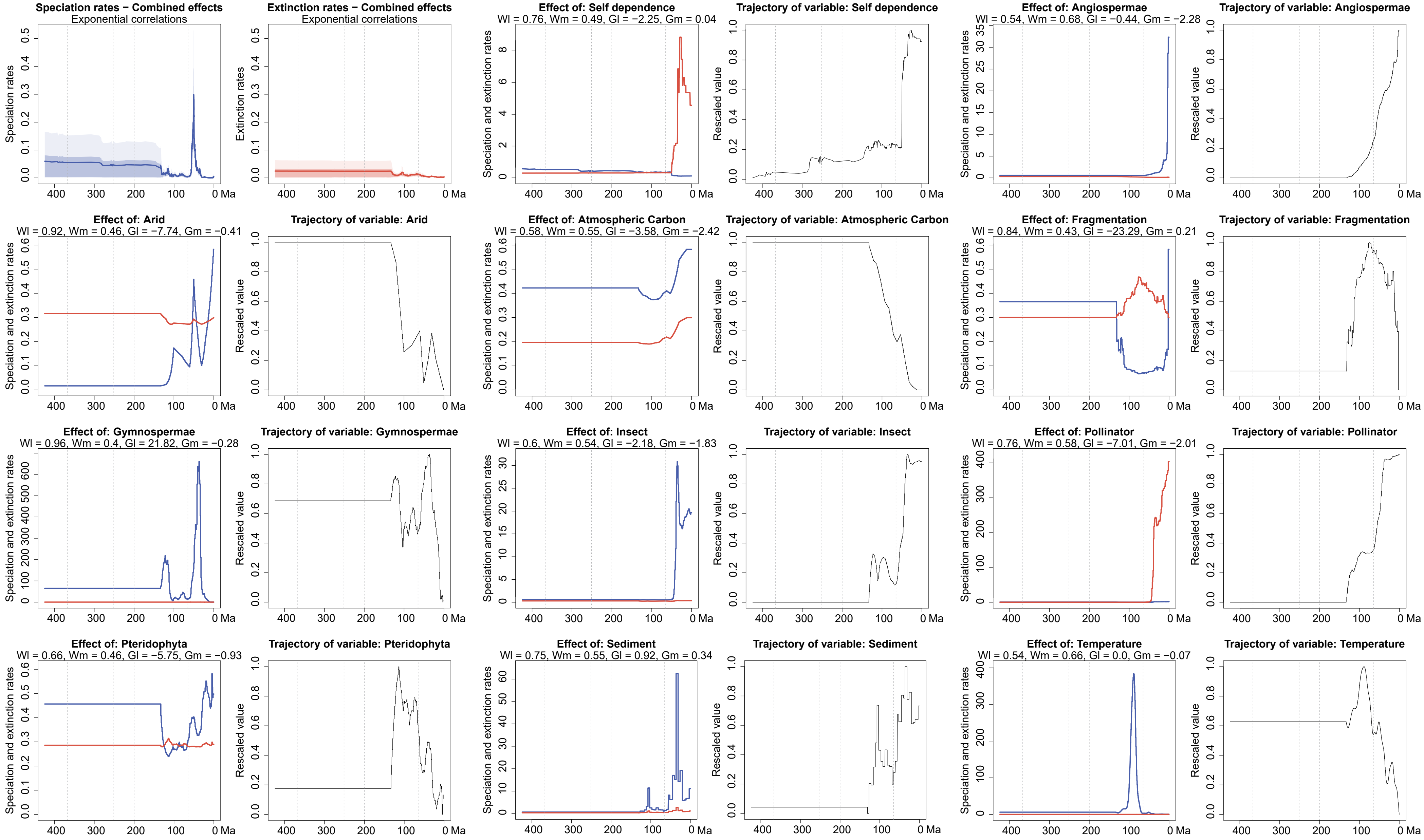

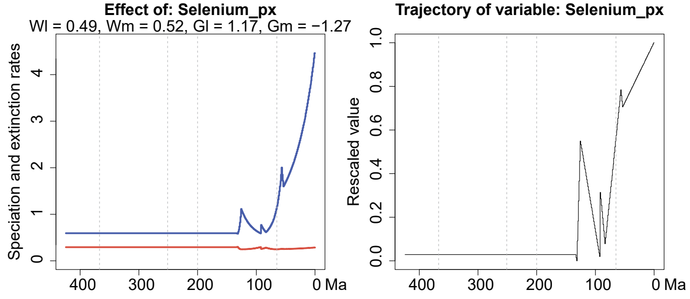

Supplementary Fig.8 The shrinkage weights (Wl and Wm for speciation and extinction, respectively) of Bryophyta during the time period from 133 to 0 Ma for all predictors are computed using MBD model. The shrinkage weights quantify the statistical support for each correlation factor. The correlation parameters are indicated by Gl and Gm.

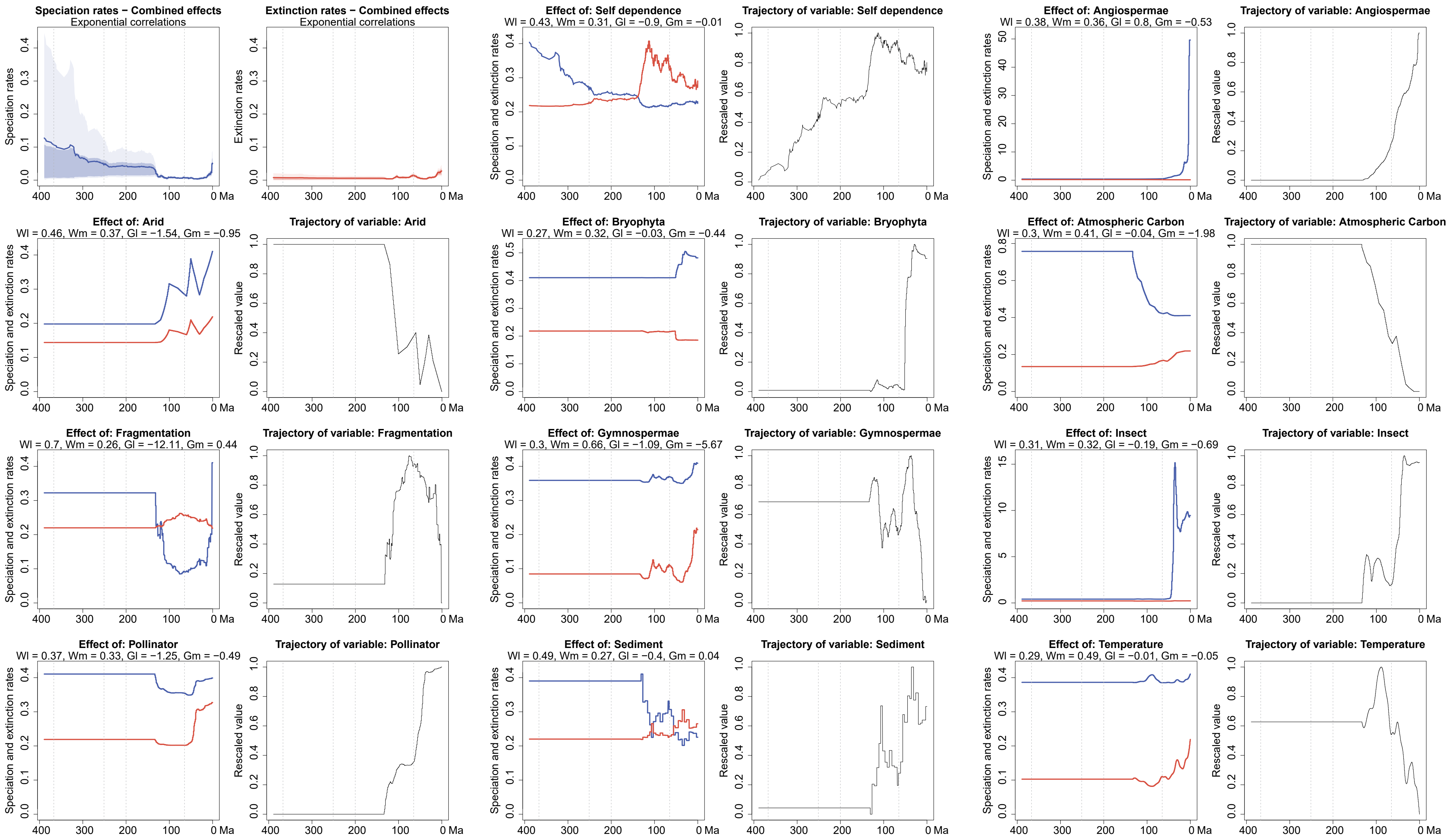

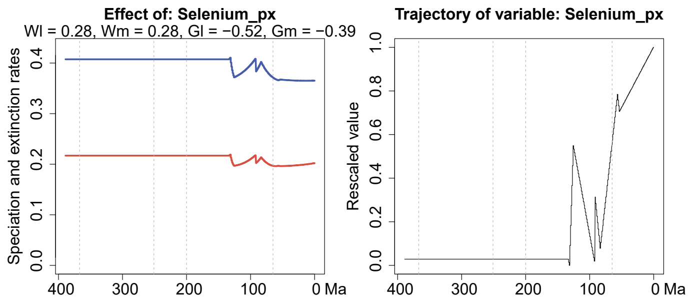

Supplementary Fig.9 The shrinkage weights (Wl and Wm for speciation and extinction, respectively) of Pteridophyta during the time period from 133 to 0 Ma for all predictors are computed using MBD model.The shrinkage weights quantify the statistical support for each correlation factor. The correlation parameters are indicated by Gl and Gm.

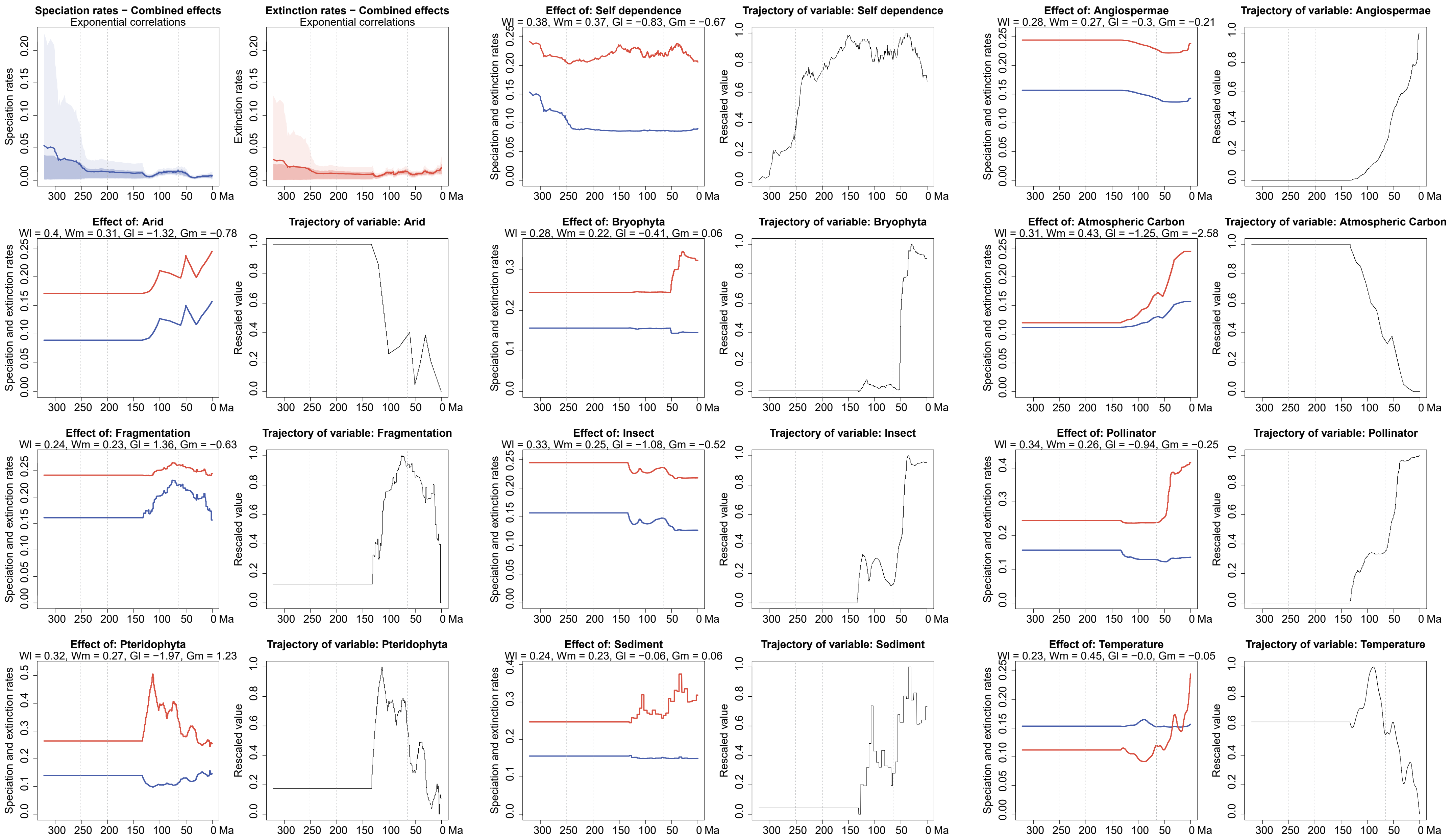

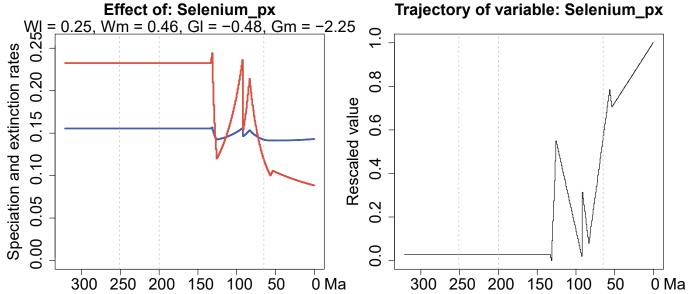

Supplementary Fig.10 The shrinkage weights (Wl and Wm for speciation and extinction, respectively) of Gymnospermae during the time period from 133 to 0 Ma for all predictors are computed using MBD model.The shrinkage weights quantify the statistical support for each correlation factor. The correlation parameters are indicated by Gl and Gm.

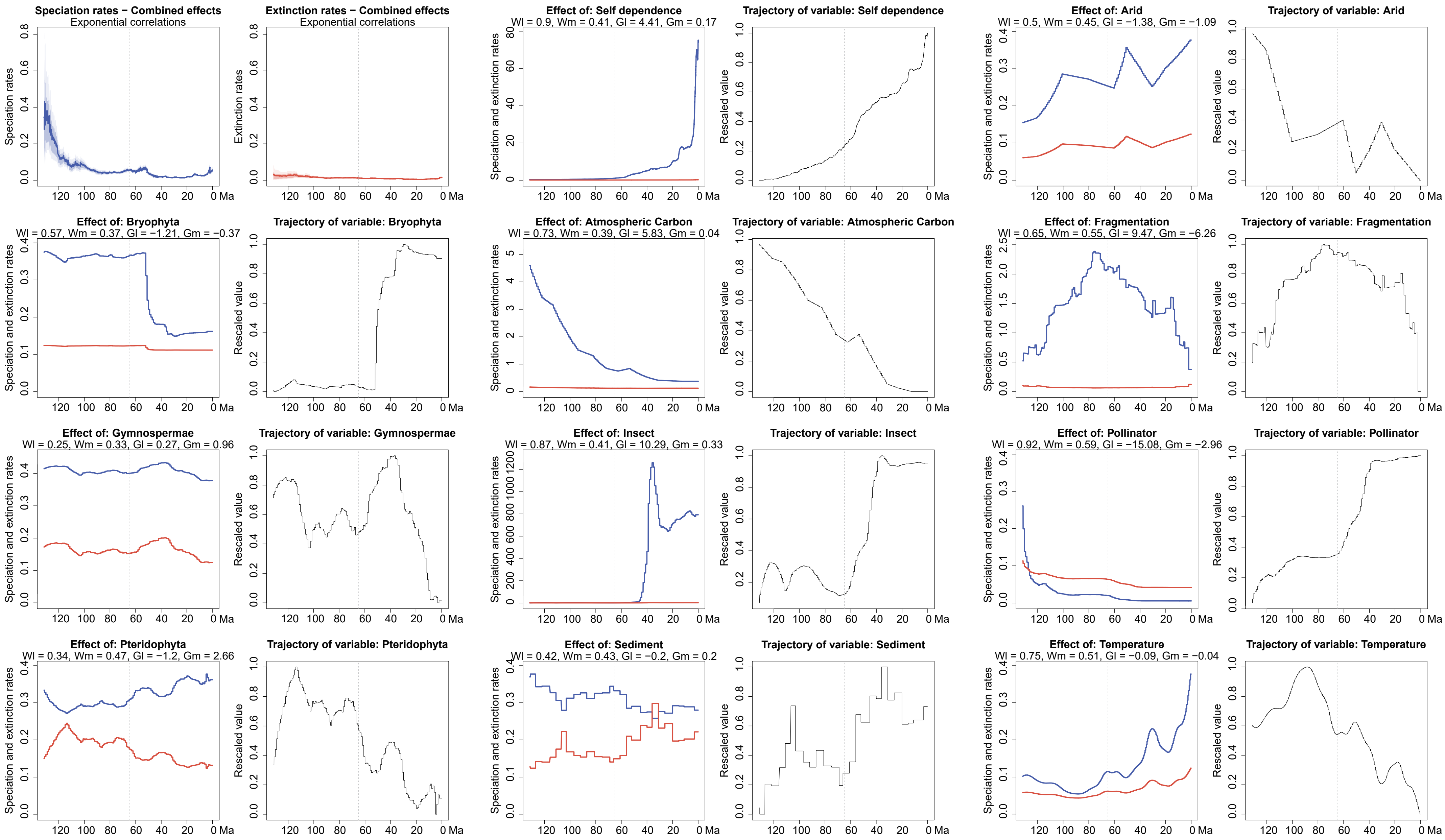

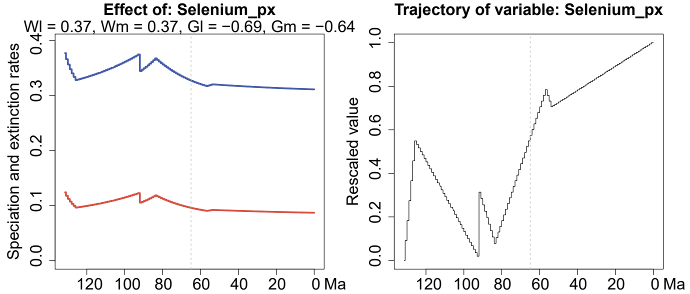

Supplementary Fig.11 The shrinkage weights (Wl and Wm for speciation and extinction, respectively) of Angiospermae during the time period from 133 to 0 Ma for all predictors are computed using MBD model.The shrinkage weights quantify the statistical support for each correlation factor. The correlation parameters are indicated by Gl and Gm.

**Supplementary Table 1**. Results of the multivariate birth-death model applied to Bryophyta in two set of analyses. The table reports the posterior parameter estimates and 95% credibility interval (CI) for the MBD correlation parameters associated with each predictor: G$\lambda$ for origination and G$\mu$ for extinction. A positive G implies that the factor correlates positively with the rate and *vice versa*. Shrinkage weights (ω) quantify support for retaining each predictor in the model, with values greater than 0.5 highlighted in bold. (a) Diversification from the origin of angiosperms to the present (133-0 Ma); (b) Diversification during Permian to the Early Cretaceous (300-100 Ma).

| | **Parameters** | **Mean (a)** | **Median (a)** | **95%CI (a)** | **Mean (b)** | **Median (b)** | **95%CI (b)** |
|---|---|---|---|---|---|---|---|
| **Bryophyta** | G λ Self dependence | -2.1676 | -2.2233 | [-4.1972，0.1006] | -2.6582 | -0.4567 | [-16.4596,4.8667] |
| | G λ Angiospermae | -0.3625 | -0.0858 | [-5.1554,3.6124] | —— | —— | —— |
| | G λ Arid | -7.6299 | -7.7931 | [-12.2697,-2.7393] | -0.525 | -0.2237 | [-3.2871,1.4228] |
| | G λ Atmospheric Carbon | -3.5608 | -1.6774 | [-15.7745,4.9811] | 4.1667 | 3.836 | [-1.1255,10.5046] |
| | G λ Continental fragmention | -22.6442 | -22.5117 | [-46.5688,1.8029] | -1.2368 | -0.0369 | [-23.9732,17.0317] |
| | G λ Gymnospermae | 21.8882 | 21.7505 | [15.708,28.1665] | -5.7511 | -5.7571 | [-10.0491,-1.4531] |
| | G λ Insect | -2.205 | -0.9227 | [-13.6634,6.3207] | 2.4313 | 1.4515 | [-3.2802,10.9807] |
| | G λ Pollinator | -7.1073 | -6.0254 | [-21.2996,2.1738] | 3.7284 | 1.9484 | [-3.6631,15.9865] |
| | G λ Pteridophyta | -5.8733 | -5.6741 | [-14.8429,1.5471] | -0.6384 | -0.2188 | [-5.0404,2.764] |
| | G λ Sediment | 0.9208 | 0.8914 | [-0.1284,2.1151] | -0.0408 | -0.0097 | [-0.6411,0.4691] |
| | G λ Temperature | 0.0024 | -0.0011 | [-0.1573,0.1778] | 0.0223 | 0.0079 | [-0.0629,0.1312] |
| | G λ Selenium | 0.9664 | 0.447 | [-2.3663,5.7668] | -1.5522 | -1.3197 | [-4.7671,0.7915] |
| | G μ Self dependence | 0.0813 | -0.0007 | [-2.7984,3.0185] | 10.3992 | 2.8047 | [-4.2914,41.8218] |
| | G μ Angiospermae | -2.2559 | -1.6874 | [-7.4261,1.186] | —— | —— | —— |
| | G μ Arid | -0.416 | -0.1805 | [-3.7833,2.4255] | 1.1044 | 0.7086 | [-1.2701,4.6429] |
| | G μ Atmospheric Carbon | -2.6034 | -1.6945 | [-10.326,2.7205] | 1.0279 | 0.3792 | [-4.021,8.2472] |
| | G μ Continental fragmention | 0.0991 | 0.0146 | [-11.1867,11.5933] | 1.3911 | 0.1032 | [-21.1685,26.868] |
| | G μ Gymnospermae | -0.3555 | -0.1245 | [-4.855,3.703] | -9.5304 | -9.4566 | [-15.2865,-3.4201] |
| | G μ Insect | -1.8823 | -1.0013 | [-9.4735,3.3456] | -0.9022 | -0.2125 | [-9.521,6.0269] |
| | G μ Pollinator | -2.1757 | -0.8103 | [-12.2485,4.1605] | 1.5388 | 0.3498 | [-6.8249,13.0226] |
| | G μ Pteridophyta | -0.7921 | -0.3226 | [-8.0493,4.9251] | 5.7515 | 5.1981 | [-1.5865,15.5238] |
| | G μ Sediment | 0.3681 | 0.2491 | [-0.3994,1.3695] | -0.4448 | -0.3201 | [-1.5011,0.3012] |
| | G μ Temperature | -0.0626 | -0.0552 | [-0.1792,0.0244] | -0.0176 | -0.0063 | [-0.1455,0.1032] |
| | G μ Selenium | -1.2041 | -0.6267 | [-6.7327,2.7032] | -3.4593 | -3.2425 | [-8.1341,0.4449] |

| | Parameters | Mean (a) | Median (a) | 95%CI (a) | Mean (b) | Median (b) | 95%CI (b) |
|---|---|---|---|---|---|---|---|
| Bryophyta | ω λ Self dependence | **0.7503** | **0.8391** | **[0.1646,1]** | **0.6308** | **0.7586** | **[0.0146,1]** |
| | ω λ Angiospermae | **0.5364** | **0.5904** | **[0,0.983]** | —— | —— | —— |
| | ω λ Arid | **0.9163** | **0.9502** | **[0.7357,1]** | **0.5036** | **0.526** | **[0,0.9759]** |
| | ω λ Atmospheric Carbon | **0.5843** | **0.6742** | **[0.0121,0.9998]** | **0.7214** | **0.8467** | **[0.0515,1]** |
| | ω λ Continental fragmention | **0.8303** | **0.9156** | **[0.2834,1]** | **0.5367** | **0.5879** | **[0,0.9826]** |
| | ω λ Gymnospermae | **0.9578** | **0.9709** | **[0.874,1]** | **0.9142** | **0.9564** | **[0.688,1]** |
| | ω λ Insect | **0.604** | **0.7121** | **[0.0135,1]** | **0.6156** | **0.7307** | **[0.0151,0.9999]** |
| | ω λ Pollinator | **0.7648** | **0.9258** | **[0.0452,1]** | **0.6408** | **0.7757** | **[0.0169,1]** |
| | ω λ Pteridophyta | **0.6639** | **0.7679** | **[0.0382,1]** | **0.516** | **0.5555** | **[0,0.9774]** |
| | ω λ Sediment | **0.7486** | **0.8631** | **[0.0858,1]** | 0.4729 | 0.4747 | [0,0.9678] |
| | ω λ Temperature | **0.5402** | **0.5922** | **[0,0.9835]** | 0.4945 | 0.5087 | [0,0.9769] |
| | ω λ Selenium | 0.4735 | 0.4718 | [0,0.9681] | **0.6332** | **0.7464** | **[0.0216,0.9999]** |
| | ω μ Self dependence | 0.4905 | 0.502 | [0,0.9746] | **0.7399** | **0.9465** | **[0.026,1]** |
| | ω μ Angiospermae | **0.6753** | **0.8194** | **[0.025,1]** | —— | —— | —— |
| | ω μ Arid | 0.4586 | 0.4443 | [0,0.9625] | **0.5867** | **0.6754** | **[0,0.9851]** |
| | ω μ Atmospheric Carbon | **0.5624** | **0.6384** | **[0,0.9809]** | **0.5441** | **0.6033** | **[0,0.9817]** |
| | ω μ Continental fragmention | 0.4337 | 0.3986 | [0,0.9594] | **0.5508** | **0.6138** | **[0,0.9855]** |
| | ω μ Gymnospermae | 0.399 | 0.3403 | [0,0.9486] | **0.9652** | **0.982** | **[0.8847,1]** |
| | ω μ Insect | **0.5465** | **0.6089** | **[0,0.9805]** | **0.552** | **0.6136** | **[0,0.985]** |
| | ω μ Pollinator | **0.5859** | **0.6821** | **[0.0115,1]** | **0.5861** | **0.6781** | **[0.0117,1]** |
| | ω μ Pteridophyta | 0.4497 | 0.4295 | [0,0.9629] | **0.7849** | **0.9309** | **[0.0634,1]** |
| | ω μ Sediment | **0.5601** | **0.6363** | **[0,0.9819]** | **0.6356** | **0.7629** | **[0.0187,1]** |
| | ω μ Temperature | **0.6535** | **0.7675** | **[0.0297,1]** | **0.5197** | **0.5615** | **[0,0.9797]** |
| | ω μ Selenium | **0.5111** | **0.5436** | **[0,0.9705]** | **0.7832** | **0.9013** | **[0.0972,1]** |

**Supplementary Table 2**. Results of the multivariate birth-death model applied to Pteridophyta in two set of analyses. The table reports the posterior parameter estimates and 95% credibility interval (CI) for the MBD correlation parameters associated with each predictor: G$\lambda$ for origination and G$\mu$ for extinction. A positive G implies that the factor correlates positively with the rate and *vice versa*. Shrinkage weights ($\omega$) quantify support for retaining each predictor in the model, with values greater than 0.5 highlighted in bold. (a) Diversification from the origin of angiosperms to the present (133-0 Ma); (b) Diversification during Permian to the Early Cretaceous (300-100 Ma).

| | Parameters | Mean (a) | Median (a) | 95%CI (a) | Mean (b) | Median (b) | 95%CI (b) |
|---|---|---|---|---|---|---|---|
| **Pteridophyta** | G λ Self dependence | -0.8773 | -0.4692 | [-3.5616,0.6281] | -2.0993 | -1.8698 | [-5.8947,0.6528] |
| | G λ Angiospermae | 0.6594 | 0.1823 | [-1.0334,3.8056] | —— | —— | —— |
| | G λ Arid | -1.5685 | -1.3256 | [-4.6497,0.6122] | 0.9491 | 0.7563 | [-0.7537,3.2409] |
| | G λ Bryophyta | -0.0314 | -0.0122 | [-1.7108,1.14198] | 7.6237 | 6.5284 | [-3.1429,22.9441] |
| | G λ Atmospheric Carbon | -0.1388 | -0.041 | [-4.4213,3.9289] | 1.5554 | 1.3111 | [-1.3043,5.3138] |
| | G λ Continental fragmention | -12.4148 | -12.7832 | [-21.6278,-2.4194] | 3.8998 | 1.4942 | [-10.5124,24.431] |
| | G λ Gymnospermae | -1.0756 | -0.6044 | [-5.2445,2.0313] | -5.5648 | -5.6057 | [-8.612,-2.298] |
| | G λ Insect | -0,2695 | -0.1015 | [-4.1656,2.6118] | 0.9903 | 0.4833 | [-2.7145,6.4105] |
| | G λ Pollinator | -0.9961 | -0.2074 | [-6.2101,2.7935] | 3.5875 | 2.5277 | [-2.7232,12.9774] |
| | G λ Sediment | -0.4057 | -0.3289 | [-1.2116,0.1493] | -0.7196 | -0.7358 | [-1.347,0.0294] |
| | G λ Temperature | -0.0074 | -0.002 | [-0.0713,0.037] | -0.0396 | -0.0246 | [-0.1615,0.0362] |
| | G λ Selenium | -0.4995 | -0.1666 | [-3.4441,1.5526] | -2.1847 | -2.2051 | [-4.3907,0.1901] |
| | G μ Self dependence | -0.0128 | -0.0072 | [-2.0098,1.8042] | 1.5351 | 0.8575 | [-1.4555,6.5584] |
| | G μ Angiospermae | -0.5362 | -0.1605 | [-3.1973,1.0077] | —— | —— | —— |
| | G μ Arid | -0.9372 | -0.4227 | [-4.5074,1.3276] | 0.1677 | 0.0631 | [-1.7065,2.3335] |
| | G μ Bryophyta | -0.3969 | -0.1784 | [-2.4735,0.9507] | 2.1956 | 0.9626 | [-6.86,14.3033] |
| | G μ Atmospheric Carbon | -2.0152 | -1.206 | [-7.7377,1.6987] | 2.8151 | 2.5727 | [-0.8743,7.494] |
| | G μ Continental fragmention | 0.3692 | 0.0962 | [-6.3621,7.9316] | 5.2636 | 2.6386 | [-8.4341,25.8511] |
| | G μ Gymnospermae | -5.6738 | -5.7563 | [-10.1191,-0.1532] | -4.4286 | -4.4583 | [-8.0797,0.087] |
| | G μ Insect | -0.7252 | -0.2999 | [-4.3929,1.8055] | -2.2231 | -1.146 | [-9.8756,2.7305] |
| | G μ Pollinator | -0.5147 | -0.118 | [-4.5307,2.3952] | 0.3692 | 0.0861 | [-5.996,7.9194] |
| | G μ Sediment | 0.0336 | 0.0076 | [-0.3755,0.5021] | 0.0362 | 0.0107 | [-0.471,0.5979] |
| | G μ Temperature | -0.0442 | -0.0327 | [-0.1421,0.0178] | -0.0011 | -0.0003 | [-0.0822,0.0794] |
| | G μ Selenium | -0.3706 | -0.1017 | [-3.3062,1.9719] | -2.9702 | -2.9552 | [-6.273,0.2468] |

| | Parameters | Mean (a) | Median (a) | 95%CI (a) | Mean (b) | Median (b) | 95%CI (b) |
|---|---|---|---|---|---|---|---|
| **Pteridophyta** | **ω λ Self dependence** | 0.4265 | 0.3762 | [0,0.9621] | **0.684** | **0.8169** | **[0.0301,1]** |
| | **ω λ Angiospermae** | 0.3641 | 0.2501 | [0,0.9579] | —— | —— | —— |
| | **ω λ Arid** | 0.4639 | 0.4719 | [0,0.9589] | **0.5531** | **0.6166** | **[0,0.976]** |
| | **ω λ Bryophyta** | 0.2677 | 0.1533 | [0,0.8742] | **0.6445** | **0.761** | **[0.0255,1]** |
| | **ω λ Atmospheric Carbon** | 0.2994 | 0.1853 | [0,0.8977] | **0.552** | **0.6088** | **[0,0.9754]** |
| | **ω λ Continental fragmention** | **0.7052** | **0.7611** | **[0.232,0.9999]** | **0.5506** | **0.6152** | **[0,0.982]** |
| | **ω λ Gymnospermae** | 0.2944 | 0.1957 | [0,0.8793] | **0.9221** | **0.9523** | **[0.7495,1]** |
| | **ω λ Insect** | 0.304 | 0.1923 | [0,0.9089] | **0.5124** | **0.5506** | **[0,0.9746]** |
| | **ω λ Pollinator** | 0.3565 | 0.2452 | [0,0.958] | **0.663** | **0.8054** | **[0.0211,1]** |
| | **ω λ Sediment** | 0.4892 | 0.5173 | [0,0.9653] | **0.8042** | **0.8878** | **[0.2529,1]** |
| | **ω λ Temperature** | 0.2858 | 0.1679 | [0,0.8924] | **0.5486** | **0.6142** | **[0.0000,0.9778]** |
| | **ω λ Selenium** | 0.2806 | 0.165 | [0,0.8784] | **0.7311** | **0.82** | **[0.146,1]** |
| | **ω μ Self dependence** | 0.3077 | 0.1897 | [0,0.9135] | **0.6094** | **0.7233** | **[0.0134,0.9999]** |
| | ω μ Angiospermae | 0.3565 | 0.2478 | [0,0.9435] | —— | —— | —— |
| | ω μ Arid | 0.3701 | 0.2873 | [0,0.94] | 0.4556 | 0.443 | [0,0.9624] |
| | ω μ Bryophyta | 0.3074 | 0.2009 | [0,0.9037] | 0.4871 | 0.4992 | [0,0.9701] |
| | **ω μ Atmospheric Carbon** | 0.4137 | 0.3643 | [0,0.9558] | **0.6472** | **0.755** | **[0.0303,1]** |
| | **ω μ Continental fragmention** | 0.2539 | 0.1387 | [0,0.8614] | **0.5591** | **0.6248** | **[0,0.9842]** |
| | **ω μ Gymnospermae** | **0.6627** | **0.7138** | **[0.1619,1]** | **0.8582** | **0.9305** | **[0.4127,1]** |
| | **ω μ Insect** | 0.3194 | 0.216 | [0,0.9071] | **0.5798** | **0.6689** | **[0,0.9853]** |
| | **ω μ Pollinator** | 0.3261 | 0.2118 | [0,0.9297] | **0.5158** | **0.5491** | **[0,0.9802]** |
| | ω μ Sediment | 0.2637 | 0.1443 | [0,0.8672] | 0.4538 | 0.432 | [0,0.9655] |
| | ω μ Temperature | 0.4809 | 0.4973 | [0,0.9722] | 0.4543 | 0.4381 | [0,0.9615] |
| | **ω μ Selenium** | 0.2789 | 0.1605 | [0,0.8858] | **0.7765** | **0.8762** | **[0.148,1]** |

**Supplementary Table 3**. Results of the multivariate birth-death model applied to Gymnosperms in two set of analyses. The table reports the posterior parameter estimates and 95% credibility interval (CI) for the MBD correlation parameters associated with each predictor: $G\lambda$ for origination and $G\mu$ for extinction. A positive G implies that the factor correlates positively with the rate and *vice versa*. Shrinkage weights (ω) quantify support for retaining each predictor in the model, with values greater than 0.5 highlighted in bold. (a) Diversification from the origin of angiosperms to the present (133-0 Ma); (b) Diversification during Permian to the Early Cretaceous (300-100 Ma).

| | Parameters | Mean (a) | Median (a) | 95%CI (a) | Mean (b) | Median (b) | 95%CI (b) |
|---|---|---|---|---|---|---|---|
| **Gymnosperms** | **G λ Self dependence** | -0.8561 | -0.2991 | [-3.9545,0.888] | -7.338 | -7.2897 | [-9.5677,-5.0414] |
| | G λ Angiospermae | -0.3067 | -0.0501 | [-2.6158,1.4011] | —— | —— | —— |
| | G λ Arid | -1.3625 | -1.021 | [-4.4154,0.7127] | 1.7343 | 1.747 | [-0.2354,3.8916] |
| | G λ Bryophyta | -0.3885 | -0.1655 | [-2.3157,0.8465] | -3.8551 | -1.9115 | [-17.6524,4.6604] |
| | G λ Atmospheric Carbon | -1.2273 | -0.458 | [-6.321,1.7068] | 2.224 | 2.0286 | [-0.7566,6.0784] |
| | G λ Continental fragmention | 1.4164 | 0.4673 | [-4.5762,10.2617] | 0.7192 | 0.1725 | [-10.0174,12.6202] |
| | G λ Insect | -1.1369 | -0.5397 | [-5.2978,1.4557] | 0.0262 | 0.0047 | [-3.4727,3.8985] |
| | G λ Pollinator | -0.9623 | -0.2426 | [-5.6348,2.179] | 0.1342 | 0.0349 | [-4.5793,4.996] |
| | G λ Pteridophyta | -2.0544 | -1.2536 | [-7.6686,1.6348] | 2.8988 | 2.391 | [-0.8126,8.3644] |
| | G λ Sediment | -0.0642 | -0.0148 | [-0.564,0.3203] | -0.0676 | -0.0292 | [-0.4459,0.2596] |
| | G λ Temperature | -0.002 | -0.0002 | [-0.0567,0.0431] | -0.025 | -0.0166 | [-0.1023,0.0314] |
| | G λ Selenium | -0.4485 | -0.1479 | [-3.0876,1.5679] | -1.8091 | -1.8173 | [-3.6772,0.1292] |
| | G μ Self dependence | -0.7054 | -0.214 | [-3.8942,1.052] | -2.587 | -2.6265 | [-5.7379,0.4144] |
| | G μ Angiospermae | -0.2231 | -0.0347 | [-2.2648,1.2932] | —— | —— | —— |
| | G μ Arid | -0.8512 | -0.3813 | [-3.8277,1.0978] | 0.5467 | 0.2639 | [-0.8893,2.6958] |
| | G μ Bryophyta | 0.0791 | 0.0083 | [-1.2248,1.5869] | 0.1221 | 0.05 | [-8.9571,9.9607] |
| | G μ Atmospheric Carbon | -2.5399 | -1.6635 | [-8.6505,1.2187] | -0.1797 | -0.0352 | [-3.0337,2.3443] |
| | G μ Continental fragmention | -0.5214 | -0.1135 | [-8.5394,6.5447] | 11.2989 | 8.5989 | [-3.9207,35.5565] |
| | G μ Insect | -0.5246 | -0.1334 | [-3.5764,1.7488] | -1.7801 | -0.8056 | [-8.2859,2.1603] |
| | G μ Pollinator | -0.2745 | -0.0381 | [-3.9023,2.2388] | -0.5926 | -0.0369 | [-8.8759,5.9747] |
| | G μ Pteridophyta | 1.3805 | 0.4922 | [-2.1905,7.2371] | 0.7517 | 0.2606 | [-1.9982,4.8238] |
| | G μ Sediment | 0.0682 | 0.0181 | [-0.2514,0.5195] | -0.0921 | -0.0318 | [-0.6619,0.3725] |
| | G μ Temperature | -0.0468 | -0.029 | [-0.1603,0.0158] | -0.0055 | -0.002 | [-0.0723,0.0523] |
| | G μ Selenium | -2.2137 | -1.5747 | [-7.033,0.7926] | -1.8106 | -1.7117 | [-4.3432,0.292] |

| | Parameters | Mean (a) | Median (a) | 95%CI (a) | Mean (b) | Median (b) | 95%CI (b) |
|---|---|---|---|---|---|---|---|
| **Gymnosperms** | **ω λ Self dependence** | 0.3925 | 0.3008 | [0,0.9619] | **0.9646** | **0.9757** | **[0.8933,1]** |
| | ω λ Angiospermae | 0.2858 | 0.146 | [0,0.9159] | —— | —— | —— |
| | **ω λ Arid** | 0.4126 | 0.378 | [0,0.9481] | **0.6604** | **0.7562** | **[0.0421,1]** |
| | ω λ Bryophyta | 0.2802 | 0.1646 | [0,0.882] | 0.4643 | 0.4531 | [0,0.9705] |
| | **ω λ Atmospheric Carbon** | 0.312 | 0.189 | [0,0.9168] | **0.5699** | **0.6469** | **[0,0.9767]** |
| | ω λ Continental fragmention | 0.2457 | 0.1225 | [0,0.8511] | 0.4018 | 0.3408 | [0,0.9521] |
| | ω λ Insect | 0.3414 | 0.2399 | [0,0.9298] | 0.3923 | 0.3316 | [0,0.9486] |
| | ω λ Pollinator | 0.3453 | 0.2249 | [0,0.9483] | 0.4222 | 0.3753 | [0,0.9626] |
| | **ω λ Pteridophyta** | 0.329 | 0.2342 | [0,0.9156] | **0.6464** | **0.7773** | **[0.0189,1]** |
| | ω λ Sediment | 0.242 | 0.1158 | [0,0.8556] | 0.3693 | 0.3001 | [0,0.9308] |
| | ω λ Temperature | 0.2385 | 0.116 | [0,0.8544] | 0.4457 | 0.4322 | [0,0.9556] |
| | **ω λ Selenium** | 0.2523 | 0.1273 | [0,0.8611] | **0.6639** | **0.739** | **[0.0776,0.9999]** |
| | **ω μ Self dependence** | 0.3781 | 0.2795 | [0,0.9578] | **0.7363** | **0.8504** | **[0.0699,1]** |
| | ω μ Angiospermae | 0.2745 | 0.138 | [0,0.9018] | —— | —— | —— |
| | ω μ Arid | 0.322 | 0.2078 | [0,0.9222] | 0.4196 | 0.3759 | [0,0.9563] |
| | ω μ Bryophyta | 0.2195 | 0.0959 | [0,0.8327] | 0.3891 | 0.3233 | [0,0.9445] |
| | ω μ Atmospheric Carbon | 0.4308 | 0.3999 | [0,0.9654] | 0.3589 | 0.2739 | [0,0.9348] |
| | **ω μ Continental fragmention** | 0.2297 | 0.1075 | [0,0.8406] | **0.6397** | **0.7733** | **[0.0163,1]** |
| | ω μ Insect | 0.2486 | 0.115 | [0,0.8759] | 0.4975 | 0.5164 | [0,0.9785] |
| | ω μ Pollinator | 0.2672 | 0.131 | [0,0.906] | 0.4611 | 0.4366 | [0,0.9782] |
| | ω μ Pteridophyta | 0.2761 | 0.1427 | [0,0.8987] | 0.4466 | 0.418 | [0,0.9671] |
| | ω μ Sediment | 0.2367 | 01112 | [0,0.8483] | 0.413 | 0.3605 | [0,0.9514] |
| | ω μ Temperature | 0.4601 | 0.4567 | [0,0.9726] | 0.3687 | 0.2914 | [0,0.9373] |
| | **ω μ Selenium** | 0.4524 | 0.4474 | [0,0.9675] | **0.6352** | **0.7345** | **[0.0259,1]** |

**Supplementary Table 4**. Results of the multivariate birth-death model applied to Angiosperms in two set of analyses. The table reports the posterior parameter estimates and 95% credibility interval (CI) for the MBD correlation parameters associated with each predictor: G$\lambda$ for origination and G$\mu$ for extinction. A positive G implies that the factor correlates positively with the rate and *vice versa*. Shrinkage weights (ω) quantify support for retaining each predictor in the model, with values greater than 0.5 highlighted in bold. (a) Diversification from the origin of angiosperms to the present (133-0 Ma).

| | **Parameters** | **Mean (a)** | **Median (a)** | **95%CI (a)** |
|---|---|---|---|---|
| **Angiosperms** | G $\lambda$ Self dependence | 4.3834 | 4.4424 | [1.6131,7.0661] |
| | G $\lambda$ Arid | -1.2243 | -1.1655 | [-3.3053,0.4372] |
| | G $\lambda$ Bryophyta | -1.2429 | -1.2548 | [-2.153,-0.3952] |
| | G $\lambda$ Atmospheric Carbon | 5.6314 | 5.5706 | [-0.3664,11.72] |
| | G $\lambda$ Continental fragmention | 9.0027 | 9.0837 | [-0.2141,17.2135] |
| | G $\lambda$ Gymnospermae | 0.2466 | 0.1478 | [-1.0867,1.7787] |
| | G $\lambda$ Insect | 10.1485 | 10.6527 | [-0.3486,18.0447] |
| | G $\lambda$ Pollinator | -14.8874 | -15.7816 | [-26.3841,0.461] |
| | G $\lambda$ Pteridophyta | -1.2353 | -1.0968 | [-3.8012,0.8203] |
| | G $\lambda$ Sediment | -0.1848 | -0.1535 | [-0.586,0.1096] |
| | G $\lambda$ Temperature | -0.0828 | -0.0846 | [-0.1496,0.0029] |
| | G $\lambda$ Selenium | -0.6212 | -0.3502 | [-3.2832,1.198] |
| | G $\mu$ Self dependence | 0.2022 | 0.0454 | [-1.717,2.4966] |
| | G $\mu$ Arid | -1.0507 | -0.8333 | [-3.4228,0.7548] |
| | G $\mu$ Bryophyta | -0.3344 | -0.2184 | [-1.7449,0.8145] |
| | G $\mu$ Atmospheric Carbon | -0.0357 | -0.0068 | [-4.6006,4.5324] |
| | G $\mu$ Continental fragmention | -6.4433 | -6.5971 | [-13.8029,0.7996] |
| | G $\mu$ Gymnospermae | 0.9641 | 0.7647 | [-1.0757,3.5845] |
| | G $\mu$ Insect | 0.3018 | -0.0113 | [-4.0907,5.4809] |
| | G $\mu$ Pollinator | -3.0783 | -2.1563 | [-10.8006,1.6162] |
| | G $\mu$ Pteridophyta | 2.6264 | 2.4478 | [-0.85,6.9094] |
| | G $\mu$ Sediment | 0.2114 | 0.1656 | [-0.1721,0.7181] |
| | G $\mu$ Temperature | -0.0343 | -0.0262 | [-0.1132,0.021] |
| | G $\mu$ Selenium | -0.5893 | -0.3128 | [-3.4774,1.55] |

| Angiosperms | Parameters | Mean (a) | Median (a) | 95%CI (a) |
|---|---|---|---|---|
| | **ω λ Self dependence** | **0.8992** | **0.9385** | **[0.6728,1]** |
| | ω λ Arid | 0.4792 | 0.4857 | [0,0.9556] |
| | **ω λ Bryophyta** | **0.5745** | **0.5886** | **[0.1331,0.9999]** |
| | **ω λ Atmospheric Carbon** | **0.7226** | **0.8201** | **[0.0905,1]** |
| | **ω λ Continental fragmention** | **0.6334** | **0.6922** | **[0.0822,0.9998]** |
| | ω λ Gymnospermae | 0.2431 | 0.1143 | [0,0.8618] |
| | **ω λ Insect** | **0.8602** | **0.9502** | **[0.1849,1]** |
| | **ω λ Pollinator** | **0.9091** | **0.983** | **[0.2953,1]** |
| | ω λ Pteridophyta | 0.3461 | 0.2677 | [0,0.9079] |
| | ω λ Sediment | 0.4041 | 0.3598 | [0,0.9363] |
| | **ω λ Temperature** | **0.7323** | **0.8134** | **[0.1489,1]** |
| | ω λ Selenium | 0.3569 | 0.278 | [0,0.9264] |
| | ω μ Self dependence | 0.4018 | 0.3418 | [0,0.9493] |
| | ω μ Arid | 0.4437 | 0.4217 | [0,0.9551] |
| | ω μ Bryophyta | 0.3581 | 0.2817 | [0,0.9215] |
| | ω μ Atmospheric Carbon | 0.3836 | 0.3163 | [0,0.9352] |
| | **ω μ Continental fragmention** | **0.5557** | **0.5908** | **[0.0441,0.9996]** |
| | ω μ Gymnospermae | 0.3284 | 0.2381 | [0,0.9056] |
| | ω μ Insect | 0.4106 | 0.3551 | [0,0.9567] |
| | **ω μ Pollinator** | **0.5995** | **0.7009** | **[0.0126,1]** |
| | ω μ Pteridophyta | 0.4678 | 0.4605 | [0,0.9536] |
| | ω μ Sediment | 0.433 | 0.4098 | [0,0.9461] |
| | ω μ Temperature | 0.4926 | 0.5126 | [0,0.9668] |
| | ω μ Selenium | 0.3654 | 0.2912 | [0,0.9343] |

**Supplementary Table 5**. Time series of biotic and abiotic factors (0-133 Ma) used in the multivariate birth-death (MBD) model analyses. This table presents the temporal series of 12 predictor variables used in the MBD model to evaluate the drivers of diversification rates of four major land plant groups (Bryophytes, Pteridophytes, Gymnosperms, and Angiosperms). These data were used in the MBD analyses for Figure 5D-G.

| Factors | Age | Value | resolution | references |
|---|---|---|---|---|
| Angiospermae | 1.0053544020125 | 1 | 1 Ma | This study |
| | 2.010708804 | 0.988496732026144 | | |
| | 3.0160632060375 | 0.956993464052288 | | |
| | 4.021417608 | 0.861830065359477 | | |
| | 5.0267720100625 | 0.81921568627451 | | |
| | 6.03212641207499 | 0.799607843137255 | | |
| | 7.03748081408749 | 0.789673202614379 | | |
| | 8.04283521609999 | 0.783660130718954 | | |
| | 9.04818961811249 | 0.780915032679739 | | |
| | 10.053544020125 | 0.781176470588235 | | |
| | 11.0588984221375 | 0.779346405228758 | | |
| | 12.06425282 | 0.785490196078431 | | |
| | 13.0696072261625 | 0.785228758169935 | | |
| | 14.074961628175 | 0.771372549019608 | | |
| | 15.0803160301875 | 0.747843137254902 | | |
| | 16.08567043 | 0.725882352941177 | | |
| | 17.0910248342125 | 0.70640522875817 | | |
| | 18.096379236225 | 0.690196078431373 | | |
| | 19.1017336382375 | 0.678823529411765 | | |
| | 20.10708804 | 0.668758169934641 | | |
| | 21.1124424422625 | 0.656209150326797 | | |
| | 22.117796844275 | 0.642222222222222 | | |
| | 23.1231512462875 | 0.632679738562092 | | |
| | 24.12850565 | 0.623398692810458 | | |
| | 25.1338600503125 | 0.616209150326797 | | |
| | 26.139214452325 | 0.611764705882353 | | |
| | 27.1445688543375 | 0.608235294117647 | | |
| | 28.14992326 | 0.606797385620915 | | |
| | 29.1552776583625 | 0.603137254901961 | | |
| | 30.160632060375 | 0.597647058823529 | | |
| | 31.1659864623875 | 0.593202614379085 | | |
| | 32.17134086 | 0.590588235294118 | | |
| | 33.1766952664125 | 0.589542483660131 | | |
| | 34.182049668425 | 0.592679738562092 | | |
| | 35.1874040704375 | 0.589019607843137 | | |
| | 36.19275847 | 0.583267973856209 | | |
| | 37.1981128744625 | 0.576993464052288 | | |

| | | |
|---|---|---|
| | 38.203467276475 | 0.57202614379085 |
| | 39.2088216784875 | 0.564575163398693 |
| | 40.21417608 | 0.553856209150327 |
| | 41.2195304825125 | 0.545882352941176 |
| | 42.224884884525 | 0.534117647058824 |
| | 43.2302392865375 | 0.523267973856209 |
| | 44.23559369 | 0.51359477124183 |
| | 45.2409480905625 | 0.505490196078431 |
| | 46.246302492575 | 0.501960784313726 |
| | 47.2516568945875 | 0.497124183006536 |
| | 48.2570113 | 0.491111111111111 |
| | 49.2623656986125 | 0.479346405228758 |
| | 50.267720100625 | 0.467058823529412 |
| | 51.2730745026375 | 0.458823529411765 |
| | 52.2784289 | 0.440392156862745 |
| | 53.2837833066625 | 0.423921568627451 |
| | 54.289137708675 | 0.410588235294118 |
| | 55.2944921106875 | 0.397385620915033 |
| | 56.29984651 | 0.37843137254902 |
| | 57.3052009147125 | 0.343137254901961 |
| | 58.310555316725 | 0.324967320261438 |
| | 59.3159097187375 | 0.302091503267974 |
| | 60.32126412 | 0.286666666666667 |
| | 61.3266185227624 | 0.277385620915033 |
| | 62.3319729247749 | 0.267058823529412 |
| | 63.3373273267874 | 0.261045751633987 |
| | 64.3426817287999 | 0.25437908496732 |
| | 65.3480361308124 | 0.247712418300654 |
| | 66.3533905328249 | 0.239869281045752 |
| | 67.3587449348374 | 0.226274509803922 |
| | 68.3640993368499 | 0.212679738562092 |
| | 69.3694537388624 | 0.206143790849673 |
| | 70.3748081408749 | 0.200130718954248 |
| | 71.3801625428874 | 0.193856209150327 |
| | 72.3855169448999 | 0.189150326797386 |
| | 73.3908713469124 | 0.183137254901961 |
| | 74.3962257489249 | 0.173856209150327 |
| | 75.4015801509374 | 0.164183006535948 |
| | 76.4069345529499 | 0.160653594771242 |
| | 77.4122889549624 | 0.152549019607843 |
| | 78.4176433569749 | 0.146928104575163 |
| | 79.4229977589874 | 0.14156862745098 |
| | 80.4283521609999 | 0.136601307189542 |

| | | |
|---|---|---|
| | 81.4337065630124 | 0.13281045751634 |
| | 82.4390609650249 | 0.128496732026144 |
| | 83.4444153670374 | 0.124575163398693 |
| | 84.4497697690499 | 0.122352941176471 |
| | 85.4551241710624 | 0.120392156862745 |
| | 86.4604785730749 | 0.117385620915033 |
| | 87.4658329750874 | 0.115686274509804 |
| | 88.4711873770999 | 0.110718954248366 |
| | 89.4765417791124 | 0.10797385620915 |
| | 90.4818961811249 | 0.104313725490196 |
| | 91.4872505831374 | 0.0993464052287582 |
| | 92.4926049851499 | 0.0959477124183007 |
| | 93.4979593871624 | 0.0924183006535948 |
| | 94.5033137891749 | 0.0895424836601307 |
| | 95.5086681911874 | 0.0844444444444444 |
| | 96.5140225931999 | 0.0815686274509804 |
| | 97.5193769952124 | 0.0797385620915033 |
| | 98.5247313972249 | 0.0764705882352941 |
| | 99.5300857992374 | 0.0734640522875817 |
| | 100.53544020125 | 0.070718954248366 |
| | 101.540794603262 | 0.0649673202614379 |
| | 102.546149005275 | 0.0584313725490196 |
| | 103.551503407287 | 0.0525490196078431 |
| | 104.5568578 | 0.0501960784313726 |
| | 105.562212211312 | 0.0471895424836601 |
| | 106.567566613325 | 0.0428758169934641 |
| | 107.572921015337 | 0.0402614379084967 |
| | 108.57827541735 | 0.0393464052287582 |
| | 109.583629819362 | 0.0379084967320261 |
| | 110.588984221375 | 0.0349019607843137 |
| | 111.594338623387 | 0.0326797385620915 |
| | 112.599693 | 0.0311111111111111 |
| | 113.605047427412 | 0.0283660130718954 |
| | 114.610401829425 | 0.0244444444444444 |
| | 115.615756231437 | 0.0210457516339869 |
| | 116.62111063345 | 0.0177777777777778 |
| | 117.626465035462 | 0.0150326797385621 |
| | 118.631819437475 | 0.0125490196078431 |
| | 119.637173839487 | 0.0105882352941176 |
| | 120.6425282 | 0.0100653594771242 |
| | 121.647882643512 | 0.0104575163398693 |
| | 122.653237045525 | 0.0104575163398693 |
| | 123.658591447537 | 0.00967320261437908 |

| | | | | |
|---|---|---|---|---|
| | 124.66394584955 | 0.00875816993464052 | | |
| | 125.669300251562 | 0.00797385620915033 | | |
| | 126.674654653575 | 0.00758169934640523 | | |
| | 127.680009055587 | 0.0061437908496732 | | |
| | 128.6853635 | 0.00313725490196078 | | |
| | 129.690717859612 | 0.00117647058823529 | | |
| | 130.696072261625 | 0.00104575163398693 | | |
| | 131.701426663637 | 0.000522875816993464 | | |
| | 132.70678106565 | 0.000261437908496732 | | |
| | 133.712135467662 | 0 | | |
| Arid | 0 | 0.06 | 1 Ma | Lehtonen et al. (2017) |
| | 1 | 0.0665 | | |
| | 2 | 0.073 | | |
| | 3 | 0.0795 | | |
| | 4 | 0.086 | | |
| | 5 | 0.0925 | | |
| | 6 | 0.099 | | |
| | 7 | 0.1055 | | |
| | 8 | 0.112 | | |
| | 9 | 0.1185 | | |
| | 10 | 0.125 | | |
| | 11 | 0.1315 | | |
| | 12 | 0.138 | | |
| | 13 | 0.1445 | | |
| | 14 | 0.151 | | |
| | 15 | 0.1575 | | |
| | 16 | 0.164 | | |
| | 17 | 0.1705 | | |
| | 18 | 0.177 | | |
| | 19 | 0.1835 | | |
| | 20 | 0.19 | | |
| | 21 | 0.201 | | |
| | 22 | 0.212 | | |
| | 23 | 0.223 | | |
| | 24 | 0.234 | | |
| | 25 | 0.245 | | |
| | 26 | 0.256 | | |
| | 27 | 0.267 | | |
| | 28 | 0.278 | | |
| | 29 | 0.289 | | |
| | 30 | 0.3 | | |
| | 31 | 0.288 | | |
| | 32 | 0.276 | | |

| | | |
|---|---|---|
| | 33 | 0.264 |
| | 34 | 0.252 |
| | 35 | 0.24 |
| | 36 | 0.228 |
| | 37 | 0.216 |
| | 38 | 0.204 |
| | 39 | 0.192 |
| | 40 | 0.18 |
| | 41 | 0.171 |
| | 42 | 0.162 |
| | 43 | 0.153 |
| | 44 | 0.144 |
| | 45 | 0.135 |
| | 46 | 0.126 |
| | 47 | 0.117 |
| | 48 | 0.108 |
| | 49 | 0.099 |
| | 50 | 0.09 |
| | 51 | 0.112 |
| | 52 | 0.134 |
| | 53 | 0.156 |
| | 54 | 0.178 |
| | 55 | 0.2 |
| | 56 | 0.222 |
| | 57 | 0.244 |
| | 58 | 0.266 |
| | 59 | 0.288 |
| | 60 | 0.31 |
| | 61 | 0.307 |
| | 62 | 0.304 |
| | 63 | 0.301 |
| | 64 | 0.298 |
| | 65 | 0.295 |
| | 66 | 0.292 |
| | 67 | 0.289 |
| | 68 | 0.286 |
| | 69 | 0.283 |
| | 70 | 0.28 |
| | 71 | 0.277 |
| | 72 | 0.274 |
| | 73 | 0.271 |
| | 74 | 0.268 |
| | 75 | 0.265 |

| | | |
|---|---|---|
| | 76 | 0.262 |
| | 77 | 0.259 |
| | 78 | 0.256 |
| | 79 | 0.253 |
| | 80 | 0.25 |
| | 81 | 0.2485 |
| | 82 | 0.247 |
| | 83 | 0.2455 |
| | 84 | 0.244 |
| | 85 | 0.2425 |
| | 86 | 0.241 |
| | 87 | 0.2395 |
| | 88 | 0.238 |
| | 89 | 0.2365 |
| | 90 | 0.235 |
| | 91 | 0.2335 |
| | 92 | 0.232 |
| | 93 | 0.2305 |
| | 94 | 0.229 |
| | 95 | 0.2275 |
| | 96 | 0.226 |
| | 97 | 0.2245 |
| | 98 | 0.223 |
| | 99 | 0.2215 |
| | 100 | 0.22 |
| | 101 | 0.239 |
| | 102 | 0.258 |
| | 103 | 0.277 |
| | 104 | 0.296 |
| | 105 | 0.315 |
| | 106 | 0.334 |
| | 107 | 0.353 |
| | 108 | 0.372 |
| | 109 | 0.391 |
| | 110 | 0.41 |
| | 111 | 0.429 |
| | 112 | 0.448 |
| | 113 | 0.467 |
| | 114 | 0.486 |
| | 115 | 0.505 |
| | 116 | 0.524 |
| | 117 | 0.543 |
| | 118 | 0.562 |

| | | | | |
|---|---|---|---|---|
| | 119 | 0.581 | | |
| | 120 | 0.6 | | |
| | 121 | 0.6065 | | |
| | 122 | 0.613 | | |
| | 123 | 0.6195 | | |
| | 124 | 0.626 | | |
| | 125 | 0.6325 | | |
| | 126 | 0.639 | | |
| | 127 | 0.6455 | | |
| | 128 | 0.652 | | |
| | 129 | 0.6585 | | |
| | 130 | 0.665 | | |
| | 131 | 0.6715 | | |
| | 132 | 0.678 | | |
| | 133 | 0.6845 | | |
| Atmospheric Carbon | 0 | 0 | 1 Ma | Berner (2001) |
| | 1 | 0 | | |
| | 2 | 0 | | |
| | 3 | 0 | | |
| | 4 | 0 | | |
| | 5 | 0 | | |
| | 6 | 0 | | |
| | 7 | 0 | | |
| | 8 | 0 | | |
| | 9 | 0 | | |
| | 10 | 0 | | |
| | 11 | 0 | | |
| | 12 | 0 | | |
| | 13 | 0.000909090909090909 | | |
| | 14 | 0.00181818181818182 | | |
| | 15 | 0.00272727272727273 | | |
| | 16 | 0.00363636363636364 | | |
| | 17 | 0.00454545454545455 | | |
| | 18 | 0.00545454545454545 | | |
| | 19 | 0.00636363636363636 | | |
| | 20 | 0.00727272727272727 | | |
| | 21 | 0.00818181818181818 | | |
| | 22 | 0.00909090909090909 | | |
| | 23 | 0.01 | | |
| | 24 | 0.01125 | | |
| | 25 | 0.0125 | | |
| | 26 | 0.01375 | | |
| | 27 | 0.015 | | |

| | | |
|---|---|---|
| | 28 | 0.01625 |
| | 29 | 0.0175 |
| | 30 | 0.01875 |
| | 31 | 0.02 |
| | 32 | 0.0255555555555556 |
| | 33 | 0.0311111111111111 |
| | 34 | 0.0366666666666667 |
| | 35 | 0.0422222222222222 |
| | 36 | 0.0477777777777778 |
| | 37 | 0.0533333333333333 |
| | 38 | 0.0588888888888889 |
| | 39 | 0.0644444444444444 |
| | 40 | 0.07 |
| | 41 | 0.0761538461538462 |
| | 42 | 0.0823076923076923 |
| | 43 | 0.0884615384615385 |
| | 44 | 0.0946153846153846 |
| | 45 | 0.100769230769231 |
| | 46 | 0.106923076923077 |
| | 47 | 0.113076923076923 |
| | 48 | 0.119230769230769 |
| | 49 | 0.125384615384615 |
| | 50 | 0.131538461538462 |
| | 51 | 0.137692307692308 |
| | 52 | 0.143846153846154 |
| | 53 | 0.15 |
| | 54 | 0.147777777777778 |
| | 55 | 0.145555555555556 |
| | 56 | 0.143333333333333 |
| | 57 | 0.141111111111111 |
| | 58 | 0.138888888888889 |
| | 59 | 0.136666666666667 |
| | 60 | 0.134444444444444 |
| | 61 | 0.132222222222222 |
| | 62 | 0.13 |
| | 63 | 0.132222222222222 |
| | 64 | 0.134444444444444 |
| | 65 | 0.136666666666667 |
| | 66 | 0.138888888888889 |
| | 67 | 0.141111111111111 |
| | 68 | 0.143333333333333 |
| | 69 | 0.145555555555556 |
| | 70 | 0.147777777777778 |

| | | |
|---|---|---|
| | 71 | 0.15 |
| | 72 | 0.156363636363636 |
| | 73 | 0.162727272727273 |
| | 74 | 0.169090909090909 |
| | 75 | 0.175454545454545 |
| | 76 | 0.181818181818182 |
| | 77 | 0.188181818181818 |
| | 78 | 0.194545454545455 |
| | 79 | 0.200909090909091 |
| | 80 | 0.207272727272727 |
| | 81 | 0.213636363636364 |
| | 82 | 0.22 |
| | 83 | 0.221818181818182 |
| | 84 | 0.223636363636364 |
| | 85 | 0.225454545454545 |
| | 86 | 0.227272727272727 |
| | 87 | 0.229090909090909 |
| | 88 | 0.230909090909091 |
| | 89 | 0.232727272727273 |
| | 90 | 0.234545454545455 |
| | 91 | 0.236363636363636 |
| | 92 | 0.238181818181818 |
| | 93 | 0.24 |
| | 94 | 0.245555555555556 |
| | 95 | 0.251111111111111 |
| | 96 | 0.256666666666667 |
| | 97 | 0.262222222222222 |
| | 98 | 0.267777777777778 |
| | 99 | 0.273333333333333 |
| | 100 | 0.278888888888889 |
| | 101 | 0.284444444444444 |
| | 102 | 0.29 |
| | 103 | 0.294545454545455 |
| | 104 | 0.299090909090909 |
| | 105 | 0.303636363636364 |
| | 106 | 0.308181818181818 |
| | 107 | 0.312727272727273 |
| | 108 | 0.317272727272727 |
| | 109 | 0.321818181818182 |
| | 110 | 0.326363636363636 |
| | 111 | 0.330909090909091 |
| | 112 | 0.335454545454545 |
| | 113 | 0.34 |

| | | | | |
|---|---|---|---|---|
| | 114 | 0.34125 | | |
| | 115 | 0.3425 | | |
| | 116 | 0.34375 | | |
| | 117 | 0.345 | | |
| | 118 | 0.34625 | | |
| | 119 | 0.3475 | | |
| | 120 | 0.34875 | | |
| | 121 | 0.35 | | |
| | 122 | 0.353636363636364 | | |
| | 123 | 0.357272727272727 | | |
| | 124 | 0.360909090909091 | | |
| | 125 | 0.364545454545455 | | |
| | 126 | 0.368181818181818 | | |
| | 127 | 0.371818181818182 | | |
| | 128 | 0.375454545454545 | | |
| | 129 | 0.379090909090909 | | |
| | 130 | 0.382727272727273 | | |
| | 131 | 0.386363636363636 | | |
| | 132 | 0.39 | | |
| | 133 | 0.399166666666667 | | |
| Bryophyta | 1.00052749635994 | 0.922823525087638 | 1 Ma | This study |
| | 2.00105499271987 | 0.922823525087638 | | |
| | 3.00158248907981 | 0.922823525087638 | | |
| | 4.00210998543974 | 0.932517676895299 | | |
| | 5.00263748179968 | 0.938458216367955 | | |
| | 6.00316497815962 | 0.942395604092678 | | |
| | 7.00369247451955 | 0.942395604092678 | | |
| | 8.00421997087949 | 0.942395604092678 | | |
| | 9.00474746723943 | 0.942395604092678 | | |
| | 10.0052749635994 | 0.943351236119215 | | |
| | 11.0058024599593 | 0.944352811993182 | | |
| | 12.0063299563192 | 0.944352811993182 | | |
| | 13.0068574526792 | 0.946280156392857 | | |
| | 14.0073849490391 | 0.946310019893686 | | |
| | 15.007912445399 | 0.948232769908618 | | |
| | 16.008439941759 | 0.949209076666498 | | |
| | 17.0089674381189 | 0.949245831744442 | | |
| | 18.0094949344789 | 0.953973453644955 | | |
| | 19.0100224308388 | 0.955073808790896 | | |
| | 20.0105499271987 | 0.958985927399532 | | |
| | 21.0110774235587 | 0.95717114542606 | | |
| | 22.0116049199186 | 0.962643057654934 | | |
| | 23.0121324162785 | 0.969690843850646 | | |

| | | |
|---|---|---|
| | 24.0126599126385 | 0.970775118649986 |
| | 25.0131874089984 | 0.975668138401246 |
| | 26.0137149053583 | 0.990048562646733 |
| | 27.0142424017183 | 0.996280845550568 |
| | 28.0147698980782 | 0.998047386484239 |
| | 29.0152973944382 | 1 |
| | 30.0158248907981 | 0.9886656528391 |
| | 31.016352387158 | 0.967501619520622 |
| | 32.016879883518 | 0.961283119771016 |
| | 33.0174073798779 | 0.964600265555438 |
| | 34.0179348762378 | 0.966194517061248 |
| | 35.0184623725978 | 0.921079956077682 |
| | 36.0189898689577 | 0.8568022163312 |
| | 37.0195173653176 | 0.827515310787156 |
| | 38.0200448616776 | 0.821310594191779 |
| | 39.0205723580375 | 0.82104871426143 |
| | 40.0210998543974 | 0.82104871426143 |
| | 41.0216273507574 | 0.819279876135387 |
| | 42.0221548471173 | 0.818995024281323 |
| | 43.0226823434773 | 0.820949934989456 |
| | 44.0232098398372 | 0.815783549345989 |
| | 45.0237373361971 | 0.797410604758864 |
| | 46.0242648325571 | 0.762249778320936 |
| | 47.024792328917 | 0.729916795692305 |
| | 48.0253198252769 | 0.698544039474954 |
| | 49.0258473216369 | 0.642968064431652 |
| | 50.0263748179968 | 0.550995373454564 |
| | 51.0269023143567 | 0.349485658628026 |
| | 52.0274298107167 | 0.202906407788401 |
| | 53.0279573070766 | 0.202571017702164 |
| | 54.0284848034366 | 0.201592413751912 |
| | 55.0290122997965 | 0.205254138392057 |
| | 56.0295397961564 | 0.207206751907818 |
| | 57.0300672925164 | 0.206354493537998 |
| | 58.0305947888763 | 0.204794699917761 |
| | 59.0311222852362 | 0.208036038353924 |
| | 60.0316497815962 | 0.21039984930418 |
| | 61.0321772779561 | 0.215012611586119 |
| | 62.032704774316 | 0.217107651028912 |
| | 63.033232270676 | 0.214603711343995 |
| | 64.0337597670359 | 0.21529286905544 |
| | 65.0342872633959 | 0.222225795632578 |
| | 66.0348147597558 | 0.223948689911191 |

| | | |
|---|---|---|
| | 67.0353422561157 | 0.227552984742048 |
| | 68.0358697524757 | 0.230794323178212 |
| | 69.0363972488356 | 0.231929136209724 |
| | 70.0369247451955 | 0.231929136209724 |
| | 71.0374522415555 | 0.230135028967596 |
| | 72.0379797379154 | 0.230619736557979 |
| | 73.0385072342753 | 0.22882562931585 |
| | 74.0390347306353 | 0.229801936073731 |
| | 75.0395622269952 | 0.228993324358968 |
| | 76.0400897233552 | 0.226581272368911 |
| | 77.0406172197151 | 0.22507890855796 |
| | 78.041144716075 | 0.215030989125091 |
| | 79.041672212435 | 0.213335661154936 |
| | 80.0421997087949 | 0.218472183297574 |
| | 81.0427272051548 | 0.222935628075366 |
| | 82.0432547015148 | 0.222744961108533 |
| | 83.0437821978747 | 0.221545826690619 |
| | 84.0443096942346 | 0.221164492756952 |
| | 85.0448371905946 | 0.2201858888067 |
| | 86.0453646869545 | 0.2201858888067 |
| | 87.0458921833145 | 0.2201858888067 |
| | 88.0464196796744 | 0.21471857096257 |
| | 89.0469471760343 | 0.211787353496557 |
| | 90.0474746723943 | 0.207670784766858 |
| | 91.0480021687542 | 0.208233596897871 |
| | 92.0485296651141 | 0.21039984930418 |
| | 93.0490571614741 | 0.212694744483292 |
| | 94.049584657834 | 0.214314265105188 |
| | 95.0501121541939 | 0.216595377130072 |
| | 96.0506396505539 | 0.217250076955944 |
| | 97.0511671469138 | 0.218984457196415 |
| | 98.0516946432737 | 0.218757035151638 |
| | 99.0522221396337 | 0.221164492756952 |
| | 100.052749635994 | 0.220415608043849 |
| | 101.053277132354 | 0.221911080277685 |
| | 102.053804628713 | 0.226057512508212 |
| | 103.054332125073 | 0.227272727272727 |
| | 104.054859621433 | 0.228253628415351 |
| | 105.055387117793 | 0.227759732055482 |
| | 106.055914614153 | 0.228501725191471 |
| | 107.056442110513 | 0.230704732675724 |
| | 108.056969606873 | 0.232163449831616 |
| | 109.057497103233 | 0.231200926227964 |

| | | | | |
|---|---|---|---|---|
| | 110.058024599593 | 0.232655048999113 | | |
| | 111.058552095953 | 0.235078586951028 | | |
| | 112.059079592313 | 0.236100837556339 | | |
| | 113.059607088673 | 0.249564682045604 | | |
| | 114.060134585033 | 0.256849079055578 | | |
| | 115.060662081393 | 0.257637016038997 | | |
| | 116.061189577753 | 0.252034163844949 | | |
| | 117.061717074113 | 0.247414509985895 | | |
| | 118.062244570472 | 0.240842242611081 | | |
| | 119.062772066832 | 0.236663649687352 | | |
| | 120.063299563192 | 0.231777521513207 | | |
| | 121.063827059552 | 0.227169353616011 | | |
| | 122.064354555912 | 0.216974413871366 | | |
| | 123.064882052272 | 0.214314265105188 | | |
| | 124.065409548632 | 0.212357057204684 | | |
| | 125.065937044992 | 0.207634029688914 | | |
| | 126.066464541352 | 0.202171306229526 | | |
| | 127.066992037712 | 0.199240088763513 | | |
| | 128.067519534072 | 0.196993434624202 | | |
| | 129.068047030432 | 0.193763582149896 | | |
| | 130.068574526792 | 0.193083613207937 | | |
| | 131.069102023152 | 0.19737706575024 | | |
| | 132.069629519512 | 0.200310580408625 | | |
| | 133.070157015872 | 0.200308283216253 | | |
| Fragmention | 1 | 0.390019097 | 1 Ma | Zaffos et al. (2017) |
| | 2 | 0.451370613 | | |
| | 3 | 0.451276905 | | |
| | 4 | 0.450635508 | | |
| | 5 | 0.462296319 | | |
| | 6 | 0.455377551 | | |
| | 7 | 0.456101011 | | |
| | 8 | 0.457427958 | | |
| | 9 | 0.471878061 | | |
| | 10 | 0.471818565 | | |
| | 11 | 0.472092834 | | |
| | 12 | 0.491021922 | | |
| | 13 | 0.503465119 | | |
| | 14 | 0.513732777 | | |
| | 15 | 0.514107911 | | |
| | 16 | 0.504454352 | | |
| | 17 | 0.504431354 | | |
| | 18 | 0.504486085 | | |
| | 19 | 0.504452044 | | |

| | | |
|---|---|---|
| | 20 | 0.499900375 |
| | 21 | 0.499437412 |
| | 22 | 0.499518212 |
| | 23 | 0.499468683 |
| | 24 | 0.499342603 |
| | 25 | 0.498899203 |
| | 26 | 0.498704412 |
| | 27 | 0.498671438 |
| | 28 | 0.511151784 |
| | 29 | 0.511411975 |
| | 30 | 0.495216833 |
| | 31 | 0.49908992 |
| | 32 | 0.50056441 |
| | 33 | 0.501669088 |
| | 34 | 0.502249921 |
| | 35 | 0.503005105 |
| | 36 | 0.511397517 |
| | 37 | 0.514286919 |
| | 38 | 0.515820595 |
| | 39 | 0.517600388 |
| | 40 | 0.518784635 |
| | 41 | 0.520547858 |
| | 42 | 0.521662459 |
| | 43 | 0.521948138 |
| | 44 | 0.522097081 |
| | 45 | 0.522220792 |
| | 46 | 0.522462222 |
| | 47 | 0.522978858 |
| | 48 | 0.523753689 |
| | 49 | 0.521844409 |
| | 50 | 0.521542365 |
| | 51 | 0.527391223 |
| | 52 | 0.527552801 |
| | 53 | 0.527676321 |
| | 54 | 0.52768445 |
| | 55 | 0.527654922 |
| | 56 | 0.536111742 |
| | 57 | 0.535040363 |
| | 58 | 0.532740141 |
| | 59 | 0.532468583 |
| | 60 | 0.531692225 |
| | 61 | 0.53479689 |
| | 62 | 0.53519875 |

| | | |
|---|---|---|
| | 63 | 0.535512164 |
| | 64 | 0.535821435 |
| | 65 | 0.53376153 |
| | 66 | 0.533163123 |
| | 67 | 0.533589932 |
| | 68 | 0.537438526 |
| | 69 | 0.538814495 |
| | 70 | 0.539581963 |
| | 71 | 0.543123084 |
| | 72 | 0.543398635 |
| | 73 | 0.543349118 |
| | 74 | 0.543382893 |
| | 75 | 0.544245874 |
| | 76 | 0.543441653 |
| | 77 | 0.53878069 |
| | 78 | 0.535073748 |
| | 79 | 0.534373582 |
| | 80 | 0.533803129 |
| | 81 | 0.531563144 |
| | 82 | 0.529626462 |
| | 83 | 0.529229458 |
| | 84 | 0.528635507 |
| | 85 | 0.52573418 |
| | 86 | 0.516348709 |
| | 87 | 0.516673003 |
| | 88 | 0.515809698 |
| | 89 | 0.514559122 |
| | 90 | 0.517379917 |
| | 91 | 0.521695946 |
| | 92 | 0.520482294 |
| | 93 | 0.514892456 |
| | 94 | 0.512957032 |
| | 95 | 0.51137412 |
| | 96 | 0.508706724 |
| | 97 | 0.50852325 |
| | 98 | 0.507816863 |
| | 99 | 0.507534766 |
| | 100 | 0.507375269 |
| | 101 | 0.50722461 |
| | 102 | 0.507149371 |
| | 103 | 0.507064931 |
| | 104 | 0.507104446 |
| | 105 | 0.507036838 |

| | | | | |
|---|---|---|---|---|
| | 106 | 0.505580138 | | |
| | 107 | 0.499793016 | | |
| | 108 | 0.498585868 | | |
| | 109 | 0.497576577 | | |
| | 110 | 0.496779453 | | |
| | 111 | 0.486903642 | | |
| | 112 | 0.485679678 | | |
| | 113 | 0.461759614 | | |
| | 114 | 0.46023643 | | |
| | 115 | 0.449990884 | | |
| | 116 | 0.452444471 | | |
| | 117 | 0.444765012 | | |
| | 118 | 0.437508142 | | |
| | 119 | 0.435686482 | | |
| | 120 | 0.436840605 | | |
| | 121 | 0.457002974 | | |
| | 122 | 0.451363839 | | |
| | 123 | 0.451851659 | | |
| | 124 | 0.451873317 | | |
| | 125 | 0.451854357 | | |
| | 126 | 0.453521917 | | |
| | 127 | 0.438506176 | | |
| | 128 | 0.439523824 | | |
| | 129 | 0.439967133 | | |
| | 130 | 0.440364274 | | |
| | 131 | 0.420199518 | | |
| | 132 | 0.409684229 | | |
| | 133 | 0.409752889 | | |
| Gymnospermae | 1.00255721561398 | 0.708843537414966 | 1 Ma | This study |
| | 2.00511443122795 | 0.704761904761905 | | |
| | 3.00767164684193 | 0.715646258503401 | | |
| | 4.01022886245591 | 0.718367346938776 | | |
| | 5.01278607806989 | 0.717006802721088 | | |
| | 6.01534329368386 | 0.71156462585034 | | |
| | 7.01790050929784 | 0.710204081632653 | | |
| | 8.02045772491182 | 0.71156462585034 | | |
| | 9.0230149405258 | 0.729251700680272 | | |
| | 10.0255721561398 | 0.751020408163265 | | |
| | 11.0281293717538 | 0.764625850340136 | | |
| | 12.0306865873677 | 0.794557823129252 | | |
| | 13.0332438029817 | 0.806802721088435 | | |
| | 14.0358010185957 | 0.820408163265306 | | |
| | 15.0383582342097 | 0.828571428571429 | | |

| | | |
|---|---|---|
| | 16.0409154498236 | 0.834013605442177 |
| | 17.0434726654376 | 0.839455782312925 |
| | 18.0460298810516 | 0.853061224489796 |
| | 19.0485870966656 | 0.853061224489796 |
| | 20.0511443122796 | 0.85578231292517 |
| | 21.0537015278935 | 0.863945578231293 |
| | 22.0562587435075 | 0.869387755102041 |
| | 23.0588159591215 | 0.884353741496599 |
| | 24.0613731747355 | 0.88843537414966 |
| | 25.0639303903494 | 0.884353741496599 |
| | 26.0664876059634 | 0.882993197278912 |
| | 27.0690448215774 | 0.88843537414966 |
| | 28.0716020371914 | 0.892517006802721 |
| | 29.0741592528053 | 0.904761904761905 |
| | 30.0767164684193 | 0.925170068027211 |
| | 31.0792736840333 | 0.930612244897959 |
| | 32.0818308996473 | 0.955102040816327 |
| | 33.0843881152613 | 0.980952380952381 |
| | 34.0869453308752 | 0.993197278911565 |
| | 35.0895025464892 | 0.994557823129252 |
| | 36.0920597621032 | 1 |
| | 37.0946169777172 | 0.998639455782313 |
| | 38.0971741933311 | 0.997278911564626 |
| | 39.0997314089451 | 0.998639455782313 |
| | 40.1022886245591 | 0.993197278911565 |
| | 41.1048458401731 | 0.993197278911565 |
| | 42.1074030557871 | 0.97687074829932 |
| | 43.109960271401 | 0.97687074829932 |
| | 44.112517487015 | 0.974149659863946 |
| | 45.115074702629 | 0.97687074829932 |
| | 46.117631918243 | 0.971428571428571 |
| | 47.1201891338569 | 0.968707482993197 |
| | 48.1227463494709 | 0.964625850340136 |
| | 49.1253035650849 | 0.949659863945578 |
| | 50.1278607806989 | 0.942857142857143 |
| | 51.1304179963129 | 0.938775510204082 |
| | 52.1329752119268 | 0.933333333333333 |
| | 53.1355324275408 | 0.929251700680272 |
| | 54.1380896431548 | 0.925170068027211 |
| | 55.1406468587688 | 0.922448979591837 |
| | 56.1432040743827 | 0.908843537414966 |
| | 57.1457612899967 | 0.885714285714286 |
| | 58.1483185056107 | 0.863945578231293 |

| | | |
|---|---|---|
| | 59.1508757212247 | 0.858503401360544 |
| | 60.1534329368386 | 0.859863945578231 |
| | 61.1559901524526 | 0.851700680272109 |
| | 62.1585473680666 | 0.850340136054422 |
| | 63.1611045836806 | 0.847619047619048 |
| | 64.1636617992946 | 0.846258503401361 |
| | 65.1662190149085 | 0.842176870748299 |
| | 66.1687762305225 | 0.840816326530612 |
| | 67.1713334461365 | 0.85578231292517 |
| | 68.1738906617505 | 0.850340136054422 |
| | 69.1764478773645 | 0.848979591836735 |
| | 70.1790050929784 | 0.858503401360544 |
| | 71.1815623085924 | 0.857142857142857 |
| | 72.1841195242064 | 0.870748299319728 |
| | 73.1866767398204 | 0.880272108843538 |
| | 74.1892339554343 | 0.88843537414966 |
| | 75.1917911710483 | 0.889795918367347 |
| | 76.1943483866623 | 0.892517006802721 |
| | 77.1969056022763 | 0.893877551020408 |
| | 78.1994628178903 | 0.88843537414966 |
| | 79.2020200335042 | 0.882993197278912 |
| | 80.2045772491182 | 0.882993197278912 |
| | 81.2071344647322 | 0.882993197278912 |
| | 82.2096916803462 | 0.87891156462585 |
| | 83.2122488959601 | 0.868027210884354 |
| | 84.2148061115741 | 0.862585034013606 |
| | 85.2173633271881 | 0.861224489795918 |
| | 86.2199205428021 | 0.850340136054422 |
| | 87.222477758416 | 0.846258503401361 |
| | 88.22503497 | 0.844897959183673 |
| | 89.227592189644 | 0.836734693877551 |
| | 90.230149405258 | 0.842176870748299 |
| | 91.232706620872 | 0.848979591836735 |
| | 92.2352638364859 | 0.853061224489796 |
| | 93.2378210520999 | 0.853061224489796 |
| | 94.2403782677139 | 0.85578231292517 |
| | 95.2429354833279 | 0.86530612244898 |
| | 96.2454926989418 | 0.862585034013606 |
| | 97.2480499145558 | 0.862585034013606 |
| | 98.2506071301698 | 0.85578231292517 |
| | 99.2531643457838 | 0.851700680272109 |
| | 100.255721561398 | 0.850340136054422 |
| | 101.258278777012 | 0.828571428571429 |

| | | | | |
|---|---|---|---|---|
| | 102.260835992626 | 0.814965986394558 | | |
| | 103.26339320824 | 0.814965986394558 | | |
| | 104.265950423854 | 0.83265306122449 | | |
| | 105.268507639468 | 0.842176870748299 | | |
| | 106.271064855082 | 0.853061224489796 | | |
| | 107.273622070696 | 0.86530612244898 | | |
| | 108.27617928631 | 0.87891156462585 | | |
| | 109.278736501924 | 0.887074829931973 | | |
| | 110.281293717538 | 0.902040816326531 | | |
| | 111.283850933151 | 0.922448979591837 | | |
| | 112.286408148765 | 0.938775510204082 | | |
| | 113.288965364379 | 0.94421768707483 | | |
| | 114.291522579993 | 0.948299319727891 | | |
| | 115.294079795607 | 0.952380952380952 | | |
| | 116.296637011221 | 0.952380952380952 | | |
| | 117.299194226835 | 0.948299319727891 | | |
| | 118.301751442449 | 0.949659863945578 | | |
| | 119.304308658063 | 0.949659863945578 | | |
| | 120.306865873677 | 0.951020408163265 | | |
| | 121.309423089291 | 0.956462585034014 | | |
| | 122.311980304905 | 0.953741496598639 | | |
| | 123.314537520519 | 0.951020408163265 | | |
| | 124.317094736133 | 0.949659863945578 | | |
| | 125.319651951747 | 0.949659863945578 | | |
| | 126.322209167361 | 0.94421768707483 | | |
| | 127.324766382975 | 0.941496598639456 | | |
| | 128.327323598589 | 0.936054421768708 | | |
| | 129.329880814203 | 0.927891156462585 | | |
| | 130.332438029817 | 0.922448979591837 | | |
| | 131.334995245431 | 0.915646258503401 | | |
| | 132.337552461045 | 0.912925170068027 | | |
| | 133.340109676659 | 0.907482993197279 | | |
| Insect | 1.00219379106929 | 0.977571196694703 | 1 Ma | Peris & Condamine (2024) |
| | 2.00438758213858 | 0.976685849195809 | | |
| | 3.00658137320787 | 0.977128522945256 | | |
| | 4.00877516427716 | 0.977866312527667 | | |
| | 5.01096895534645 | 0.979046775859525 | | |
| | 6.01316274641574 | 0.979637007525454 | | |
| | 7.01535653748503 | 0.979470497216029 | | |
| | 8.01755032855431 | 0.97875166002656 | | |
| | 9.0197441196236 | 0.978308986277114 | | |
| | 10.0219379106929 | 0.97739656393116 | | |
| | 11.0241317017622 | 0.976685849195809 | | |

| | | |
|---|---|---|
| | 12.0263254928315 | 0.976063127713407 |
| | 13.0285192839008 | 0.975582549178075 |
| | 14.0307130749701 | 0.975062712114505 |
| | 15.0329068660393 | 0.974915154198023 |
| | 16.0351006571086 | 0.975062712114505 |
| | 17.0372944481779 | 0.974574011125055 |
| | 18.0394882392472 | 0.973144459200236 |
| | 19.0416820303165 | 0.972650343241374 |
| | 20.0438758213858 | 0.971614730341292 |
| | 21.0460696124551 | 0.969932028596454 |
| | 22.0482634035244 | 0.967920367459824 |
| | 23.0504571945937 | 0.967537258373912 |
| | 24.0526509856629 | 0.968405007222215 |
| | 25.0548447767322 | 0.969375640489351 |
| | 26.0570385678015 | 0.969821021723503 |
| | 27.0592323588708 | 0.970340858787074 |
| | 28.0614261499401 | 0.97004574295411 |
| | 29.0636199410094 | 0.971078648369485 |
| | 30.0658137320787 | 0.975550059361602 |
| | 31.068007523148 | 0.982860268068061 |
| | 32.0702013142173 | 0.987804135141392 |
| | 33.0723951052865 | 0.991391552376969 |
| | 34.0745888963558 | 0.998468917398703 |
| | 35.0767826874251 | 1 |
| | 36.0789764784944 | 0.998376862918696 |
| | 37.0811702695637 | 0.99378632259951 |
| | 38.083364060633 | 0.983518186851642 |
| | 39.0855578517023 | 0.951112302404382 |
| | 40.0877516427716 | 0.936903422666791 |
| | 41.0899454338409 | 0.924004965526951 |
| | 42.0921392249101 | 0.904469109630117 |
| | 43.0943330159794 | 0.870709482829809 |
| | 44.0965268070487 | 0.833589867509236 |
| | 45.098720598118 | 0.796987381767527 |
| | 46.1009143891873 | 0.762016155561241 |
| | 47.1031081802566 | 0.740796244177216 |
| | 48.1053019713259 | 0.730352121923451 |
| | 49.1074957623952 | 0.724583825757114 |
| | 50.1096895534645 | 0.720488755139144 |
| | 51.1118833445338 | 0.714513336392791 |
| | 52.114077135603 | 0.707835325365206 |
| | 53.1162709266723 | 0.702096811530636 |
| | 54.1184647177416 | 0.694855643684183 |

| | | |
|---|---|---|
| | 55.1206585088109 | 0.679485253007677 |
| | 56.1228522998802 | 0.662949290165197 |
| | 57.1250460909495 | 0.645048755030845 |
| | 58.1272398820188 | 0.630439167556752 |
| | 59.1294336730881 | 0.615932464501491 |
| | 60.1316274641574 | 0.599998917006118 |
| | 61.1338212552266 | 0.592079524240787 |
| | 62.1360150462959 | 0.58454324056137 |
| | 63.1382088373652 | 0.578971237036225 |
| | 64.1404026284345 | 0.57616222165365 |
| | 65.1425964195038 | 0.575447445691241 |
| | 66.1447902105731 | 0.573992172661715 |
| | 67.1469840016424 | 0.572752144666323 |
| | 68.1491777927117 | 0.571823477412132 |
| | 69.151371583781 | 0.572563974479249 |
| | 70.1535653748502 | 0.575960514043046 |
| | 71.1557591659195 | 0.578945515931517 |
| | 72.1579529569888 | 0.580982898172854 |
| | 73.1601467480581 | 0.582853770104766 |
| | 74.1623405391274 | 0.584034233436624 |
| | 75.1645343301967 | 0.585214696768482 |
| | 76.166728121266 | 0.586923119618028 |
| | 77.1689219123353 | 0.588313413014608 |
| | 78.1711157034046 | 0.590442849735953 |
| | 79.1733094944739 | 0.593758164758567 |
| | 80.1755032855431 | 0.598033283109492 |
| | 81.1776970766124 | 0.601358074328578 |
| | 82.1798908676817 | 0.607130431721974 |
| | 83.182084658751 | 0.613777306675439 |
| | 84.1842784498203 | 0.620210290337122 |
| | 85.1864722408896 | 0.626483532401146 |
| | 86.1886660319589 | 0.633217046865206 |
| | 87.1908598230282 | 0.636797695389018 |
| | 88.1930536140975 | 0.641388235708204 |
| | 89.1952474051668 | 0.64804458685814 |
| | 90.197441196236 | 0.652398222265542 |
| | 91.1996349873053 | 0.655945027230527 |
| | 92.2018287783746 | 0.659344274279031 |
| | 93.2040225694439 | 0.659688124836705 |
| | 94.2062163605132 | 0.660240451716748 |
| | 95.2084101515825 | 0.661687602292157 |
| | 96.2106039426518 | 0.662872126851073 |
| | 97.2127977337211 | 0.661189425106235 |

| | | | | |
|---|---|---|---|---|
| | 98.2149915247903 | 0.66067635675443 | | |
| | 99.2171853158596 | 0.658819022246048 | | |
| | 100.219379106929 | 0.6553778091846 | | |
| | 101.22157289798 | 0.653260556144434 | | |
| | 102.223766689068 | 0.649273784914708 | | |
| | 103.225960480137 | 0.647221511507487 | | |
| | 104.228154271206 | 0.643876414153106 | | |
| | 105.230348062275 | 0.636356375381925 | | |
| | 106.232541853345 | 0.628677948755437 | | |
| | 107.234735644414 | 0.61374210937426 | | |
| | 108.236929435483 | 0.601467727459175 | | |
| | 109.239123226553 | 0.592297476759628 | | |
| | 110.241317017622 | 0.586242187214445 | | |
| | 111.243510808691 | 0.588241664669897 | | |
| | 112.24570459976 | 0.611609965168209 | | |
| | 113.24789839083 | 0.624666810163356 | | |
| | 114.250092181899 | 0.636055844579548 | | |
| | 115.252285972968 | 0.644925564476718 | | |
| | 116.254479764038 | 0.654423420825701 | | |
| | 117.256673555107 | 0.659605546553169 | | |
| | 118.258867346176 | 0.666469020283122 | | |
| | 119.261061137245 | 0.669443192232768 | | |
| | 120.263254928315 | 0.6711231864929 | | |
| | 121.265448719384 | 0.671550969076463 | | |
| | 122.267642510453 | 0.674269283721384 | | |
| | 123.269836301523 | 0.667067374403169 | | |
| | 124.272030092592 | 0.660199839446157 | | |
| | 125.274223883661 | 0.651078323471321 | | |
| | 126.27641767473 | 0.639498411383349 | | |
| | 127.2786115 | 0.624665456421003 | | |
| | 128.280805256869 | 0.612986721141259 | | |
| | 129.282999047938 | 0.597673187643581 | | |
| | 130.285192839008 | 0.574214186407615 | | |
| | 131.287386630077 | 0.54908602085034 | | |
| | 132.289580421146 | 0.530251403492385 | | |
| | 133.291774212215 | 0.51513010140884 | | |
| Pollinator | 1.00242158447329 | 1 | 1 Ma | Peris & Condamine (2024) |
| | 2.00484316894658 | 0.997627520759193 | | |
| | 3.00726475341986 | 0.997034400948992 | | |
| | 4.00968633789315 | 0.99644128113879 | | |
| | 5.01210792236644 | 0.99644128113879 | | |
| | 6.01452950683973 | 0.99644128113879 | | |
| | 7.01695109131302 | 0.994661921708185 | | |

| | | |
|---|---|---|
| | 8.0193726757863 | 0.994661921708185 |
| | 9.02179426025959 | 0.994068801897983 |
| | 10.0242158447329 | 0.99288256227758 |
| | 11.0266374292062 | 0.99288256227758 |
| | 12.0290590136795 | 0.992289442467378 |
| | 13.0314805981527 | 0.992289442467378 |
| | 14.033902182626 | 0.991696322657177 |
| | 15.0363237670993 | 0.991696322657177 |
| | 16.0387453515726 | 0.991696322657177 |
| | 17.0411669360459 | 0.989916963226572 |
| | 18.0435885205192 | 0.98576512455516 |
| | 19.0460101049925 | 0.984960043433476 |
| | 20.0484316894658 | 0.983392645314354 |
| | 21.050853273939 | 0.982565252475671 |
| | 22.0532748584123 | 0.980427046263345 |
| | 23.0556964428856 | 0.979833926453144 |
| | 24.0581180273589 | 0.980427046263345 |
| | 25.0605396118322 | 0.979833926453144 |
| | 26.0629611963055 | 0.97864768683274 |
| | 27.0653827807788 | 0.97864768683274 |
| | 28.0678043652521 | 0.978054567022539 |
| | 29.0702259497254 | 0.978054567022539 |
| | 30.0726475341986 | 0.97779612296731 |
| | 31.0750691186719 | 0.979833926453144 |
| | 32.0774907031452 | 0.981377153545518 |
| | 33.0799122876185 | 0.981613285883748 |
| | 34.0823338720918 | 0.981613285883748 |
| | 35.0847554565651 | 0.981020166073547 |
| | 36.0871770410384 | 0.979240806642942 |
| | 37.0895986255117 | 0.977461447212337 |
| | 38.0920202099849 | 0.970937129300119 |
| | 39.0944417944582 | 0.945907101447659 |
| | 40.0968633789315 | 0.937276185588862 |
| | 41.0992849634048 | 0.929418742586002 |
| | 42.1017065478781 | 0.915557588400882 |
| | 43.1041281323514 | 0.88454244246366 |
| | 44.1065497168247 | 0.851147380046631 |
| | 45.108971301298 | 0.832329306068415 |
| | 46.1113928857712 | 0.802730954160577 |
| | 47.1138144702445 | 0.783717652658627 |
| | 48.1162360547178 | 0.77128630767114 |
| | 49.1186576391911 | 0.762298404340373 |
| | 50.1210792236644 | 0.756820877817319 |

| | | |
|---|---|---|
| | 51.1235008081377 | 0.746762012070639 |
| | 52.125922392611 | 0.743772241992882 |
| | 53.1283439770843 | 0.736662241509462 |
| | 54.1307655615576 | 0.730130486358244 |
| | 55.1331871460308 | 0.716345563872868 |
| | 56.1356087305041 | 0.702190638747271 |
| | 57.1380303149774 | 0.686154092154828 |
| | 58.1404518994507 | 0.675509543836946 |
| | 59.142873483924 | 0.664757155553572 |
| | 60.1452950683973 | 0.647686832740213 |
| | 61.1477166528706 | 0.639976275207592 |
| | 62.1501382373439 | 0.628608455397018 |
| | 63.1525598218171 | 0.619700502385494 |
| | 64.1549814062904 | 0.617316867286189 |
| | 65.1574029907637 | 0.616119471807286 |
| | 66.159824575237 | 0.611956477277376 |
| | 67.1622461597103 | 0.610165961988271 |
| | 68.1646677441836 | 0.608540925266904 |
| | 69.1670893286569 | 0.606761565836299 |
| | 70.1695109131302 | 0.604982206405694 |
| | 71.1719324976034 | 0.603795966785291 |
| | 72.1743540820767 | 0.603795966785291 |
| | 73.17677567 | 0.603795966785291 |
| | 74.1791972510233 | 0.603202846975089 |
| | 75.1816188354966 | 0.603202846975089 |
| | 76.1840404199699 | 0.603202846975089 |
| | 77.1864620044432 | 0.603202846975089 |
| | 78.1888835889165 | 0.603795966785291 |
| | 79.1913051733898 | 0.603795966785291 |
| | 80.193726757863 | 0.603202846975089 |
| | 81.1961483423363 | 0.603202846975089 |
| | 82.1985699268096 | 0.603202846975089 |
| | 83.2009915112829 | 0.602609727164887 |
| | 84.2034130957562 | 0.602953699468609 |
| | 85.2058346802295 | 0.602016607354686 |
| | 86.2082562647028 | 0.604389086595492 |
| | 87.2106778491761 | 0.606168446026097 |
| | 88.2130994336493 | 0.607354685646501 |
| | 89.2155210181226 | 0.606361814240082 |
| | 90.2179426025959 | 0.607354685646501 |
| | 91.2203641870692 | 0.608540925266904 |
| | 92.2227857715425 | 0.608540925266904 |
| | 93.2252073560158 | 0.607354685646501 |

| | | | | |
|---|---|---|---|---|
| | 94.2276289404891 | 0.606761565836299 | | |
| | 95.2300505249624 | 0.606635132773309 | | |
| | 96.2324721094356 | 0.605097483610185 | | |
| | 97.2348936939089 | 0.600934489080274 | | |
| | 98.2373152783822 | 0.598457888493476 | | |
| | 99.2397368628555 | 0.594899169632266 | | |
| | 100.242158447329 | 0.593119810201661 | | |
| | 101.244580031802 | 0.591340450771056 | | |
| | 102.247001616275 | 0.588374851720047 | | |
| | 103.249423200749 | 0.586632678484439 | | |
| | 104.251844785222 | 0.58306280376473 | | |
| | 105.254266369695 | 0.574154850753206 | | |
| | 106.256687954169 | 0.567626814221488 | | |
| | 107.259109538642 | 0.563456382452578 | | |
| | 108.261531123115 | 0.556902315584362 | | |
| | 109.263952707588 | 0.554537273582555 | | |
| | 110.266374292062 | 0.548001799811838 | | |
| | 111.268795876535 | 0.542111506524318 | | |
| | 112.271217461008 | 0.538022884384401 | | |
| | 113.273639045482 | 0.530693485350498 | | |
| | 114.276060629955 | 0.530249110320285 | | |
| | 115.278482214428 | 0.529159554806873 | | |
| | 116.280903798901 | 0.53202846975089 | | |
| | 117.283325383375 | 0.532978577033062 | | |
| | 118.285746967848 | 0.536773428232503 | | |
| | 119.288168552321 | 0.53202846975089 | | |
| | 120.290590136795 | 0.529503527110595 | | |
| | 121.293011721268 | 0.526199533685115 | | |
| | 122.295433305741 | 0.523131672597865 | | |
| | 123.297854890214 | 0.516828612545878 | | |
| | 124.300276474688 | 0.513048635824436 | | |
| | 125.302698059161 | 0.504120230405664 | | |
| | 126.305119643634 | 0.491696322657177 | | |
| | 127.307541228108 | 0.478186578014778 | | |
| | 128.309962812581 | 0.470102299222437 | | |
| | 129.312384397054 | 0.463732304019456 | | |
| | 130.314805981527 | 0.44147636631377 | | |
| | 131.317227566001 | 0.425860023724792 | | |
| | 132.319649150474 | 0.414631652145458 | | |
| | 133.322070734947 | 0.40539646061796 | | |
| Pteridophyta | 1.00024094985104 | 0.757624001797793 | 1 Ma | This study |
| | 2.00048189970207 | 0.763856138607281 | | |
| | 3.00072284955311 | 0.746489081891354 | | |

| | | |
|---|---|---|
| | 4.00096379940415 | 0.727871799660065 |
| | 5.00120474925518 | 0.768477225322289 |
| | 6.00144569910622 | 0.780121730818534 |
| | 7.00168664895726 | 0.775294910790871 |
| | 8.00192759880829 | 0.78384075608575 |
| | 9.00216854865933 | 0.780254666189787 |
| | 10.0024094985104 | 0.769205204736297 |
| | 11.0026504483614 | 0.763922606292907 |
| | 12.0028913982124 | 0.762637564370788 |
| | 13.0031323480635 | 0.757842395621995 |
| | 14.0033732979145 | 0.756386436793979 |
| | 15.0036142477656 | 0.755136211278617 |
| | 16.0038551976166 | 0.751436176778723 |
| | 17.0040961474676 | 0.745242021503879 |
| | 18.0043370973187 | 0.740190477396239 |
| | 19.0045780471697 | 0.737693191493402 |
| | 20.0048189970207 | 0.745007802040242 |
| | 21.0050599468718 | 0.747634858186445 |
| | 22.0053008967228 | 0.753677087322713 |
| | 23.0055418465738 | 0.753958783704656 |
| | 24.0057827964249 | 0.753885985763255 |
| | 25.0060237462759 | 0.758807759627528 |
| | 26.006264696127 | 0.763805496561089 |
| | 27.006505645978 | 0.766302782463926 |
| | 28.006746595829 | 0.777108529070117 |
| | 29.0069875456801 | 0.790750230263054 |
| | 30.0072284955311 | 0.805540872878969 |
| | 31.0074694453821 | 0.829070433590869 |
| | 32.0077103952332 | 0.839746409953694 |
| | 33.0079513450842 | 0.846298224679768 |
| | 34.0081922949352 | 0.850045736097967 |
| | 35.0084332447863 | 0.848903124930763 |
| | 36.0086741946373 | 0.853676137784347 |
| | 37.0089151444884 | 0.860819831425289 |
| | 38.0091560943394 | 0.861285105224677 |
| | 39.0093970441904 | 0.860278594556613 |
| | 40.0096379940415 | 0.860028549453541 |
| | 41.0098789438925 | 0.861275609841016 |
| | 42.0101198937435 | 0.854568703848479 |
| | 43.0103608435946 | 0.850153350446125 |
| | 44.0106017934456 | 0.840569343204312 |
| | 45.0108427432966 | 0.83779352604742 |
| | 46.0110836931477 | 0.83073212573154 |

| | | |
|---|---|---|
| | 47.0113246429987 | 0.819641517615519 |
| | 48.0115655928498 | 0.809807465270634 |
| | 49.0118065427008 | 0.806395457408457 |
| | 50.0120474925518 | 0.804053262772082 |
| | 51.0122884424029 | 0.808734486916944 |
| | 52.0125293922539 | 0.807481096273695 |
| | 53.0127703421049 | 0.806227705630446 |
| | 54.013011291956 | 0.804974314987197 |
| | 55.013252241807 | 0.809069990472965 |
| | 56.0134931916581 | 0.81818872391539 |
| | 57.0137341415091 | 0.821037339013683 |
| | 58.0139750913601 | 0.822648389108162 |
| | 59.0142160412112 | 0.821584906138133 |
| | 60.0144569910622 | 0.834501793044948 |
| | 61.0146979409132 | 0.855246041216295 |
| | 62.0149388907643 | 0.871479982148679 |
| | 63.0151798406153 | 0.878509731185689 |
| | 64.0154207904664 | 0.882051509291233 |
| | 65.0156617403174 | 0.892882576920521 |
| | 66.0159026901684 | 0.896411694514517 |
| | 67.0161436400195 | 0.919719696274328 |
| | 68.0163845898705 | 0.927410957039719 |
| | 69.0166255397215 | 0.936200517181897 |
| | 70.0168664895726 | 0.93428244968238 |
| | 71.0171074394236 | 0.938919362036823 |
| | 72.0173483892746 | 0.937441247313598 |
| | 73.0175893391257 | 0.942764992419519 |
| | 74.0178302889767 | 0.942670038582909 |
| | 75.0180712388278 | 0.93814390570451 |
| | 76.0183121886788 | 0.92661017968431 |
| | 77.0185531385298 | 0.916659017607607 |
| | 78.0187940883809 | 0.919425339380838 |
| | 79.0190350382319 | 0.919915934203322 |
| | 80.0192759880829 | 0.919912769075435 |
| | 81.019516937934 | 0.918178278993363 |
| | 82.019757887785 | 0.915943698705146 |
| | 83.0199988376361 | 0.912927331828843 |
| | 84.0202397874871 | 0.909977432638166 |
| | 85.0204807373381 | 0.902393786220932 |
| | 86.0207216871892 | 0.892227395447913 |
| | 87.0209626370402 | 0.898510174303593 |
| | 88.0212035868912 | 0.912522195459308 |
| | 89.0214445367423 | 0.916538742747901 |

| | | |
|---|---|---|
| | 90.0216854865933 | 0.923631794342651 |
| | 91.0219264364443 | 0.925166881367842 |
| | 92.0221673862954 | 0.932753692912962 |
| | 93.0224083361464 | 0.939286516871714 |
| | 94.0226492859975 | 0.936061251554869 |
| | 95.0228902358485 | 0.934867998341473 |
| | 96.0231311856995 | 0.933883643568619 |
| | 97.0233721355506 | 0.933611442570337 |
| | 98.0236130854016 | 0.936099233089513 |
| | 99.0238540352526 | 0.932354886799201 |
| | 100.024094985104 | 0.936418911006099 |
| | 101.024335934955 | 0.926154401268583 |
| | 102.024576884806 | 0.918551764084028 |
| | 103.024817834657 | 0.928863750739849 |
| | 104.025058784508 | 0.933918459975375 |
| | 105.025299734359 | 0.940755136211279 |
| | 106.02554068421 | 0.95132982848172 |
| | 107.025781634061 | 0.956083850567982 |
| | 108.026022583912 | 0.96198997920511 |
| | 109.026263533763 | 0.968234776526146 |
| | 110.026504483614 | 0.973783245712043 |
| | 111.026745433465 | 0.980373041972761 |
| | 112.026986383316 | 0.993929284712749 |
| | 113.027227333167 | 1 |
| | 114.027468283018 | 0.9950117584501 |
| | 115.027709232869 | 0.985019449710866 |
| | 116.02795018272 | 0.979293733363297 |
| | 117.028191132571 | 0.974666316392514 |
| | 118.028432082422 | 0.970921970102202 |
| | 119.028673032273 | 0.963306672406098 |
| | 120.028913982124 | 0.955172293736528 |
| | 121.029154931975 | 0.945452185995575 |
| | 122.029395881826 | 0.932190300149078 |
| | 123.029636831678 | 0.926806417613304 |
| | 124.029877781529 | 0.911268804816059 |
| | 125.03011873138 | 0.903849745048949 |
| | 126.030359681231 | 0.896357887340438 |
| | 127.030600631082 | 0.881766647781404 |
| | 128.030841580933 | 0.867552058440921 |
| | 129.031082530784 | 0.847576936346113 |
| | 130.031323480635 | 0.833852941828115 |
| | 131.031564430486 | 0.818885052050528 |
| | 132.031805380337 | 0.80140405073067 |

| | | | | |
|---|---|---|---|---|
| | 133.032046330188 | 0.77561142357957 | | |
| Sediment | 0 | 4.39777227186223 | variable | This study |
| | 3 | 4.14007652981768 | | |
| | 11 | 4.08255388673005 | | |
| | 15 | 4.05963226442009 | | |
| | 20 | 4.6571134113196 | | |
| | 26 | 4.51977652840603 | | |
| | 31 | 5.14822968151906 | | |
| | 36 | 4.54694570471865 | | |
| | 40 | 4.60053673118484 | | |
| | 45 | 4.09807264308877 | | |
| | 52 | 4.24505595316465 | | |
| | 56 | 3.34451495638251 | | |
| | 61 | 3.12989494085987 | | |
| | 66 | 2.89735981925637 | | |
| | 69 | 3.2437257438989 | | |
| | 75 | 3.26983431173867 | | |
| | 81 | 3.56261608206661 | | |
| | 87 | 3.24369440320473 | | |
| | 92 | 3.33942266862923 | | |
| | 97 | 3.55328996054109 | | |
| | 103 | 4.41089230860165 | | |
| | 107 | 3.69563920688673 | | |
| | 111 | 3.2358614010072 | | |
| | 116 | 2.89046758323695 | | |
| | 122 | 2.92304158640879 | | |
| | 127 | 2.35077737809747 | | |
| | 131 | 2.46993916427186 | | |
| Temperature | 0 | 1.17548792447447 | 0.5 Ma | Peris & Condamine (2024) |
| | 0.5 | 1.62097067878976 | | |
| | 1 | 2.06676385228642 | | |
| | 1.5 | 2.51249642682119 | | |
| | 2 | 2.95584686243243 | | |
| | 2.5 | 3.39285306292583 | | |
| | 3 | 3.81708930840564 | | |
| | 3.5 | 4.218680857 | | |
| | 4 | 4.58938506569937 | | |
| | 4.5 | 4.92586486744603 | | |
| | 5 | 5.22567085078944 | | |
| | 5.5 | 5.48688662057616 | | |
| | 6 | 5.70755599016075 | | |
| | 6.5 | 5.88354579011277 | | |
| | 7 | 6.00978807348009 | | |

| | | |
|---|---|---|
| | 7.5 | 6.09213134545395 |
| | 8 | 6.14901338582887 |
| | 8.5 | 6.19869659121235 |
| | 9 | 6.25560353317631 |
| | 9.5 | 6.3329227978124 |
| | 10 | 6.44084512363611 |
| | 10.5 | 6.58706027395652 |
| | 11 | 6.77796897371783 |
| | 11.5 | 7.01708145835566 |
| | 12 | 7.30716328681904 |
| | 12.5 | 7.64536035408088 |
| | 13 | 8.02402545335558 |
| | 13.5 | 8.43082330880991 |
| | 14 | 8.84395088804806 |
| | 14.5 | 9.24091689 |
| | 15 | 9.60175945638438 |
| | 15.5 | 9.90813792596568 |
| | 16 | 10.1523614889028 |
| | 16.5 | 10.3341973168494 |
| | 17 | 10.4539531271709 |
| | 17.5 | 10.5125114699412 |
| | 18 | 10.5120964503801 |
| | 18.5 | 10.4622565582927 |
| | 19 | 10.3751704011242 |
| | 19.5 | 10.2657808986242 |
| | 20 | 10.152946809643 |
| | 20.5 | 10.0533517460733 |
| | 21 | 9.96543952006742 |
| | 21.5 | 9.8795092696787 |
| | 22 | 9.78680881397512 |
| | 22.5 | 9.6801064130819 |
| | 23 | 9.55230022188102 |
| | 23.5 | 9.39634573632495 |
| | 24 | 9.20533784700675 |
| | 24.5 | 8.97340839698132 |
| | 25 | 8.70049334920442 |
| | 25.5 | 8.38969268439382 |
| | 26 | 8.05924749444203 |
| | 26.5 | 7.73823674213118 |
| | 27 | 7.44811051950403 |
| | 27.5 | 7.19807707580706 |
| | 28 | 6.99606564012444 |
| | 28.5 | 6.84326825728201 |

| | | |
|---|---|---|
| | 29 | 6.73307911492559 |
| | 29.5 | 6.66041547080171 |
| | 30 | 6.6271849744661 |
| | 30.5 | 6.6369861959064 |
| | 31 | 6.69594676201703 |
| | 31.5 | 6.81247655119057 |
| | 32 | 6.99592038771653 |
| | 32.5 | 7.25754022443286 |
| | 33 | 7.6052554361432 |
| | 33.5 | 8.0295101006167 |
| | 34 | 8.51492402842034 |
| | 34.5 | 9.0397816578118 |
| | 35 | 9.57824112492371 |
| | 35.5 | 10.1085701985209 |
| | 36 | 10.6187949438421 |
| | 36.5 | 11.0984383109489 |
| | 37 | 11.5388820273597 |
| | 37.5 | 11.9330804752471 |
| | 38 | 12.2748509629365 |
| | 38.5 | 12.5668385019039 |
| | 39 | 12.8170988939902 |
| | 39.5 | 13.0334732602524 |
| | 40 | 13.2226311255033 |
| | 40.5 | 13.3909619494015 |
| | 41 | 13.5460299243656 |
| | 41.5 | 13.6975012721076 |
| | 42 | 13.8552199326252 |
| | 42.5 | 14.0280734917196 |
| | 43 | 14.2239791072318 |
| | 43.5 | 14.4486499299701 |
| | 44 | 14.6976029262957 |
| | 44.5 | 14.9633503043963 |
| | 45 | 15.2390099874977 |
| | 45.5 | 15.5201244628862 |
| | 46 | 15.8028440446553 |
| | 46.5 | 16.0836764347355 |
| | 47 | 16.3599267145246 |
| | 47.5 | 16.6289904297057 |
| | 48 | 16.8862747798961 |
| | 48.5 | 17.1233883431067 |
| | 49 | 17.3315158393844 |
| | 49.5 | 17.5019224255177 |
| | 50 | 17.6259487731912 |

| | | |
|---|---|---|
| | 50.5 | 17.6949970152694 |
| | 51 | 17.7043439395506 |
| | 51.5 | 17.6554902034071 |
| | 52 | 17.550509781061 |
| | 52.5 | 17.394624361604 |
| | 53 | 17.2004518701611 |
| | 53.5 | 16.9816654115395 |
| | 54 | 16.7508600625081 |
| | 54.5 | 16.5183244084937 |
| | 55 | 16.2939542503648 |
| | 55.5 | 16.0855106560958 |
| | 56 | 15.8986532104201 |
| | 56.5 | 15.7394829387327 |
| | 57 | 15.6158736914932 |
| | 57.5 | 15.5360409036348 |
| | 58 | 15.5043867450666 |
| | 58.5 | 15.5177008289654 |
| | 59 | 15.5705103576343 |
| | 59.5 | 15.6473043506473 |
| | 60 | 15.7280747782631 |
| | 60.5 | 15.7955709870313 |
| | 61 | 15.8407985514855 |
| | 61.5 | 15.856289495614 |
| | 62 | 15.834607373195 |
| | 62.5 | 15.7750000533934 |
| | 63 | 15.6916289958364 |
| | 63.5 | 15.6006735666832 |
| | 64 | 15.5180473010416 |
| | 64.5 | 15.4583511547636 |
| | 65 | 15.4357816740315 |
| | 65.5 | 15.4642687122618 |
| | 66 | 15.5524797269871 |
| | 66.5 | 15.7042615407421 |
| | 67 | 15.9230027418817 |
| | 67.5 | 16.2078989982907 |
| | 68 | 16.5549182593731 |
| | 68.5 | 16.9597532805051 |
| | 69 | 17.4132325065171 |
| | 69.5 | 17.9010595134162 |
| | 70 | 18.4086935551718 |
| | 70.5 | 18.9215415027777 |
| | 71 | 19.424647012201 |
| | 71.5 | 19.9028998586709 |

| | | |
|---|---|---|
| | 72 | 20.3424104832812 |
| | 72.5 | 20.7383149057167 |
| | 73 | 21.0897944367009 |
| | 73.5 | 21.3960494685186 |
| | 74 | 21.6579526089386 |
| | 74.5 | 21.8843874865007 |
| | 75 | 22.0866338637933 |
| | 75.5 | 22.2759719382775 |
| | 76 | 22.4636604210983 |
| | 76.5 | 22.6580805480676 |
| | 77 | 22.8618691485416 |
| | 77.5 | 23.07698076 |
| | 78 | 23.3053699273628 |
| | 78.5 | 23.5489911843894 |
| | 79 | 23.8097990722798 |
| | 79.5 | 24.0897481303736 |
| | 80 | 24.3892840603781 |
| | 80.5 | 24.7028172134703 |
| | 81 | 25.0232491031945 |
| | 81.5 | 25.3434812430955 |
| | 82 | 25.6565769781859 |
| | 82.5 | 25.9567217665496 |
| | 83 | 26.2385764639661 |
| | 83.5 | 26.4968051287756 |
| | 84 | 26.7272779446936 |
| | 84.5 | 26.9292459900927 |
| | 85 | 27.1025482770112 |
| | 85.5 | 27.2470238174875 |
| | 86 | 27.36251162 |
| | 86.5 | 27.4491399326924 |
| | 87 | 27.5089199301232 |
| | 87.5 | 27.5446164861343 |
| | 88 | 27.5589964697582 |
| | 88.5 | 27.5548267500273 |
| | 89 | 27.5348405718171 |
| | 89.5 | 27.5010658838508 |
| | 90 | 27.4548611973246 |
| | 90.5 | 27.3975585771247 |
| | 91 | 27.330490088137 |
| | 91.5 | 27.2549877952478 |
| | 92 | 27.1719425006544 |
| | 92.5 | 27.0804799557992 |
| | 93 | 26.9792846494357 |

| | | |
|---|---|---|
| | 93.5 | 26.8670410703174 |
| | 94 | 26.7424337071979 |
| | 94.5 | 26.6041470488308 |
| | 95 | 26.4507541676139 |
| | 95.5 | 26.2803824705224 |
| | 96 | 26.0910479481758 |
| | 96.5 | 25.8807665911935 |
| | 97 | 25.6477972503415 |
| | 97.5 | 25.3925845848272 |
| | 98 | 25.116729990795 |
| | 98.5 | 24.8218453980935 |
| | 99 | 24.5095427365717 |
| | 99.5 | 24.1816895309902 |
| | 100 | 23.8426154231428 |
| | 100.5 | 23.4980260249617 |
| | 101 | 23.1536404071147 |
| | 101.5 | 22.8145842396795 |
| | 102 | 22.4845006871027 |
| | 102.5 | 22.1668051973073 |
| | 103 | 21.8649132182161 |
| | 103.5 | 21.5822401977518 |
| | 104 | 21.3222015838376 |
| | 104.5 | 21.0882066096289 |
| | 105 | 20.8823469776193 |
| | 105.5 | 20.7037748053822 |
| | 106 | 20.5512444653852 |
| | 106.5 | 20.423510330096 |
| | 107 | 20.3193267719824 |
| | 107.5 | 20.237448163512 |
| | 108 | 20.1765696680054 |
| | 108.5 | 20.1345005045045 |
| | 109 | 20.1083678903911 |
| | 109.5 | 20.0952814995964 |
| | 110 | 20.0923510060513 |
| | 110.5 | 20.0966860836867 |
| | 111 | 20.1054487107786 |
| | 111.5 | 20.1161874274033 |
| | 112 | 20.1266240317797 |
| | 112.5 | 20.1344811393825 |
| | 113 | 20.137481365686 |
| | 113.5 | 20.1333473261648 |
| | 114 | 20.1198016362934 |
| | 114.5 | 20.0945669115462 |

| | | | | |
|---|---|---|---|---|
| | 115 | 20.0553657673979 | | |
| | 115.5 | 20.0004453618623 | | |
| | 116 | 19.9301510231116 | | |
| | 116.5 | 19.8453526218573 | | |
| | 117 | 19.7469200288108 | | |
| | 117.5 | 19.6357231146838 | | |
| | 118 | 19.5126317501878 | | |
| | 118.5 | 19.3785158060343 | | |
| | 119 | 19.2342451529349 | | |
| | 119.5 | 19.080689661601 | | |
| | 120 | 18.9187192027444 | | |
| | 120.5 | 18.7492036470764 | | |
| | 121 | 18.5730128653087 | | |
| | 121.5 | 18.3911189071187 | | |
| | 122 | 18.2056589532954 | | |
| | 122.5 | 18.0195112421022 | | |
| | 123 | 17.8355658934106 | | |
| | 123.5 | 17.656713027092 | | |
| | 124 | 17.4858426821414 | | |
| | 124.5 | 17.3256594142951 | | |
| | 125 | 17.1782867225006 | | |
| | 125.5 | 17.0457346287223 | | |
| | 126 | 16.9300131549245 | | |
| | 126.5 | 16.8331323230718 | | |
| | 127 | 16.7571021551285 | | |
| | 127.5 | 16.7039326085554 | | |
| | 128 | 16.6753097214662 | | |
| | 128.5 | 16.6718429780052 | | |
| | 129 | 16.6939184960176 | | |
| | 129.5 | 16.7419223933488 | | |
| | 130 | 16.816062312464 | | |
| | 130.5 | 16.9152268512212 | | |
| | 131 | 17.0377134077817 | | |
| | 131.5 | 17.1818165916287 | | |
| | 132 | 17.3458310122457 | | |
| | 132.5 | 17.5279207578679 | | |
| | 133 | 17.7257278317387 | | |
| Selenium | 0 | 0.61 | 1 Ma | Large et al. (2015) |
| | 1 | 0.607169811320755 | | |
| | 2 | 0.604339622641509 | | |
| | 3 | 0.601509433962264 | | |
| | 4 | 0.598679245283019 | | |
| | 5 | 0.595849056603774 | | |

| | | |
|---|---|---|
| | 6 | 0.593018867924528 |
| | 7 | 0.590188679245283 |
| | 8 | 0.587358490566038 |
| | 9 | 0.584528301886792 |
| | 10 | 0.581698113207547 |
| | 11 | 0.578867924528302 |
| | 12 | 0.576037735849057 |
| | 13 | 0.573207547169811 |
| | 14 | 0.570377358490566 |
| | 15 | 0.567547169811321 |
| | 16 | 0.564716981132075 |
| | 17 | 0.56188679245283 |
| | 18 | 0.559056603773585 |
| | 19 | 0.55622641509434 |
| | 20 | 0.553396226415094 |
| | 21 | 0.550566037735849 |
| | 22 | 0.547735849056604 |
| | 23 | 0.544905660377359 |
| | 24 | 0.542075471698113 |
| | 25 | 0.539245283018868 |
| | 26 | 0.536415094339623 |
| | 27 | 0.533584905660377 |
| | 28 | 0.530754716981132 |
| | 29 | 0.527924528301887 |
| | 30 | 0.525094339622642 |
| | 31 | 0.522264150943396 |
| | 32 | 0.519433962264151 |
| | 33 | 0.516603773584906 |
| | 34 | 0.51377358490566 |
| | 35 | 0.510943396226415 |
| | 36 | 0.50811320754717 |
| | 37 | 0.505283018867924 |
| | 38 | 0.502452830188679 |
| | 39 | 0.499622641509434 |
| | 40 | 0.496792452830189 |
| | 41 | 0.493962264150943 |
| | 42 | 0.491132075471698 |
| | 43 | 0.488301886792453 |
| | 44 | 0.485471698113208 |
| | 45 | 0.482641509433962 |
| | 46 | 0.479811320754717 |
| | 47 | 0.476981132075472 |
| | 48 | 0.474150943396226 |

| | | |
|---|---|---|
| | 49 | 0.471320754716981 |
| | 50 | 0.468490566037736 |
| | 51 | 0.465660377358491 |
| | 52 | 0.462830188679245 |
| | 53 | 0.46 |
| | 54 | 0.473333333333333 |
| | 55 | 0.486666666666667 |
| | 56 | 0.5 |
| | 57 | 0.486666666666667 |
| | 58 | 0.473333333333333 |
| | 59 | 0.46 |
| | 60 | 0.446666666666667 |
| | 61 | 0.433333333333333 |
| | 62 | 0.42 |
| | 63 | 0.406666666666667 |
| | 64 | 0.393333333333333 |
| | 65 | 0.38 |
| | 66 | 0.366666666666667 |
| | 67 | 0.353333333333333 |
| | 68 | 0.34 |
| | 69 | 0.326666666666667 |
| | 70 | 0.313333333333333 |
| | 71 | 0.3 |
| | 72 | 0.286666666666667 |
| | 73 | 0.273333333333333 |
| | 74 | 0.26 |
| | 75 | 0.246666666666667 |
| | 76 | 0.233333333333333 |
| | 77 | 0.22 |
| | 78 | 0.206666666666667 |
| | 79 | 0.193333333333333 |
| | 80 | 0.18 |
| | 81 | 0.166666666666667 |
| | 82 | 0.153333333333333 |
| | 83 | 0.14 |
| | 84 | 0.155 |
| | 85 | 0.17 |
| | 86 | 0.185 |
| | 87 | 0.2 |
| | 88 | 0.215 |
| | 89 | 0.23 |
| | 90 | 0.245 |
| | 91 | 0.26 |

| | | |
|---|---|---|
| | 92 | 0.11 |
| | 93 | 0.118181818181818 |
| | 94 | 0.126363636363636 |
| | 95 | 0.134545454545455 |
| | 96 | 0.142727272727273 |
| | 97 | 0.150909090909091 |
| | 98 | 0.159090909090909 |
| | 99 | 0.167272727272727 |
| | 100 | 0.175454545454545 |
| | 101 | 0.183636363636364 |
| | 102 | 0.191818181818182 |
| | 103 | 0.2 |
| | 104 | 0.208181818181818 |
| | 105 | 0.216363636363636 |
| | 106 | 0.224545454545455 |
| | 107 | 0.232727272727273 |
| | 108 | 0.240909090909091 |
| | 109 | 0.249090909090909 |
| | 110 | 0.257272727272727 |
| | 111 | 0.265454545454545 |
| | 112 | 0.273636363636364 |
| | 113 | 0.281818181818182 |
| | 114 | 0.29 |
| | 115 | 0.298181818181818 |
| | 116 | 0.306363636363636 |
| | 117 | 0.314545454545455 |
| | 118 | 0.322727272727273 |
| | 119 | 0.330909090909091 |
| | 120 | 0.339090909090909 |
| | 121 | 0.347272727272727 |
| | 122 | 0.355454545454546 |
| | 123 | 0.363636363636364 |
| | 124 | 0.371818181818182 |
| | 125 | 0.38 |
| | 126 | 0.333333333333333 |
| | 127 | 0.286666666666667 |
| | 128 | 0.24 |
| | 129 | 0.193333333333333 |
| | 130 | 0.146666666666667 |
| | 131 | 0.1 |
| | 132 | 0.107272727272727 |
| | 133 | 0.114545454545455 |

**Supplementary Table 6**. Time series of biotic and abiotic factors (100-300 Ma) used in the multivariate birth-death (MBD) model analyses. This table presents the temporal series of 11 predictor variables used in the MBD model to evaluate the drivers of diversification rates of four major land plant groups (Bryophytes, Pteridophytes, Gymnosperms, and Angiosperms). These data were used in the MBD analyses for Figure 5A-C.

| Factors | Age | Value | resolution | references |
|---|---|---|---|---|
| Arid | 100 | 0.22 | 1 Ma | Lehtonen et al. (2017) |
| | 101 | 0.239 | | |
| | 102 | 0.258 | | |
| | 103 | 0.277 | | |
| | 104 | 0.296 | | |
| | 105 | 0.315 | | |
| | 106 | 0.334 | | |
| | 107 | 0.353 | | |
| | 108 | 0.372 | | |
| | 109 | 0.391 | | |
| | 110 | 0.41 | | |
| | 111 | 0.429 | | |
| | 112 | 0.448 | | |
| | 113 | 0.467 | | |
| | 114 | 0.486 | | |
| | 115 | 0.505 | | |
| | 116 | 0.524 | | |
| | 117 | 0.543 | | |
| | 118 | 0.562 | | |
| | 119 | 0.581 | | |
| | 120 | 0.6 | | |
| | 121 | 0.6065 | | |
| | 122 | 0.613 | | |
| | 123 | 0.6195 | | |
| | 124 | 0.626 | | |
| | 125 | 0.6325 | | |
| | 126 | 0.639 | | |
| | 127 | 0.6455 | | |
| | 128 | 0.652 | | |
| | 129 | 0.6585 | | |
| | 130 | 0.665 | | |
| | 131 | 0.6715 | | |
| | 132 | 0.678 | | |
| | 133 | 0.6845 | | |
| | 134 | 0.691 | | |
| | 135 | 0.6975 | | |
| | 136 | 0.704 | | |

| | | |
|---|---|---|
| | 137 | 0.7105 |
| | 138 | 0.717 |
| | 139 | 0.7235 |
| | 140 | 0.73 |
| | 141 | 0.7365 |
| | 142 | 0.743 |
| | 143 | 0.7495 |
| | 144 | 0.756 |
| | 145 | 0.7625 |
| | 146 | 0.769 |
| | 147 | 0.7755 |
| | 148 | 0.782 |
| | 149 | 0.7885 |
| | 150 | 0.795 |
| | 151 | 0.8015 |
| | 152 | 0.808 |
| | 153 | 0.8145 |
| | 154 | 0.821 |
| | 155 | 0.8275 |
| | 156 | 0.834 |
| | 157 | 0.8405 |
| | 158 | 0.847 |
| | 159 | 0.8535 |
| | 160 | 0.86 |
| | 161 | 0.8485 |
| | 162 | 0.837 |
| | 163 | 0.8255 |
| | 164 | 0.814 |
| | 165 | 0.8025 |
| | 166 | 0.791 |
| | 167 | 0.7795 |
| | 168 | 0.768 |
| | 169 | 0.7565 |
| | 170 | 0.745 |
| | 171 | 0.7335 |
| | 172 | 0.722 |
| | 173 | 0.7105 |
| | 174 | 0.699 |
| | 175 | 0.6875 |
| | 176 | 0.676 |
| | 177 | 0.6645 |
| | 178 | 0.653 |
| | 179 | 0.6415 |

| | | |
|---|---|---|
| | 180 | 0.63 |
| | 181 | 0.630666666666667 |
| | 182 | 0.631333333333333 |
| | 183 | 0.632 |
| | 184 | 0.632666666666667 |
| | 185 | 0.633333333333333 |
| | 186 | 0.634 |
| | 187 | 0.634666666666667 |
| | 188 | 0.635333333333333 |
| | 189 | 0.636 |
| | 190 | 0.636666666666667 |
| | 191 | 0.637333333333333 |
| | 192 | 0.638 |
| | 193 | 0.638666666666667 |
| | 194 | 0.639333333333333 |
| | 195 | 0.64 |
| | 196 | 0.640666666666667 |
| | 197 | 0.641333333333333 |
| | 198 | 0.642 |
| | 199 | 0.642666666666667 |
| | 200 | 0.643333333333333 |
| | 201 | 0.644 |
| | 202 | 0.644666666666667 |
| | 203 | 0.645333333333333 |
| | 204 | 0.646 |
| | 205 | 0.646666666666667 |
| | 206 | 0.647333333333333 |
| | 207 | 0.648 |
| | 208 | 0.648666666666667 |
| | 209 | 0.649333333333333 |
| | 210 | 0.65 |
| | 211 | 0.666 |
| | 212 | 0.682 |
| | 213 | 0.698 |
| | 214 | 0.714 |
| | 215 | 0.73 |
| | 216 | 0.746 |
| | 217 | 0.762 |
| | 218 | 0.778 |
| | 219 | 0.794 |
| | 220 | 0.81 |
| | 221 | 0.8195 |
| | 222 | 0.829 |

| | | |
|---|---|---|
| | 223 | 0.8385 |
| | 224 | 0.848 |
| | 225 | 0.8575 |
| | 226 | 0.867 |
| | 227 | 0.8765 |
| | 228 | 0.886 |
| | 229 | 0.8955 |
| | 230 | 0.905 |
| | 231 | 0.9145 |
| | 232 | 0.924 |
| | 233 | 0.9335 |
| | 234 | 0.943 |
| | 235 | 0.9525 |
| | 236 | 0.962 |
| | 237 | 0.9715 |
| | 238 | 0.981 |
| | 239 | 0.9905 |
| | 240 | 1 |
| | 241 | 0.9795 |
| | 242 | 0.959 |
| | 243 | 0.9385 |
| | 244 | 0.918 |
| | 245 | 0.8975 |
| | 246 | 0.877 |
| | 247 | 0.8565 |
| | 248 | 0.836 |
| | 249 | 0.8155 |
| | 250 | 0.795 |
| | 251 | 0.7745 |
| | 252 | 0.754 |
| | 253 | 0.7335 |
| | 254 | 0.713 |
| | 255 | 0.6925 |
| | 256 | 0.672 |
| | 257 | 0.6515 |
| | 258 | 0.631 |
| | 259 | 0.6105 |
| | 260 | 0.59 |
| | 261 | 0.5605 |
| | 262 | 0.531 |
| | 263 | 0.5015 |
| | 264 | 0.472 |
| | 265 | 0.4425 |

| | | | | |
|---|---|---|---|---|
| | 266 | 0.413 | | |
| | 267 | 0.3835 | | |
| | 268 | 0.354 | | |
| | 269 | 0.3245 | | |
| | 270 | 0.295 | | |
| | 271 | 0.2655 | | |
| | 272 | 0.236 | | |
| | 273 | 0.2065 | | |
| | 274 | 0.177 | | |
| | 275 | 0.1475 | | |
| | 276 | 0.118 | | |
| | 277 | 0.0885 | | |
| | 278 | 0.0590000000000001 | | |
| | 279 | 0.0295 | | |
| | 280 | 0 | | |
| | 281 | 0.005 | | |
| | 282 | 0.01 | | |
| | 283 | 0.015 | | |
| | 284 | 0.02 | | |
| | 285 | 0.025 | | |
| | 286 | 0.03 | | |
| | 287 | 0.035 | | |
| | 288 | 0.04 | | |
| | 289 | 0.045 | | |
| | 290 | 0.05 | | |
| | 291 | 0.055 | | |
| | 292 | 0.06 | | |
| | 293 | 0.065 | | |
| | 294 | 0.07 | | |
| | 295 | 0.075 | | |
| | 296 | 0.08 | | |
| | 297 | 0.085 | | |
| | 298 | 0.09 | | |
| | 299 | 0.095 | | |
| | 300 | 0.1 | | |
| Atmospheric Carbon | 100 | 0.278888888888889 | 1 Ma | Berner (2001) |
| | 101 | 0.284444444444444 | | |
| | 102 | 0.29 | | |
| | 103 | 0.294545454545455 | | |
| | 104 | 0.299090909090909 | | |
| | 105 | 0.303636363636364 | | |
| | 106 | 0.308181818181818 | | |
| | 107 | 0.312727272727273 | | |

| | | |
|---|---|---|
| | 108 | 0.317272727272727 |
| | 109 | 0.321818181818182 |
| | 110 | 0.326363636363636 |
| | 111 | 0.330909090909091 |
| | 112 | 0.335454545454545 |
| | 113 | 0.34 |
| | 114 | 0.34125 |
| | 115 | 0.3425 |
| | 116 | 0.34375 |
| | 117 | 0.345 |
| | 118 | 0.34625 |
| | 119 | 0.3475 |
| | 120 | 0.34875 |
| | 121 | 0.35 |
| | 122 | 0.353636363636364 |
| | 123 | 0.357272727272727 |
| | 124 | 0.360909090909091 |
| | 125 | 0.364545454545455 |
| | 126 | 0.368181818181818 |
| | 127 | 0.371818181818182 |
| | 128 | 0.375454545454545 |
| | 129 | 0.379090909090909 |
| | 130 | 0.382727272727273 |
| | 131 | 0.386363636363636 |
| | 132 | 0.39 |
| | 133 | 0.399166666666667 |
| | 134 | 0.408333333333333 |
| | 135 | 0.4175 |
| | 136 | 0.426666666666667 |
| | 137 | 0.435833333333333 |
| | 138 | 0.445 |
| | 139 | 0.454166666666667 |
| | 140 | 0.463333333333333 |
| | 141 | 0.4725 |
| | 142 | 0.481666666666667 |
| | 143 | 0.490833333333333 |
| | 144 | 0.5 |
| | 145 | 0.495 |
| | 146 | 0.49 |
| | 147 | 0.485 |
| | 148 | 0.48 |
| | 149 | 0.475 |
| | 150 | 0.47 |

| | | |
|---|---|---|
| | 151 | 0.465 |
| | 152 | 0.46 |
| | 153 | 0.47 |
| | 154 | 0.48 |
| | 155 | 0.49 |
| | 156 | 0.5 |
| | 157 | 0.51 |
| | 158 | 0.52 |
| | 159 | 0.53 |
| | 160 | 0.54 |
| | 161 | 0.55 |
| | 162 | 0.56 |
| | 163 | 0.57 |
| | 164 | 0.565555555555555 |
| | 165 | 0.561111111111111 |
| | 166 | 0.556666666666667 |
| | 167 | 0.552222222222222 |
| | 168 | 0.547777777777778 |
| | 169 | 0.543333333333333 |
| | 170 | 0.538888888888889 |
| | 171 | 0.534444444444444 |
| | 172 | 0.53 |
| | 173 | 0.506363636363636 |
| | 174 | 0.482727272727273 |
| | 175 | 0.459090909090909 |
| | 176 | 0.435454545454545 |
| | 177 | 0.411818181818182 |
| | 178 | 0.388181818181818 |
| | 179 | 0.364545454545455 |
| | 180 | 0.340909090909091 |
| | 181 | 0.317272727272727 |
| | 182 | 0.293636363636364 |
| | 183 | 0.27 |
| | 184 | 0.266666666666667 |
| | 185 | 0.263333333333333 |
| | 186 | 0.26 |
| | 187 | 0.256666666666667 |
| | 188 | 0.253333333333333 |
| | 189 | 0.25 |
| | 190 | 0.246666666666667 |
| | 191 | 0.243333333333333 |
| | 192 | 0.24 |
| | 193 | 0.247 |

| | | |
|---|---|---|
| | 194 | 0.254 |
| | 195 | 0.261 |
| | 196 | 0.268 |
| | 197 | 0.275 |
| | 198 | 0.282 |
| | 199 | 0.289 |
| | 200 | 0.296 |
| | 201 | 0.303 |
| | 202 | 0.31 |
| | 203 | 0.307 |
| | 204 | 0.304 |
| | 205 | 0.301 |
| | 206 | 0.298 |
| | 207 | 0.295 |
| | 208 | 0.292 |
| | 209 | 0.289 |
| | 210 | 0.286 |
| | 211 | 0.283 |
| | 212 | 0.28 |
| | 213 | 0.284545454545455 |
| | 214 | 0.289090909090909 |
| | 215 | 0.293636363636364 |
| | 216 | 0.298181818181818 |
| | 217 | 0.302727272727273 |
| | 218 | 0.307272727272727 |
| | 219 | 0.311818181818182 |
| | 220 | 0.316363636363636 |
| | 221 | 0.320909090909091 |
| | 222 | 0.325454545454545 |
| | 223 | 0.33 |
| | 224 | 0.325 |
| | 225 | 0.32 |
| | 226 | 0.315 |
| | 227 | 0.31 |
| | 228 | 0.305 |
| | 229 | 0.3 |
| | 230 | 0.295 |
| | 231 | 0.29 |
| | 232 | 0.301818181818182 |
| | 233 | 0.313636363636364 |
| | 234 | 0.325454545454545 |
| | 235 | 0.337272727272727 |
| | 236 | 0.349090909090909 |

| | | |
|---|---|---|
| | 237 | 0.360909090909091 |
| | 238 | 0.372727272727273 |
| | 239 | 0.384545454545455 |
| | 240 | 0.396363636363636 |
| | 241 | 0.408181818181818 |
| | 242 | 0.42 |
| | 243 | 0.412222222222222 |
| | 244 | 0.404444444444444 |
| | 245 | 0.396666666666667 |
| | 246 | 0.388888888888889 |
| | 247 | 0.381111111111111 |
| | 248 | 0.373333333333333 |
| | 249 | 0.365555555555556 |
| | 250 | 0.357777777777778 |
| | 251 | 0.35 |
| | 252 | 0.323636363636364 |
| | 253 | 0.297272727272727 |
| | 254 | 0.270909090909091 |
| | 255 | 0.244545454545455 |
| | 256 | 0.218181818181818 |
| | 257 | 0.191818181818182 |
| | 258 | 0.165454545454545 |
| | 259 | 0.139090909090909 |
| | 260 | 0.112727272727273 |
| | 261 | 0.0863636363636364 |
| | 262 | 0.06 |
| | 263 | 0.0563636363636364 |
| | 264 | 0.0527272727272727 |
| | 265 | 0.0490909090909091 |
| | 266 | 0.0454545454545455 |
| | 267 | 0.0418181818181818 |
| | 268 | 0.0381818181818182 |
| | 269 | 0.0345454545454545 |
| | 270 | 0.0309090909090909 |
| | 271 | 0.0272727272727273 |
| | 272 | 0.0236363636363636 |
| | 273 | 0.02 |
| | 274 | 0.01875 |
| | 275 | 0.0175 |
| | 276 | 0.01625 |
| | 277 | 0.015 |
| | 278 | 0.01375 |
| | 279 | 0.0125 |

| | | | | |
|---|---|---|---|---|
| | 280 | 0.01125 | | |
| | 281 | 0.01 | | |
| | 282 | 0.011 | | |
| | 283 | 0.012 | | |
| | 284 | 0.013 | | |
| | 285 | 0.014 | | |
| | 286 | 0.015 | | |
| | 287 | 0.016 | | |
| | 288 | 0.017 | | |
| | 289 | 0.018 | | |
| | 290 | 0.019 | | |
| | 291 | 0.02 | | |
| | 292 | 0.0192307692307692 | | |
| | 293 | 0.0184615384615385 | | |
| | 294 | 0.0176923076923077 | | |
| | 295 | 0.0169230769230769 | | |
| | 296 | 0.0161538461538462 | | |
| | 297 | 0.0153846153846154 | | |
| | 298 | 0.0146153846153846 | | |
| | 299 | 0.0138461538461538 | | |
| | 300 | 0.0130769230769231 | | |
| Bryophyta | 100.052749635994 | 0.220415608043849 | 1 Ma | This study |
| | 101.053277132354 | 0.221911080277685 | | |
| | 102.053804628713 | 0.226057512508212 | | |
| | 103.054332125073 | 0.227272727272727 | | |
| | 104.054859621433 | 0.228253628415351 | | |
| | 105.055387117793 | 0.227759732055482 | | |
| | 106.055914614153 | 0.228501725191471 | | |
| | 107.056442110513 | 0.230704732675724 | | |
| | 108.056969606873 | 0.232163449831616 | | |
| | 109.057497103233 | 0.231200926227964 | | |
| | 110.058024599593 | 0.232655048999113 | | |
| | 111.058552095953 | 0.235078586951028 | | |
| | 112.059079592313 | 0.236100837556339 | | |
| | 113.059607088673 | 0.249564682045604 | | |
| | 114.060134585033 | 0.256849079055578 | | |
| | 115.060662081393 | 0.257637016038997 | | |
| | 116.061189577753 | 0.252034163844949 | | |
| | 117.061717074113 | 0.247414509985895 | | |
| | 118.062244570472 | 0.240842242611081 | | |
| | 119.062772066832 | 0.236663649687352 | | |
| | 120.063299563192 | 0.231777521513207 | | |
| | 121.063827059552 | 0.227169353616011 | | |

| | | |
|---|---|---|
| | 122.064354555912 | 0.216974413871366 |
| | 123.064882052272 | 0.214314265105188 |
| | 124.065409548632 | 0.212357057204684 |
| | 125.065937044992 | 0.207634029688914 |
| | 126.066464541352 | 0.202171306229526 |
| | 127.066992037712 | 0.199240088763513 |
| | 128.067519534072 | 0.196993434624202 |
| | 129.068047030432 | 0.193763582149896 |
| | 130.068574526792 | 0.193083613207937 |
| | 131.069102023152 | 0.19737706575024 |
| | 132.069629519512 | 0.200310580408625 |
| | 133.070157015872 | 0.200308283216253 |
| | 134.070684512231 | 0.205561962169836 |
| | 135.071212008591 | 0.207512278493226 |
| | 136.071739504951 | 0.214001846942667 |
| | 137.072267001311 | 0.212986487914471 |
| | 138.072794497671 | 0.209421245353928 |
| | 139.073321994031 | 0.208761951143313 |
| | 140.073849490391 | 0.206471650348944 |
| | 141.074376986751 | 0.201916317876292 |
| | 142.074904483111 | 0.196373192683902 |
| | 143.075431979471 | 0.187905741602614 |
| | 144.075959475831 | 0.179759897453333 |
| | 145.076486972191 | 0.171609458919309 |
| | 146.077014468551 | 0.156016117101678 |
| | 147.077541964911 | 0.148444571045269 |
| | 148.078069461271 | 0.144534749629003 |
| | 149.078596957631 | 0.140282646549387 |
| | 150.07912445399 | 0.137983156985532 |
| | 151.07965195035 | 0.13539422118287 |
| | 152.08017944671 | 0.133439310474738 |
| | 153.08070694307 | 0.131484399766605 |
| | 154.08123443943 | 0.130508093008725 |
| | 155.08176193579 | 0.130154325383516 |
| | 156.08228943215 | 0.130154325383516 |
| | 157.08281692851 | 0.129536380635587 |
| | 158.08334442487 | 0.128197117483012 |
| | 159.08387192123 | 0.12697041675664 |
| | 160.08439941759 | 0.122817092948998 |
| | 161.08492691395 | 0.120738133852805 |
| | 162.08545441031 | 0.120368285880996 |
| | 163.08598190667 | 0.120368285880996 |
| | 164.08650940303 | 0.120368285880996 |

| | | |
|---|---|---|
| | 165.087036899389 | 0.120368285880996 |
| | 166.087564395749 | 0.120368285880996 |
| | 167.088091892109 | 0.120368285880996 |
| | 168.088619388469 | 0.120368285880996 |
| | 169.089146884829 | 0.120368285880996 |
| | 170.089674381189 | 0.118213519436545 |
| | 171.090201877549 | 0.116846689975512 |
| | 172.090729373909 | 0.115870383217631 |
| | 173.091256870269 | 0.113915472509499 |
| | 174.091784366629 | 0.11253945427898 |
| | 175.092311862989 | 0.11253945427898 |
| | 176.092839359349 | 0.111965156186109 |
| | 177.093366855709 | 0.111560850328728 |
| | 178.093894352069 | 0.111560850328728 |
| | 179.094421848429 | 0.112695663360241 |
| | 180.094949344789 | 0.113518058229232 |
| | 181.095476841148 | 0.113518058229232 |
| | 182.096004337508 | 0.113099969217622 |
| | 183.096531833868 | 0.114634493721773 |
| | 184.097059330228 | 0.116453870079988 |
| | 185.097586826588 | 0.117007493441516 |
| | 186.098114322948 | 0.118535126368552 |
| | 187.098641819308 | 0.120368285880996 |
| | 188.099169315668 | 0.120368285880996 |
| | 189.099696812028 | 0.120368285880996 |
| | 190.100224308388 | 0.120368285880996 |
| | 191.100751804748 | 0.120368285880996 |
| | 192.101279301108 | 0.120368285880996 |
| | 193.101806797468 | 0.120368285880996 |
| | 194.102334293828 | 0.120368285880996 |
| | 195.102861790188 | 0.120368285880996 |
| | 196.103389286547 | 0.120368285880996 |
| | 197.103916782907 | 0.120368285880996 |
| | 198.104444279267 | 0.120368285880996 |
| | 199.104971775627 | 0.120368285880996 |
| | 200.105499271987 | 0.120368285880996 |
| | 201.106026768347 | 0.120368285880996 |
| | 202.106554264707 | 0.120368285880996 |
| | 203.107081761067 | 0.120368285880996 |
| | 204.107609257427 | 0.120368285880996 |
| | 205.108136753787 | 0.120368285880996 |
| | 206.108664250147 | 0.120368285880996 |
| | 207.109191746507 | 0.120368285880996 |

| | | |
|---|---|---|
| | 208.109719242867 | 0.120368285880996 |
| | 209.110246739227 | 0.120368285880996 |
| | 210.110774235587 | 0.120368285880996 |
| | 211.111301731947 | 0.120368285880996 |
| | 212.111829228306 | 0.120368285880996 |
| | 213.112356724666 | 0.11987898390587 |
| | 214.112884221026 | 0.119389681930744 |
| | 215.113411717386 | 0.119389681930744 |
| | 216.113939213746 | 0.119389681930744 |
| | 217.114466710106 | 0.118909568725104 |
| | 218.114994206466 | 0.11743247403024 |
| | 219.115521702826 | 0.116956955209343 |
| | 220.116049199186 | 0.115948487758262 |
| | 221.116576695546 | 0.116924794516142 |
| | 222.117104191906 | 0.11794245073671 |
| | 223.117631688266 | 0.115987540028577 |
| | 224.118159184626 | 0.116403331847816 |
| | 225.118686680986 | 0.11743247403024 |
| | 226.119214177346 | 0.119389681930744 |
| | 227.119741673706 | 0.119846823212669 |
| | 228.120269170065 | 0.119844526020298 |
| | 229.120796666425 | 0.120368285880996 |
| | 230.121324162785 | 0.121346889831248 |
| | 231.121851659145 | 0.121346889831248 |
| | 232.122379155505 | 0.121346889831248 |
| | 233.122906651865 | 0.121346889831248 |
| | 234.123434148225 | 0.122766554716825 |
| | 235.123961644585 | 0.123742861474706 |
| | 236.124489140945 | 0.124282701682004 |
| | 237.125016637305 | 0.124282701682004 |
| | 238.125544133665 | 0.125693177798095 |
| | 239.126071630025 | 0.12721851353276 |
| | 240.126599126385 | 0.128624395264108 |
| | 241.127126622745 | 0.131132929333768 |
| | 242.127654119105 | 0.132956900076726 |
| | 243.128181615464 | 0.134489127388506 |
| | 244.128709111824 | 0.135047345134776 |
| | 245.129236608184 | 0.136025949085028 |
| | 246.129764104544 | 0.136025949085028 |
| | 247.130291600904 | 0.135614751650533 |
| | 248.130819097264 | 0.131702633041896 |
| | 249.131346593624 | 0.128769118383512 |
| | 250.131874089984 | 0.126409901817998 |

| | | |
|---|---|---|
| | 251.132401586344 | 0.132915550614039 |
| | 252.132929082704 | 0.133669029711886 |
| | 253.133456579064 | 0.132662859453176 |
| | 254.133984075424 | 0.125674800259123 |
| | 255.134511571784 | 0.124468774264095 |
| | 256.135039068144 | 0.124282701682004 |
| | 257.135566564504 | 0.128585342993793 |
| | 258.136094060864 | 0.122146312776524 |
| | 259.136621557223 | 0.11806649912477 |
| | 260.137149053583 | 0.117216537947321 |
| | 261.137676549943 | 0.1223254937815 |
| | 262.138204046303 | 0.122927358182829 |
| | 263.138731542663 | 0.12721851353276 |
| | 264.139259039023 | 0.128197117483012 |
| | 265.139786535383 | 0.129175721433264 |
| | 266.140314031743 | 0.129175721433264 |
| | 267.140841528103 | 0.12954097502033 |
| | 268.141369024463 | 0.130154325383516 |
| | 269.141896520823 | 0.130154325383516 |
| | 270.142424017183 | 0.130154325383516 |
| | 271.142951513543 | 0.130154325383516 |
| | 272.143479009903 | 0.130508093008725 |
| | 273.144006506263 | 0.131132929333768 |
| | 274.144534002623 | 0.133090137234272 |
| | 275.145061498982 | 0.133437013282366 |
| | 276.145588995342 | 0.134757898895969 |
| | 277.146116491702 | 0.137346834698631 |
| | 278.146643988062 | 0.136664568564301 |
| | 279.147171484422 | 0.12910680566212 |
| | 280.147698980782 | 0.122297927473042 |
| | 281.148226477142 | 0.110520222184446 |
| | 282.148753973502 | 0.0955448251147444 |
| | 283.149281469862 | 0.0795747437481912 |
| | 284.149808966222 | 0.0681760752008895 |
| | 285.150336462582 | 0.0587231285922345 |
| | 286.150863958942 | 0.0531800033998448 |
| | 287.151391455302 | 0.0512273898840837 |
| | 288.151918951662 | 0.0495917889155874 |
| | 289.152446448022 | 0.0476368782074549 |
| | 290.152973944382 | 0.0459943856618441 |
| | 291.153501440741 | 0.0437270567911899 |
| | 292.154028937101 | 0.0430585738110881 |
| | 293.154556433461 | 0.0430585738110881 |

| | | | | |
|---|---|---|---|---|
| | 294.155083929821 | 0.0420799698608361 | | |
| | 295.155611426181 | 0.0420799698608361 | | |
| | 296.156138922541 | 0.0420799698608361 | | |
| | 297.156666418901 | 0.0420799698608361 | | |
| | 298.157193915261 | 0.0420799698608361 | | |
| | 299.157721411621 | 0.0420799698608361 | | |
| | 300.158248907981 | 0.0420799698608361 | | |
| Fragmention | 100 | 0.507375269 | 1 Ma | Zaffos et al. (2017) |
| | 101 | 0.50722461 | | |
| | 102 | 0.507149371 | | |
| | 103 | 0.507064931 | | |
| | 104 | 0.507104446 | | |
| | 105 | 0.507036838 | | |
| | 106 | 0.505580138 | | |
| | 107 | 0.499793016 | | |
| | 108 | 0.498585868 | | |
| | 109 | 0.497576577 | | |
| | 110 | 0.496779453 | | |
| | 111 | 0.486903642 | | |
| | 112 | 0.485679678 | | |
| | 113 | 0.461759614 | | |
| | 114 | 0.46023643 | | |
| | 115 | 0.449990884 | | |
| | 116 | 0.452444471 | | |
| | 117 | 0.444765012 | | |
| | 118 | 0.437508142 | | |
| | 119 | 0.435686482 | | |
| | 120 | 0.436840605 | | |
| | 121 | 0.457002974 | | |
| | 122 | 0.451363839 | | |
| | 123 | 0.451851659 | | |
| | 124 | 0.451873317 | | |
| | 125 | 0.451854357 | | |
| | 126 | 0.453521917 | | |
| | 127 | 0.438506176 | | |
| | 128 | 0.439523824 | | |
| | 129 | 0.439967133 | | |
| | 130 | 0.440364274 | | |
| | 131 | 0.420199518 | | |
| | 132 | 0.409684229 | | |
| | 133 | 0.409752889 | | |
| | 134 | 0.40699834 | | |
| | 135 | 0.408219561 | | |

| | | |
|---|---|---|
| | 136 | 0.394997976 |
| | 137 | 0.397791301 |
| | 138 | 0.396868939 |
| | 139 | 0.39544581 |
| | 140 | 0.394603246 |
| | 141 | 0.397572394 |
| | 142 | 0.397169107 |
| | 143 | 0.396888667 |
| | 144 | 0.385025964 |
| | 145 | 0.384918544 |
| | 146 | 0.390567083 |
| | 147 | 0.382128879 |
| | 148 | 0.381228038 |
| | 149 | 0.380930104 |
| | 150 | 0.380724673 |
| | 151 | 0.380228646 |
| | 152 | 0.380089064 |
| | 153 | 0.37946688 |
| | 154 | 0.379102488 |
| | 155 | 0.368623258 |
| | 156 | 0.367677472 |
| | 157 | 0.367420113 |
| | 158 | 0.367221755 |
| | 159 | 0.368626886 |
| | 160 | 0.367991487 |
| | 161 | 0.365285496 |
| | 162 | 0.364831842 |
| | 163 | 0.364035221 |
| | 164 | 0.364087192 |
| | 165 | 0.364719487 |
| | 166 | 0.36458252 |
| | 167 | 0.364339083 |
| | 168 | 0.325370866 |
| | 169 | 0.324895513 |
| | 170 | 0.324699843 |
| | 171 | 0.325273753 |
| | 172 | 0.321419973 |
| | 173 | 0.31954972 |
| | 174 | 0.319688074 |
| | 175 | 0.312802378 |
| | 176 | 0.312986948 |
| | 177 | 0.313293931 |
| | 178 | 0.3148821 |

| | | |
|---|---|---|
| | 179 | 0.316153026 |
| | 180 | 0.317832826 |
| | 181 | 0.317610584 |
| | 182 | 0.31672822 |
| | 183 | 0.316622831 |
| | 184 | 0.316423553 |
| | 185 | 0.316666936 |
| | 186 | 0.316627636 |
| | 187 | 0.316572006 |
| | 188 | 0.316504003 |
| | 189 | 0.316381967 |
| | 190 | 0.316314636 |
| | 191 | 0.316052718 |
| | 192 | 0.315814697 |
| | 193 | 0.315563055 |
| | 194 | 0.316132171 |
| | 195 | 0.315873324 |
| | 196 | 0.333155366 |
| | 197 | 0.333172978 |
| | 198 | 0.332798915 |
| | 199 | 0.332553334 |
| | 200 | 0.332647266 |
| | 201 | 0.331010949 |
| | 202 | 0.330526514 |
| | 203 | 0.330667738 |
| | 204 | 0.330215853 |
| | 205 | 0.328400115 |
| | 206 | 0.326907179 |
| | 207 | 0.320920414 |
| | 208 | 0.319465117 |
| | 209 | 0.318649592 |
| | 210 | 0.318670523 |
| | 211 | 0.32220427 |
| | 212 | 0.322735632 |
| | 213 | 0.329885318 |
| | 214 | 0.330337669 |
| | 215 | 0.330910118 |
| | 216 | 0.329714832 |
| | 217 | 0.329648595 |
| | 218 | 0.328176714 |
| | 219 | 0.327752789 |
| | 220 | 0.327156978 |
| | 221 | 0.326505829 |

| | | |
|---|---|---|
| | 222 | 0.32583617 |
| | 223 | 0.325341847 |
| | 224 | 0.324917612 |
| | 225 | 0.324845916 |
| | 226 | 0.324620739 |
| | 227 | 0.324490696 |
| | 228 | 0.324444048 |
| | 229 | 0.32440493 |
| | 230 | 0.324178003 |
| | 231 | 0.323720558 |
| | 232 | 0.321545797 |
| | 233 | 0.318859922 |
| | 234 | 0.317563638 |
| | 235 | 0.314682779 |
| | 236 | 0.315508498 |
| | 237 | 0.314928703 |
| | 238 | 0.314492154 |
| | 239 | 0.314099987 |
| | 240 | 0.313726375 |
| | 241 | 0.313720032 |
| | 242 | 0.313678779 |
| | 243 | 0.313793623 |
| | 244 | 0.31554136 |
| | 245 | 0.315251991 |
| | 246 | 0.334964673 |
| | 247 | 0.335663526 |
| | 248 | 0.335186856 |
| | 249 | 0.334848637 |
| | 250 | 0.335937236 |
| | 251 | 0.327694794 |
| | 252 | 0.325474957 |
| | 253 | 0.326689806 |
| | 254 | 0.326566238 |
| | 255 | 0.32639975 |
| | 256 | 0.327414442 |
| | 257 | 0.332217446 |
| | 258 | 0.333766757 |
| | 259 | 0.331797986 |
| | 260 | 0.332310561 |
| | 261 | 0.343486929 |
| | 262 | 0.344711088 |
| | 263 | 0.345517368 |
| | 264 | 0.347115211 |

| | | | | |
|---|---|---|---|---|
| | 265 | 0.348157117 | | |
| | 266 | 0.347117285 | | |
| | 267 | 0.34885025 | | |
| | 268 | 0.348952491 | | |
| | 269 | 0.350053648 | | |
| | 270 | 0.35162668 | | |
| | 271 | 0.354752948 | | |
| | 272 | 0.35556515 | | |
| | 273 | 0.356549941 | | |
| | 274 | 0.357646049 | | |
| | 275 | 0.358844399 | | |
| | 276 | 0.360489848 | | |
| | 277 | 0.362543608 | | |
| | 278 | 0.363690446 | | |
| | 279 | 0.364366801 | | |
| | 280 | 0.365462836 | | |
| | 281 | 0.364637398 | | |
| | 282 | 0.364326601 | | |
| | 283 | 0.363872054 | | |
| | 284 | 0.36358511 | | |
| | 285 | 0.363405847 | | |
| | 286 | 0.363145796 | | |
| | 287 | 0.362579051 | | |
| | 288 | 0.36162074 | | |
| | 289 | 0.36063529 | | |
| | 290 | 0.359807719 | | |
| | 291 | 0.359027307 | | |
| | 292 | 0.357884108 | | |
| | 293 | 0.356036547 | | |
| | 294 | 0.354592529 | | |
| | 295 | 0.353517346 | | |
| | 296 | 0.35291411 | | |
| | 297 | 0.352249371 | | |
| | 298 | 0.351534489 | | |
| | 299 | 0.350919106 | | |
| | 300 | 0.35053135 | | |
| Gymnospermae | 100.255721561398 | 0.850340136054422 | 1 Ma | This study |
| | 101.258278777012 | 0.828571428571429 | | |
| | 102.260835992626 | 0.814965986394558 | | |
| | 103.26339320824 | 0.814965986394558 | | |
| | 104.265950423854 | 0.83265306122449 | | |
| | 105.268507639468 | 0.842176870748299 | | |
| | 106.271064855082 | 0.853061224489796 | | |

| | | |
|---|---|---|
| | 107.273622070696 | 0.86530612244898 |
| | 108.27617928631 | 0.87891156462585 |
| | 109.278736501924 | 0.887074829931973 |
| | 110.281293717538 | 0.902040816326531 |
| | 111.283850933151 | 0.922448979591837 |
| | 112.286408148765 | 0.938775510204082 |
| | 113.288965364379 | 0.94421768707483 |
| | 114.291522579993 | 0.948299319727891 |
| | 115.294079795607 | 0.952380952380952 |
| | 116.296637011221 | 0.952380952380952 |
| | 117.299194226835 | 0.948299319727891 |
| | 118.301751442449 | 0.949659863945578 |
| | 119.304308658063 | 0.949659863945578 |
| | 120.306865873677 | 0.951020408163265 |
| | 121.309423089291 | 0.956462585034014 |
| | 122.311980304905 | 0.953741496598639 |
| | 123.314537520519 | 0.951020408163265 |
| | 124.317094736133 | 0.949659863945578 |
| | 125.319651951747 | 0.949659863945578 |
| | 126.322209167361 | 0.94421768707483 |
| | 127.324766382975 | 0.941496598639456 |
| | 128.327323598589 | 0.936054421768708 |
| | 129.329880814203 | 0.927891156462585 |
| | 130.332438029817 | 0.922448979591837 |
| | 131.334995245431 | 0.915646258503401 |
| | 132.337552461045 | 0.912925170068027 |
| | 133.340109676659 | 0.907482993197279 |
| | 134.342666892273 | 0.917006802721088 |
| | 135.345224107887 | 0.934693877551021 |
| | 136.347781323501 | 0.940136054421769 |
| | 137.350338539115 | 0.937414965986395 |
| | 138.352895754729 | 0.945578231292517 |
| | 139.355452970343 | 0.949659863945578 |
| | 140.358010185957 | 0.953741496598639 |
| | 141.360567401571 | 0.956462585034014 |
| | 142.363124617185 | 0.955102040816327 |
| | 143.365681832799 | 0.956462585034014 |
| | 144.368239048413 | 0.959183673469388 |
| | 145.370796264027 | 0.959183673469388 |
| | 146.373353479641 | 0.960544217687075 |
| | 147.375910695255 | 0.960544217687075 |
| | 148.378467910869 | 0.959183673469388 |
| | 149.381025126483 | 0.959183673469388 |

| | | |
|---|---|---|
| | 150.383582342097 | 0.957823129251701 |
| | 151.386139557711 | 0.953741496598639 |
| | 152.388696773325 | 0.951020408163265 |
| | 153.391253988939 | 0.949659863945578 |
| | 154.393811204553 | 0.945578231292517 |
| | 155.396368420167 | 0.946938775510204 |
| | 156.398925635781 | 0.938775510204082 |
| | 157.401482851394 | 0.930612244897959 |
| | 158.404040067008 | 0.926530612244898 |
| | 159.406597282622 | 0.912925170068027 |
| | 160.409154498236 | 0.906122448979592 |
| | 161.41171171385 | 0.899319727891156 |
| | 162.414268929464 | 0.900680272108844 |
| | 163.416826145078 | 0.903401360544218 |
| | 164.419383360692 | 0.897959183673469 |
| | 165.421940576306 | 0.893877551020408 |
| | 166.42449779192 | 0.896598639455782 |
| | 167.427055007534 | 0.893877551020408 |
| | 168.429612223148 | 0.880272108843538 |
| | 169.432169438762 | 0.868027210884354 |
| | 170.434726654376 | 0.863945578231293 |
| | 171.43728386999 | 0.863945578231293 |
| | 172.439841085604 | 0.861224489795918 |
| | 173.442398301218 | 0.854421768707483 |
| | 174.444955516832 | 0.854421768707483 |
| | 175.447512732446 | 0.850340136054422 |
| | 176.45006994806 | 0.843537414965986 |
| | 177.452627163674 | 0.836734693877551 |
| | 178.455184379288 | 0.834013605442177 |
| | 179.457741594902 | 0.831292517006803 |
| | 180.460298810516 | 0.829931972789116 |
| | 181.46285602613 | 0.825850340136054 |
| | 182.465413241744 | 0.828571428571429 |
| | 183.467970457358 | 0.827210884353742 |
| | 184.470527672972 | 0.821768707482993 |
| | 185.473084888586 | 0.810884353741497 |
| | 186.4756421 | 0.805442176870748 |
| | 187.478199319814 | 0.806802721088435 |
| | 188.480756535428 | 0.795918367346939 |
| | 189.483313751042 | 0.79047619047619 |
| | 190.485870966656 | 0.786394557823129 |
| | 191.48842818227 | 0.783673469387755 |
| | 192.490985397884 | 0.787755102040816 |

| | | |
|---|---|---|
| | 193.493542613498 | 0.787755102040816 |
| | 194.496099829112 | 0.791836734693878 |
| | 195.498657044726 | 0.791836734693878 |
| | 196.50121426034 | 0.785034013605442 |
| | 197.503771475954 | 0.77687074829932 |
| | 198.506328691568 | 0.779591836734694 |
| | 199.508885907182 | 0.774149659863946 |
| | 200.511443122795 | 0.761904761904762 |
| | 201.514000338409 | 0.75374149659864 |
| | 202.516557554023 | 0.749659863945578 |
| | 203.519114769637 | 0.73469387755102 |
| | 204.521671985251 | 0.730612244897959 |
| | 205.524229200865 | 0.725170068027211 |
| | 206.526786416479 | 0.722448979591837 |
| | 207.529343632093 | 0.717006802721088 |
| | 208.531900847707 | 0.714285714285714 |
| | 209.534458063321 | 0.706122448979592 |
| | 210.537015278935 | 0.703401360544218 |
| | 211.539572494549 | 0.704761904761905 |
| | 212.542129710163 | 0.699319727891156 |
| | 213.544686925777 | 0.693877551020408 |
| | 214.547244141391 | 0.689795918367347 |
| | 215.549801357005 | 0.691156462585034 |
| | 216.552358572619 | 0.696598639455782 |
| | 217.554915788233 | 0.706122448979592 |
| | 218.557473003847 | 0.715646258503401 |
| | 219.560030219461 | 0.714285714285714 |
| | 220.562587435075 | 0.710204081632653 |
| | 221.565144650689 | 0.71156462585034 |
| | 222.567701866303 | 0.717006802721088 |
| | 223.570259081917 | 0.714285714285714 |
| | 224.572816297531 | 0.727891156462585 |
| | 225.575373513145 | 0.73469387755102 |
| | 226.577930728759 | 0.748299319727891 |
| | 227.580487944373 | 0.751020408163265 |
| | 228.583045159987 | 0.745578231292517 |
| | 229.585602375601 | 0.738775510204082 |
| | 230.588159591215 | 0.733333333333333 |
| | 231.590716806829 | 0.726530612244898 |
| | 232.593274022443 | 0.722448979591837 |
| | 233.595831238057 | 0.718367346938776 |
| | 234.598388453671 | 0.718367346938776 |
| | 235.600945669285 | 0.72108843537415 |

| | | |
|---|---|---|
| | 236.603502884899 | 0.718367346938776 |
| | 237.606060100513 | 0.717006802721088 |
| | 238.608617316127 | 0.717006802721088 |
| | 239.611174531741 | 0.67891156462585 |
| | 240.613731747355 | 0.659863945578231 |
| | 241.616288962969 | 0.636734693877551 |
| | 242.618846178583 | 0.605442176870748 |
| | 243.621403394197 | 0.57687074829932 |
| | 244.62396060981 | 0.549659863945578 |
| | 245.626517825424 | 0.515646258503401 |
| | 246.629075041038 | 0.496598639455782 |
| | 247.631632256652 | 0.473469387755102 |
| | 248.634189472266 | 0.436734693877551 |
| | 249.63674668788 | 0.404081632653061 |
| | 250.639303903494 | 0.378231292517007 |
| | 251.641861119108 | 0.371428571428571 |
| | 252.644418334722 | 0.342857142857143 |
| | 253.646975550336 | 0.302040816326531 |
| | 254.64953276595 | 0.292517006802721 |
| | 255.652089981564 | 0.300680272108844 |
| | 256.654647197178 | 0.289795918367347 |
| | 257.657204412792 | 0.282993197278912 |
| | 258.659761628406 | 0.270748299319728 |
| | 259.66231884402 | 0.259863945578231 |
| | 260.664876059634 | 0.23265306122449 |
| | 261.667433275248 | 0.23265306122449 |
| | 262.669990490862 | 0.236734693877551 |
| | 263.672547706476 | 0.238095238095238 |
| | 264.67510492209 | 0.239455782312925 |
| | 265.677662137704 | 0.235374149659864 |
| | 266.680219353318 | 0.229931972789116 |
| | 267.682776568932 | 0.224489795918367 |
| | 268.685333784546 | 0.224489795918367 |
| | 269.68789100016 | 0.221768707482993 |
| | 270.690448215774 | 0.217687074829932 |
| | 271.693005431388 | 0.216326530612245 |
| | 272.695562647002 | 0.214965986394558 |
| | 273.698119862616 | 0.214965986394558 |
| | 274.70067707823 | 0.213605442176871 |
| | 275.703234293844 | 0.212244897959184 |
| | 276.705791509458 | 0.210884353741497 |
| | 277.708348725072 | 0.210884353741497 |
| | 278.710905940686 | 0.20952380952381 |

| | | | | |
|---|---|---|---|---|
| | 279.7134632 | 0.202721088435374 | | |
| | 280.716020371914 | 0.201360544217687 | | |
| | 281.718577587528 | 0.195918367346939 | | |
| | 282.721134803142 | 0.197278911564626 | | |
| | 283.723692018756 | 0.186394557823129 | | |
| | 284.72624923437 | 0.186394557823129 | | |
| | 285.728806449984 | 0.185034013605442 | | |
| | 286.731363665598 | 0.185034013605442 | | |
| | 287.733920881212 | 0.186394557823129 | | |
| | 288.736478096826 | 0.193197278911565 | | |
| | 289.73903531244 | 0.201360544217687 | | |
| | 290.741592528053 | 0.206802721088435 | | |
| | 291.744149743667 | 0.201360544217687 | | |
| | 292.746706959281 | 0.195918367346939 | | |
| | 293.749264174895 | 0.193197278911565 | | |
| | 294.751821390509 | 0.182312925170068 | | |
| | 295.754378606123 | 0.142857142857143 | | |
| | 296.756935821737 | 0.129251700680272 | | |
| | 297.759493037351 | 0.12108843537415 | | |
| | 298.762050252965 | 0.115646258503401 | | |
| | 299.764607468579 | 0.106122448979592 | | |
| | 300.767164684193 | 0.0843537414965986 | | |
| Insect | 100.219379106929 | 0.6553778091846 | 1 Ma | Peris & Condamine (2024) |
| | 101.221572897998 | 0.653260556144434 | | |
| | 102.223766689068 | 0.649273784914708 | | |
| | 103.225960480137 | 0.647221511507487 | | |
| | 104.228154271206 | 0.643876414153106 | | |
| | 105.230348062275 | 0.636356375381925 | | |
| | 106.232541853345 | 0.628677948755437 | | |
| | 107.234735644414 | 0.61374210937426 | | |
| | 108.236929435483 | 0.601467727459175 | | |
| | 109.239123226553 | 0.592297476759628 | | |
| | 110.241317017622 | 0.586242187214445 | | |
| | 111.243510808691 | 0.588241664669897 | | |
| | 112.24570459976 | 0.611609965168209 | | |
| | 113.24789839083 | 0.624666810163356 | | |
| | 114.250092181899 | 0.636055844579548 | | |
| | 115.252285972968 | 0.644925564476718 | | |
| | 116.254479764038 | 0.654423420825701 | | |
| | 117.256673555107 | 0.659605546553169 | | |
| | 118.258867346176 | 0.666469020283122 | | |
| | 119.261061137245 | 0.669443192232768 | | |
| | 120.263254928315 | 0.6711231864929 | | |

| | | |
|---|---|---|
| | 121.265448719384 | 0.671550969076463 |
| | 122.267642510453 | 0.674269283721384 |
| | 123.269836301523 | 0.667067374403169 |
| | 124.272030092592 | 0.660199839446157 |
| | 125.274223883661 | 0.651078323471321 |
| | 126.27641767473 | 0.639498411383349 |
| | 127.2786115 | 0.624665456421003 |
| | 128.280805256869 | 0.612986721141259 |
| | 129.282999047938 | 0.597673187643581 |
| | 130.285192839008 | 0.574214186407615 |
| | 131.287386630077 | 0.54908602085034 |
| | 132.289580421146 | 0.530251403492385 |
| | 133.291774212215 | 0.51513010140884 |
| | 134.293968003285 | 0.501868841318383 |
| | 135.296161794354 | 0.487558430904314 |
| | 136.298355585423 | 0.47432966063033 |
| | 137.300549376493 | 0.463897722057743 |
| | 138.302743167562 | 0.457853262451384 |
| | 139.304936958631 | 0.455260845845297 |
| | 140.3071307 | 0.454669260437015 |
| | 141.30932454077 | 0.452438293039192 |
| | 142.311518331839 | 0.452882320530992 |
| | 143.313712122908 | 0.450697380373173 |
| | 144.315905913978 | 0.449556175569553 |
| | 145.318099705047 | 0.451819632783849 |
| | 146.320293496116 | 0.453740593182824 |
| | 147.322487287186 | 0.452752361265099 |
| | 148.324681078255 | 0.452707687767449 |
| | 149.326874869324 | 0.453671552322819 |
| | 150.329068660393 | 0.4549955123441 |
| | 151.331262451463 | 0.451582727872066 |
| | 152.333456242532 | 0.451768190574433 |
| | 153.335650033601 | 0.452026755363866 |
| | 154.337843824671 | 0.45943714100445 |
| | 155.34003761574 | 0.473227714354948 |
| | 156.342231406809 | 0.479120554817766 |
| | 157.344425197878 | 0.483051822611017 |
| | 158.346618988948 | 0.485465545226501 |
| | 159.348812780017 | 0.485613103142984 |
| | 160.351006571086 | 0.48446106840054 |
| | 161.353200362156 | 0.481501787616777 |
| | 162.355394153225 | 0.479272173961307 |
| | 163.357587944294 | 0.473812531051465 |

| | | |
|---|---|---|
| | 164.359781735363 | 0.464644987836625 |
| | 165.361975526433 | 0.43757961697214 |
| | 166.364169317502 | 0.400223096739783 |
| | 167.366363108571 | 0.37519781560134 |
| | 168.368556899641 | 0.342779747472902 |
| | 169.37075069071 | 0.314769464445988 |
| | 170.372944481779 | 0.297595888955223 |
| | 171.375138272848 | 0.285234867529542 |
| | 172.377332063918 | 0.276726596840636 |
| | 173.379525854987 | 0.273399098136844 |
| | 174.381719646056 | 0.270093259310702 |
| | 175.383913437126 | 0.268314441858797 |
| | 176.386107228195 | 0.267261230308125 |
| | 177.388301019264 | 0.266084828203327 |
| | 178.390494810334 | 0.267119087361055 |
| | 179.392688601403 | 0.271084198713132 |
| | 180.394882392472 | 0.277748672317187 |
| | 181.397076183541 | 0.284291309109468 |
| | 182.399269974611 | 0.285919861160184 |
| | 183.40146376568 | 0.284881540775397 |
| | 184.403657556749 | 0.274659432267532 |
| | 185.405851347819 | 0.27065235490251 |
| | 186.408045138888 | 0.2698482319448 |
| | 187.410238929957 | 0.268512088242342 |
| | 188.412432721026 | 0.264291119585538 |
| | 189.414626512096 | 0.261196464566471 |
| | 190.416820303165 | 0.257049951739085 |
| | 191.419014094234 | 0.25040307678562 |
| | 192.421207885304 | 0.246998414767705 |
| | 193.423401676373 | 0.244288222576903 |
| | 194.425595467442 | 0.241157016514303 |
| | 195.427789258511 | 0.237482959768131 |
| | 196.429983049581 | 0.234441100700833 |
| | 197.43217684065 | 0.229441730191027 |
| | 198.434370631719 | 0.225335829634232 |
| | 199.436564422789 | 0.221343643435094 |
| | 200.438758213858 | 0.215186823213433 |
| | 201.440952004927 | 0.211044371613106 |
| | 202.443145795996 | 0.204289197271397 |
| | 203.445339587066 | 0.198373343188578 |
| | 204.447533378135 | 0.196437491623719 |
| | 205.449727169204 | 0.192415523092814 |
| | 206.451920960274 | 0.188726575180758 |

| | | |
|---|---|---|
| | 207.454114751343 | 0.184654518182791 |
| | 208.456308542412 | 0.182435734466145 |
| | 209.458502333481 | 0.177707112426949 |
| | 210.460696124551 | 0.176036594363288 |
| | 211.46288991562 | 0.175556015827956 |
| | 212.465083706689 | 0.175856546630332 |
| | 213.467277497759 | 0.176170614856239 |
| | 214.469471288828 | 0.176911111923357 |
| | 215.471665079897 | 0.180151971116553 |
| | 216.473858870966 | 0.182797183674409 |
| | 217.476052662036 | 0.187537989394782 |
| | 218.478246453105 | 0.189759480596134 |
| | 219.480440244174 | 0.191382617677438 |
| | 220.482634035244 | 0.202039277480631 |
| | 221.484827826313 | 0.21658930029119 |
| | 222.487021617382 | 0.226102047806057 |
| | 223.489215408452 | 0.229627192893394 |
| | 224.491409199521 | 0.232484943000678 |
| | 225.49360299059 | 0.235502434705622 |
| | 226.495796781659 | 0.236852115831611 |
| | 227.497990572729 | 0.237125571786926 |
| | 228.500184363798 | 0.237890436216398 |
| | 229.502378154867 | 0.23783357903757 |
| | 230.504571945937 | 0.237125571786926 |
| | 231.506765737006 | 0.236246992999799 |
| | 232.508959528075 | 0.234726740337326 |
| | 233.511153319144 | 0.233355399333688 |
| | 234.513347110214 | 0.229452560129851 |
| | 235.515540901283 | 0.226290217993132 |
| | 236.517734692352 | 0.218093308045427 |
| | 237.519928483422 | 0.207215988238687 |
| | 238.522122274491 | 0.202488719941843 |
| | 239.52431606556 | 0.196909947704933 |
| | 240.526509856629 | 0.189961188206738 |
| | 241.528703647699 | 0.185145926656947 |
| | 242.530897438768 | 0.179152909260004 |
| | 243.533091229837 | 0.170773244094638 |
| | 244.535285020907 | 0.16350770888583 |
| | 245.537478811976 | 0.159290801456085 |
| | 246.539672603045 | 0.153228743199137 |
| | 247.541866394114 | 0.143997574093703 |
| | 248.544060185184 | 0.137346637913179 |
| | 249.546253976253 | 0.133918962275262 |

| | | |
|---|---|---|
| | 250.548447767322 | 0.135275412113016 |
| | 251.550641558392 | 0.149546563998846 |
| | 252.552835349461 | 0.171995673439439 |
| | 253.55502914053 | 0.177058669839839 |
| | 254.557222931599 | 0.171367536987626 |
| | 255.559416722669 | 0.162013177328065 |
| | 256.561610513738 | 0.161678802966862 |
| | 257.563804304807 | 0.159045774090184 |
| | 258.565998095877 | 0.155683078085213 |
| | 259.568191886946 | 0.153866355847422 |
| | 260.570385678015 | 0.152574885642615 |
| | 261.572579469085 | 0.15484240408397 |
| | 262.574773260154 | 0.15765412695125 |
| | 263.576967051223 | 0.160561965525597 |
| | 264.579160842292 | 0.16543543799657 |
| | 265.581354633362 | 0.170958706797005 |
| | 266.583548424431 | 0.177016703826894 |
| | 267.5857422 | 0.190569018523256 |
| | 268.58793600657 | 0.204897027587915 |
| | 269.590129797639 | 0.213921074113332 |
| | 270.592323588708 | 0.222932936957572 |
| | 271.594517379777 | 0.229276573623955 |
| | 272.596711170847 | 0.233872528912552 |
| | 273.598904961916 | 0.233720909769011 |
| | 274.601098752985 | 0.227530245988523 |
| | 275.603292544055 | 0.223945536237652 |
| | 276.605486335124 | 0.216567640413542 |
| | 277.607680126193 | 0.208242124942297 |
| | 278.609873917262 | 0.196885580342579 |
| | 279.612067708332 | 0.187145404112399 |
| | 280.614261499401 | 0.173594443158389 |
| | 281.61645529047 | 0.163798763491735 |
| | 282.61864908154 | 0.157713691614785 |
| | 283.620842872609 | 0.154674540032192 |
| | 284.623036663678 | 0.151541980227239 |
| | 285.625230454747 | 0.149033495647041 |
| | 286.627424245817 | 0.147816481271652 |
| | 287.629618036886 | 0.145000697177312 |
| | 288.631811827955 | 0.143595512614848 |
| | 289.634005619025 | 0.140976021161701 |
| | 290.636199410094 | 0.135137330393005 |
| | 291.638393201163 | 0.126894393205297 |
| | 292.640586992232 | 0.124349357581566 |

| | | | | |
|---|---|---|---|---|
| | 293.642780783302 | 0.123948649845064 | | |
| | 294.644974574371 | 0.120849933598938 | | |
| | 295.64716836544 | 0.11585868554325 | | |
| | 296.64936215651 | 0.111764968667633 | | |
| | 297.651555947579 | 0.112815472733598 | | |
| | 298.653749738648 | 0.123399030449727 | | |
| | 299.655943529717 | 0.134046214056448 | | |
| | 300.658137320787 | 0.140992266069937 | | |
| Pollinator | 100.242158447329 | 0.593119810201661 | 1 Ma | Peris & Condamine (2024) |
| | 101.244580031802 | 0.591340450771056 | | |
| | 102.247001616275 | 0.588374851720047 | | |
| | 103.249423200749 | 0.586632678484439 | | |
| | 104.251844785222 | 0.58306280376473 | | |
| | 105.254266369695 | 0.574154850753206 | | |
| | 106.256687954169 | 0.567626814221488 | | |
| | 107.259109538642 | 0.563456382452578 | | |
| | 108.261531123115 | 0.556902315584362 | | |
| | 109.263952707588 | 0.554537273582555 | | |
| | 110.266374292062 | 0.548001799811838 | | |
| | 111.268795876535 | 0.542111506524318 | | |
| | 112.271217461008 | 0.538022884384401 | | |
| | 113.273639045482 | 0.530693485350498 | | |
| | 114.276060629955 | 0.530249110320285 | | |
| | 115.278482214428 | 0.529159554806873 | | |
| | 116.280903798901 | 0.53202846975089 | | |
| | 117.283325383375 | 0.532978577033062 | | |
| | 118.285746967848 | 0.536773428232503 | | |
| | 119.288168552321 | 0.53202846975089 | | |
| | 120.290590136795 | 0.529503527110595 | | |
| | 121.293011721268 | 0.526199533685115 | | |
| | 122.295433305741 | 0.523131672597865 | | |
| | 123.297854890214 | 0.516828612545878 | | |
| | 124.300276474688 | 0.513048635824436 | | |
| | 125.302698059161 | 0.504120230405664 | | |
| | 126.305119643634 | 0.491696322657177 | | |
| | 127.307541228108 | 0.478186578014778 | | |
| | 128.309962812581 | 0.470102299222437 | | |
| | 129.312384397054 | 0.463732304019456 | | |
| | 130.314805981527 | 0.44147636631377 | | |
| | 131.317227566001 | 0.425860023724792 | | |
| | 132.319649150474 | 0.414631652145458 | | |
| | 133.322070734947 | 0.40539646061796 | | |
| | 134.324492319421 | 0.39558674237776 | | |

| | | |
|---|---|---|
| | 135.326913903894 | 0.384888274077132 |
| | 136.329335488367 | 0.373334523291573 |
| | 137.33175707284 | 0.362898217665674 |
| | 138.334178657314 | 0.359430604982206 |
| | 139.336600241787 | 0.355278766310795 |
| | 140.33902182626 | 0.352313167259787 |
| | 141.341443410734 | 0.347568208778173 |
| | 142.343864995207 | 0.343213705344028 |
| | 143.34628657968 | 0.335889884239375 |
| | 144.348708164153 | 0.327920882651525 |
| | 145.351129748627 | 0.323412056508142 |
| | 146.3535513 | 0.320435301598635 |
| | 147.355972917573 | 0.311806245049588 |
| | 148.358394502047 | 0.306306406809536 |
| | 149.36081608652 | 0.301539136610925 |
| | 150.363237670993 | 0.29300118623962 |
| | 151.365659255466 | 0.284697508896797 |
| | 152.36808083994 | 0.279952550415184 |
| | 153.370502424413 | 0.276986951364176 |
| | 154.372924008886 | 0.276986951364176 |
| | 155.37534559336 | 0.276393831553974 |
| | 156.377767177833 | 0.276393831553974 |
| | 157.380188762306 | 0.275207591933571 |
| | 158.38261034678 | 0.273428232502966 |
| | 159.385031931253 | 0.27165445100161 |
| | 160.387453515726 | 0.270462633451957 |
| | 161.389875100199 | 0.270462633451957 |
| | 162.392296684673 | 0.266903914590747 |
| | 163.394718269146 | 0.264531435349941 |
| | 164.397139853619 | 0.259193357058126 |
| | 165.399561438093 | 0.244835767169796 |
| | 166.401983022566 | 0.229392340387554 |
| | 167.404404607039 | 0.217256625650294 |
| | 168.406826191512 | 0.193167408531257 |
| | 169.409247775986 | 0.170119404872135 |
| | 170.411669360459 | 0.158721836105564 |
| | 171.414090944932 | 0.14920960742534 |
| | 172.416512529406 | 0.146361144888572 |
| | 173.418934113879 | 0.145314353499407 |
| | 174.421355698352 | 0.144128113879004 |
| | 175.423777282825 | 0.1429418742586 |
| | 176.426198867299 | 0.1429418742586 |
| | 177.428620451772 | 0.142348754448399 |

| | | |
|---|---|---|
| | 178.431042036245 | 0.142348754448399 |
| | 179.433463620719 | 0.142348754448399 |
| | 180.435885205192 | 0.14257744954763 |
| | 181.438306789665 | 0.148279952550415 |
| | 182.440728374138 | 0.148279952550415 |
| | 183.443149958612 | 0.148279952550415 |
| | 184.445571543085 | 0.147093712930012 |
| | 185.447993127558 | 0.142348754448399 |
| | 186.450414712032 | 0.142348754448399 |
| | 187.452836296505 | 0.142348754448399 |
| | 188.455257880978 | 0.142348754448399 |
| | 189.457679465451 | 0.142348754448399 |
| | 190.460101049925 | 0.142348754448399 |
| | 191.462522634398 | 0.142348754448399 |
| | 192.464944218871 | 0.141393069236977 |
| | 193.467365803345 | 0.140569395017794 |
| | 194.469787387818 | 0.139976275207592 |
| | 195.472208972291 | 0.138196915776987 |
| | 196.474630556764 | 0.137010676156584 |
| | 197.477052141238 | 0.133626732411859 |
| | 198.479473725711 | 0.131836217122755 |
| | 199.481895310184 | 0.12945258202345 |
| | 200.484316894658 | 0.128255186544547 |
| | 201.486738479131 | 0.124815463507328 |
| | 202.489160063604 | 0.119217081850534 |
| | 203.491581648077 | 0.113393723714008 |
| | 204.494003232551 | 0.110913404507711 |
| | 205.496424817024 | 0.106761565836299 |
| | 206.498846401497 | 0.103795966785291 |
| | 207.501267985971 | 0.103795966785291 |
| | 208.503689570444 | 0.102609727164887 |
| | 209.506111154917 | 0.0996850329283757 |
| | 210.50853273939 | 0.0978945176392716 |
| | 211.510954323864 | 0.0966785290628707 |
| | 212.513375908337 | 0.0949066068712651 |
| | 213.51579749281 | 0.0943060498220641 |
| | 214.518219077284 | 0.0937129300118624 |
| | 215.520640661757 | 0.0936868996753645 |
| | 216.52306224623 | 0.0925266903914591 |
| | 217.525483830704 | 0.0925266903914591 |
| | 218.527905415177 | 0.0918740726692624 |
| | 219.53032699965 | 0.0901542111506524 |
| | 220.532748584123 | 0.0889679715302491 |

| | | |
|---|---|---|
| | 221.535170168597 | 0.0889679715302491 |
| | 222.53759175307 | 0.0889679715302491 |
| | 223.540013337543 | 0.0883748517200475 |
| | 224.542434922017 | 0.0871886120996441 |
| | 225.54485650649 | 0.0857271946362635 |
| | 226.547278090963 | 0.0822949088380429 |
| | 227.549699675436 | 0.0782918149466192 |
| | 228.55212125991 | 0.0771055753262159 |
| | 229.554542844383 | 0.0757371233503275 |
| | 230.556964428856 | 0.0735468564650059 |
| | 231.55938601333 | 0.0723606168446026 |
| | 232.561807597803 | 0.0703655774830153 |
| | 233.564229182276 | 0.0693950177935943 |
| | 234.566650766749 | 0.0670225385527877 |
| | 235.569072351223 | 0.0652431791221827 |
| | 236.571493935696 | 0.0652431791221827 |
| | 237.573915520169 | 0.0605484220038152 |
| | 238.576337104643 | 0.0587188612099644 |
| | 239.578758689116 | 0.0587188612099644 |
| | 240.581180273589 | 0.0587188612099644 |
| | 241.583601858062 | 0.0587188612099644 |
| | 242.586023442536 | 0.0575326215895611 |
| | 243.588445027009 | 0.0545670225385528 |
| | 244.590866611482 | 0.0500656336341696 |
| | 245.593288195956 | 0.0470888787246623 |
| | 246.595709780429 | 0.0375878059029367 |
| | 247.598131364902 | 0.0302491103202847 |
| | 248.600552949375 | 0.029655990510083 |
| | 249.602974533849 | 0.0290628706998814 |
| | 250.605396118322 | 0.0284697508896797 |
| | 251.607817702795 | 0.0396702328227665 |
| | 252.610239287269 | 0.0527876631079478 |
| | 253.612660871742 | 0.0545670225385528 |
| | 254.615082456215 | 0.0535127939103887 |
| | 255.617504040688 | 0.0497012089231993 |
| | 256.619925625162 | 0.0492289442467378 |
| | 257.622347209635 | 0.0481412480430767 |
| | 258.624768794108 | 0.0463507327539723 |
| | 259.627190378582 | 0.0421877382240619 |
| | 260.629611963055 | 0.0409252669039146 |
| | 261.632033547528 | 0.0409252669039146 |
| | 262.634455132001 | 0.0409252669039146 |
| | 263.636876716475 | 0.0409252669039146 |

| | | | | |
|---|---|---|---|---|
| | 264.639298300948 | 0.0409252669039146 | | |
| | 265.641719885421 | 0.0409252669039146 | | |
| | 266.644141469895 | 0.0409252669039146 | | |
| | 267.646563054368 | 0.0415183867141162 | | |
| | 268.648984638841 | 0.0421115065243179 | | |
| | 269.651406223314 | 0.0427046263345196 | | |
| | 270.653827807788 | 0.0456702253855279 | | |
| | 271.656249392261 | 0.0462633451957295 | | |
| | 272.658670976734 | 0.0468564650059312 | | |
| | 273.661092561208 | 0.0468564650059312 | | |
| | 274.663514145681 | 0.0468564650059312 | | |
| | 275.665935730154 | 0.0468564650059312 | | |
| | 276.668357314627 | 0.0468564650059312 | | |
| | 277.670778899101 | 0.0444839857651246 | | |
| | 278.673200483574 | 0.0427046263345196 | | |
| | 279.675622068047 | 0.0400383761532371 | | |
| | 280.678043652521 | 0.0361803084223013 | | |
| | 281.680465236994 | 0.034994068801898 | | |
| | 282.682886821467 | 0.034994068801898 | | |
| | 283.68530840594 | 0.034994068801898 | | |
| | 284.687729990414 | 0.034994068801898 | | |
| | 285.690151574887 | 0.034994068801898 | | |
| | 286.69257315936 | 0.034994068801898 | | |
| | 287.694994743834 | 0.0344009489916963 | | |
| | 288.697416328307 | 0.033214709371293 | | |
| | 289.69983791278 | 0.0302491103202847 | | |
| | 290.702259497254 | 0.0290628706998814 | | |
| | 291.704681081727 | 0.0290628706998814 | | |
| | 292.7071027 | 0.0290628706998814 | | |
| | 293.709524250673 | 0.0290628706998814 | | |
| | 294.711945835147 | 0.0290628706998814 | | |
| | 295.71436741962 | 0.0290628706998814 | | |
| | 296.716789004093 | 0.0290628706998814 | | |
| | 297.719210588567 | 0.0290628706998814 | | |
| | 298.72163217304 | 0.0290628706998814 | | |
| | 299.724053757513 | 0.0290628706998814 | | |
| | 300.726475341986 | 0.0290628706998814 | | |
| Pteridophyta | 100.024094985104 | 0.936418911006099 | 1 Ma | This study |
| | 101.024335934955 | 0.926154401268583 | | |
| | 102.024576884806 | 0.918551764084028 | | |
| | 103.024817834657 | 0.928863750739849 | | |
| | 104.025058784508 | 0.933918459975375 | | |
| | 105.025299734359 | 0.940755136211279 | | |

| | | |
|---|---|---|
| | 106.02554068421 | 0.95132982848172 |
| | 107.025781634061 | 0.956083850567982 |
| | 108.026022583912 | 0.96198997920511 |
| | 109.026263533763 | 0.968234776526146 |
| | 110.026504483614 | 0.973783245712043 |
| | 111.026745433465 | 0.980373041972761 |
| | 112.026986383316 | 0.993929284712749 |
| | 113.027227333167 | 1 |
| | 114.027468283018 | 0.9950117584501 |
| | 115.027709232869 | 0.985019449710866 |
| | 116.02795018272 | 0.979293733363297 |
| | 117.028191132571 | 0.974666316392514 |
| | 118.028432082422 | 0.970921970102202 |
| | 119.028673032273 | 0.963306672406098 |
| | 120.028913982124 | 0.955172293736528 |
| | 121.029154931975 | 0.945452185995575 |
| | 122.029395881826 | 0.932190300149078 |
| | 123.029636831678 | 0.926806417613304 |
| | 124.029877781529 | 0.911268804816059 |
| | 125.03011873138 | 0.903849745048949 |
| | 126.030359681231 | 0.896357887340438 |
| | 127.030600631082 | 0.881766647781404 |
| | 128.030841580933 | 0.867552058440921 |
| | 129.031082530784 | 0.847576936346113 |
| | 130.031323480635 | 0.833852941828115 |
| | 131.031564430486 | 0.818885052050528 |
| | 132.031805380337 | 0.80140405073067 |
| | 133.032046330188 | 0.77561142357957 |
| | 134.032287280039 | 0.757680974099758 |
| | 135.03252822989 | 0.724102132346659 |
| | 136.032769179741 | 0.686149083853733 |
| | 137.033010129592 | 0.656036057136889 |
| | 138.033251079443 | 0.638548725561256 |
| | 139.033492029294 | 0.628562747077796 |
| | 140.033732979145 | 0.62063410172088 |
| | 141.033973928996 | 0.608948449562105 |
| | 142.034214878847 | 0.594404686921375 |
| | 143.034455828698 | 0.590904055478362 |
| | 144.034696778549 | 0.59010644325084 |
| | 145.0349377 | 0.587605992220116 |
| | 146.035178678251 | 0.581816973314807 |
| | 147.035419628102 | 0.579784961211358 |
| | 148.035660577953 | 0.57573043238812 |

| | | |
|---|---|---|
| | 149.035901527804 | 0.578075792152382 |
| | 150.036142477656 | 0.580880095460257 |
| | 151.036383427507 | 0.582127155847732 |
| | 152.036624377358 | 0.581835964082129 |
| | 153.036865327209 | 0.581354864643306 |
| | 154.03710627706 | 0.579341843307179 |
| | 155.037347226911 | 0.578854413612582 |
| | 156.037588176762 | 0.576847722532229 |
| | 157.037829126613 | 0.571096685161564 |
| | 158.038070076464 | 0.566602203562035 |
| | 159.038311026315 | 0.563120562886344 |
| | 160.038551976166 | 0.557600579851429 |
| | 161.038792926017 | 0.555609714410511 |
| | 162.039033875868 | 0.550099226759257 |
| | 163.039274825719 | 0.551865368120199 |
| | 164.03951577557 | 0.547174648591676 |
| | 165.039756725421 | 0.543120119768439 |
| | 166.039997675272 | 0.551023444102259 |
| | 167.040238625123 | 0.554571552463577 |
| | 168.040479574974 | 0.553131419274996 |
| | 169.040720524825 | 0.5504537210826 |
| | 170.040961474676 | 0.546174468179387 |
| | 171.041202424527 | 0.542971358757751 |
| | 172.041443374378 | 0.543465118708122 |
| | 173.041684324229 | 0.55080188515017 |
| | 174.04192527408 | 0.553998664316032 |
| | 175.042166223931 | 0.557743010606344 |
| | 176.042407173782 | 0.558850805366791 |
| | 177.042648123634 | 0.562041254276879 |
| | 178.042889073485 | 0.562601481912877 |
| | 179.043130023336 | 0.563851707428239 |
| | 180.043370973187 | 0.564532209923942 |
| | 181.043611923038 | 0.566456607679233 |
| | 182.043852872889 | 0.566928211734395 |
| | 183.04409382274 | 0.563180700316196 |
| | 184.044334772591 | 0.564519549412394 |
| | 185.044575722442 | 0.564437256087332 |
| | 186.044816672293 | 0.561939970184495 |
| | 187.045057622144 | 0.561351256397515 |
| | 188.045298571995 | 0.559385711979692 |
| | 189.045539521846 | 0.555698337991347 |
| | 190.045780471697 | 0.546902447593395 |
| | 191.046021421548 | 0.545098324697809 |

| | | |
|---|---|---|
| | 192.046262371399 | 0.545098324697809 |
| | 193.04650332125 | 0.546348550213171 |
| | 194.046744271101 | 0.547598775728533 |
| | 195.046985220952 | 0.549482026821294 |
| | 196.047226170803 | 0.550089731375596 |
| | 197.047467120654 | 0.541968013217574 |
| | 198.047708070505 | 0.538847197 |
| | 199.047949020356 | 0.533865285826874 |
| | 200.048189970207 | 0.525094716452018 |
| | 201.048430920058 | 0.519479779580494 |
| | 202.048671869909 | 0.518204233042036 |
| | 203.04891281976 | 0.519451293429511 |
| | 204.049153769612 | 0.519530421626686 |
| | 205.049394719463 | 0.519492440092042 |
| | 206.049635669314 | 0.516995154189205 |
| | 207.049876619165 | 0.517533225929994 |
| | 208.050117569016 | 0.516409605530112 |
| | 209.050358518867 | 0.517593363359847 |
| | 210.050599468718 | 0.517007814700753 |
| | 211.050840418569 | 0.513842686813761 |
| | 212.05108136842 | 0.515580342023719 |
| | 213.051322318271 | 0.518843588875209 |
| | 214.051563268122 | 0.517697812580117 |
| | 215.051804217973 | 0.515092912329123 |
| | 216.052045167824 | 0.516792586004438 |
| | 217.052286117675 | 0.519406981639093 |
| | 218.052527067526 | 0.521214269662566 |
| | 219.052768017377 | 0.522594265421295 |
| | 220.053008967228 | 0.524255957561966 |
| | 221.053249917079 | 0.531197083018139 |
| | 222.05349086693 | 0.536894313214725 |
| | 223.053731816781 | 0.540803246155161 |
| | 224.053972766632 | 0.542430121889075 |
| | 225.054213716483 | 0.543848099182447 |
| | 226.054454666334 | 0.543132780279987 |
| | 227.054695616185 | 0.547598775728533 |
| | 228.054936566036 | 0.549906153958151 |
| | 229.055177515887 | 0.553650500248463 |
| | 230.055418465738 | 0.557917092640128 |
| | 231.055659415589 | 0.560101030882153 |
| | 232.055900365441 | 0.561132862573312 |
| | 233.056141315292 | 0.558338054649098 |
| | 234.056382265143 | 0.557600579851429 |

| | | |
|---|---|---|
| | 235.056623214994 | 0.557600579851429 |
| | 236.056864164845 | 0.557097324517397 |
| | 237.057105114696 | 0.55985098577908 |
| | 238.057346064547 | 0.560341580601564 |
| | 239.057587014398 | 0.563357947477868 |
| | 240.057827964249 | 0.55916731815549 |
| | 241.0580689 | 0.546899282465508 |
| | 242.058309863951 | 0.531424972226003 |
| | 243.058550813802 | 0.509224765226639 |
| | 244.058791763653 | 0.491632984430736 |
| | 245.059032713504 | 0.485214105075916 |
| | 246.059273663355 | 0.478671785733502 |
| | 247.059514613206 | 0.465087056842532 |
| | 248.059755563057 | 0.44589372133581 |
| | 249.059996512908 | 0.432578028315234 |
| | 250.060237462759 | 0.430995464371738 |
| | 251.06047841261 | 0.435768477225323 |
| | 252.060719362461 | 0.467691957093526 |
| | 253.060960312312 | 0.441035250029277 |
| | 254.061201262163 | 0.423114295933127 |
| | 255.061442212014 | 0.409266861427536 |
| | 256.061683161865 | 0.407573518007995 |
| | 257.061924111716 | 0.404199491680461 |
| | 258.062165061567 | 0.398771297354269 |
| | 259.062406011418 | 0.393821037339014 |
| | 260.06264696127 | 0.393821037339014 |
| | 261.062887911121 | 0.38707931493972 |
| | 262.063128860972 | 0.37881833115467 |
| | 263.063369810823 | 0.376317880123946 |
| | 264.063610760674 | 0.372988165586831 |
| | 265.063851710525 | 0.367993593781157 |
| | 266.064092660376 | 0.365907774503629 |
| | 267.064333610227 | 0.36216026308543 |
| | 268.064574560078 | 0.359662977182593 |
| | 269.064815509929 | 0.356314271878155 |
| | 270.06505645978 | 0.356314271878155 |
| | 271.065297409631 | 0.356314271878155 |
| | 272.065538359482 | 0.354285424902593 |
| | 273.065779309333 | 0.353427675245218 |
| | 274.066020259184 | 0.350547408868055 |
| | 275.066261209035 | 0.346803062577743 |
| | 276.066502158886 | 0.345062242239898 |
| | 277.066743108737 | 0.345062242239898 |

| | | | | |
|---|---|---|---|---|
| | 278.066984058588 | 0.345062242239898 | | |
| | 279.067225008439 | 0.345062242239898 | | |
| | 280.06746595829 | 0.345062242239898 | | |
| | 281.067706908141 | 0.345062242239898 | | |
| | 282.067947857992 | 0.348278012173082 | | |
| | 283.068188807843 | 0.352563595332069 | | |
| | 284.068429757694 | 0.354165150042888 | | |
| | 285.068670707545 | 0.356314271878155 | | |
| | 286.068911657396 | 0.357349268697202 | | |
| | 287.069152607248 | 0.361998841563193 | | |
| | 288.069393557099 | 0.353370702943252 | | |
| | 289.06963450695 | 0.343141009612493 | | |
| | 290.069875456801 | 0.333731084404465 | | |
| | 291.070116406652 | 0.324729460693859 | | |
| | 292.070357356503 | 0.3206559411033 | | |
| | 293.070598306354 | 0.311910692751541 | | |
| | 294.070839256205 | 0.304067505847574 | | |
| | 295.071080206056 | 0.291603232228598 | | |
| | 296.071321155907 | 0.287238520872436 | | |
| | 297.071562105758 | 0.283810687370823 | | |
| | 298.071803055609 | 0.275992821489952 | | |
| | 299.07204400546 | 0.271298936833543 | | |
| | 300.072284955311 | 0.266298034772095 | | |
| Sediment | 103 | 4.41089230860165 | variable | This study |
| | 107 | 3.69563920688673 | | |
| | 111 | 3.2358614010072 | | |
| | 116 | 2.89046758323695 | | |
| | 122 | 2.92304158640879 | | |
| | 127 | 2.35077737809747 | | |
| | 131 | 2.46993916427186 | | |
| | 136 | 2.36699036706338 | | |
| | 142 | 3.06508504762453 | | |
| | 145 | 2.79870895704394 | | |
| | 149 | 3.13080348767026 | | |
| | 155 | 2.53500419922735 | | |
| | 160 | 2.82300295983632 | | |
| | 165 | 2.24398083195091 | | |
| | 168 | 2.11681867049636 | | |
| | 172 | 2.09625120194591 | | |
| | 178 | 1.8225086234074 | | |
| | 187 | 1.82695813833921 | | |
| | 191 | 1.90399800077774 | | |
| | 196 | 1.68230529481523 | | |

| | | | | |
|---|---|---|---|---|
| | 201 | 2.0968987907075 | | |
| | 205 | 1.70254429709202 | | |
| | 213 | 1.37060178543318 | | |
| | 218 | 1.56714191213654 | | |
| | 222 | 1.80519685636678 | | |
| | 227 | 1.68335673196955 | | |
| | 232 | 1.67238144516299 | | |
| | 234 | 1.88875281365819 | | |
| | 240 | 0.521857754627427 | | |
| | 245 | 1.82186979271779 | | |
| | 252 | 1.69712744702531 | | |
| | 256 | 1.75455147006793 | | |
| | 263 | 1.67792716630001 | | |
| | 265 | 1.73635699638648 | | |
| | 269 | 2.01018191492865 | | |
| | 275 | 2.56503213343052 | | |
| | 280 | 2.71661187309353 | | |
| | 287 | 2.51249952804069 | | |
| | 293 | 2.56754579264833 | | |
| | 297 | 2.69035073920729 | | |
| Temperature | 100 | 23.8426154231428 | 0.5 Ma | Peris & Condamine (2024) |
| | 100.5 | 23.4980260249617 | | |
| | 101 | 23.1536404071147 | | |
| | 101.5 | 22.8145842396795 | | |
| | 102 | 22.4845006871027 | | |
| | 102.5 | 22.1668051973073 | | |
| | 103 | 21.8649132182161 | | |
| | 103.5 | 21.5822401977518 | | |
| | 104 | 21.3222015838376 | | |
| | 104.5 | 21.0882066096289 | | |
| | 105 | 20.8823469776193 | | |
| | 105.5 | 20.7037748053822 | | |
| | 106 | 20.5512444653852 | | |
| | 106.5 | 20.423510330096 | | |
| | 107 | 20.3193267719824 | | |
| | 107.5 | 20.237448163512 | | |
| | 108 | 20.1765696680054 | | |
| | 108.5 | 20.1345005045045 | | |
| | 109 | 20.1083678903911 | | |
| | 109.5 | 20.0952814995964 | | |
| | 110 | 20.0923510060513 | | |
| | 110.5 | 20.0966860836867 | | |
| | 111 | 20.1054487107786 | | |

| | | |
|---|---|---|
| | 111.5 | 20.1161874274033 |
| | 112 | 20.1266240317797 |
| | 112.5 | 20.1344811393825 |
| | 113 | 20.137481365686 |
| | 113.5 | 20.1333473261648 |
| | 114 | 20.1198016362934 |
| | 114.5 | 20.0945669115462 |
| | 115 | 20.0553657673979 |
| | 115.5 | 20.0004453618623 |
| | 116 | 19.9301510231116 |
| | 116.5 | 19.8453526218573 |
| | 117 | 19.7469200288108 |
| | 117.5 | 19.6357231146838 |
| | 118 | 19.5126317501878 |
| | 118.5 | 19.3785158060343 |
| | 119 | 19.2342451529349 |
| | 119.5 | 19.080689661601 |
| | 120 | 18.9187192027444 |
| | 120.5 | 18.7492036470764 |
| | 121 | 18.5730128653087 |
| | 121.5 | 18.3911189071187 |
| | 122 | 18.2056589532954 |
| | 122.5 | 18.0195112421022 |
| | 123 | 17.8355658934106 |
| | 123.5 | 17.656713027092 |
| | 124 | 17.4858426821414 |
| | 124.5 | 17.3256594142951 |
| | 125 | 17.1782867225006 |
| | 125.5 | 17.0457346287223 |
| | 126 | 16.9300131549245 |
| | 126.5 | 16.8331323230718 |
| | 127 | 16.7571021551285 |
| | 127.5 | 16.7039326085554 |
| | 128 | 16.6753097214662 |
| | 128.5 | 16.6718429780052 |
| | 129 | 16.6939184960176 |
| | 129.5 | 16.7419223933488 |
| | 130 | 16.816062312464 |
| | 130.5 | 16.9152268512212 |
| | 131 | 17.0377134077817 |
| | 131.5 | 17.1818165916287 |
| | 132 | 17.3458310122457 |
| | 132.5 | 17.5279207578679 |

| | | |
|---|---|---|
| | 133 | 17.7257278317387 |
| | 133.5 | 17.9367637158534 |
| | 134 | 18.1585398922072 |
| | 134.5 | 18.3885679762932 |
| | 135 | 18.624510692425 |
| | 135.5 | 18.8644719581722 |
| | 136 | 19.1066357793914 |
| | 136.5 | 19.3491861619388 |
| | 137 | 19.590307111671 |
| | 137.5 | 19.8281932198435 |
| | 138 | 20.061450585098 |
| | 138.5 | 20.2892198687262 |
| | 139 | 20.5106774577414 |
| | 139.5 | 20.7249997391569 |
| | 140 | 20.931363099986 |
| | 140.5 | 21.1289439272421 |
| | 141 | 21.3169186079384 |
| | 141.5 | 21.4944635290883 |
| | 142 | 21.6607550777051 |
| | 142.5 | 21.8149744556908 |
| | 143 | 21.9563573742288 |
| | 143.5 | 22.0841740289626 |
| | 144 | 22.1976951603639 |
| | 144.5 | 22.2961915089041 |
| | 145 | 22.3789338150548 |
| | 145.5 | 22.4451928192876 |
| | 146 | 22.4942392620739 |
| | 146.5 | 22.5253438838853 |
| | 147 | 22.5377859550726 |
| | 147.5 | 22.5308992454769 |
| | 148 | 22.5040389687023 |
| | 148.5 | 22.4565959886067 |
| | 149 | 22.3882669101344 |
| | 149.5 | 22.2989040410563 |
| | 150 | 22.1884472856952 |
| | 150.5 | 22.0578576807608 |
| | 151 | 21.9087633156794 |
| | 151.5 | 21.7428037743748 |
| | 152 | 21.5616186407709 |
| | 152.5 | 21.3668474987914 |
| | 153 | 21.1601944511201 |
| | 153.5 | 20.9440887844827 |
| | 154 | 20.7214161121828 |

| | | |
|---|---|---|
| | 154.5 | 20.4950691511644 |
| | 155 | 20.2679406183715 |
| | 155.5 | 20.0429161518057 |
| | 156 | 19.8227720374202 |
| | 156.5 | 19.6101985921945 |
| | 157 | 19.4078838158003 |
| | 157.5 | 19.2185088243042 |
| | 158 | 19.0442221466923 |
| | 158.5 | 18.8862695398816 |
| | 159 | 18.7458093135087 |
| | 159.5 | 18.6239984096828 |
| | 160 | 18.521703854629 |
| | 160.5 | 18.4391458339456 |
| | 161 | 18.3764570114542 |
| | 161.5 | 18.3336975044875 |
| | 162 | 18.3104918486234 |
| | 162.5 | 18.3063027617105 |
| | 163 | 18.3205926928298 |
| | 163.5 | 18.3528240910624 |
| | 164 | 18.4024594054891 |
| | 164.5 | 18.468961085191 |
| | 165 | 18.5517915792491 |
| | 165.5 | 18.6504429726374 |
| | 166 | 18.764525893903 |
| | 166.5 | 18.8936806074858 |
| | 167 | 19.0375473778261 |
| | 167.5 | 19.1957664693638 |
| | 168 | 19.3679781465391 |
| | 168.5 | 19.553822673792 |
| | 169 | 19.7529403155626 |
| | 169.5 | 19.9649713362911 |
| | 170 | 20.1895560004176 |
| | 170.5 | 20.426334572382 |
| | 171 | 20.6749473166245 |
| | 171.5 | 20.9349655665165 |
| | 172 | 21.2056849311539 |
| | 172.5 | 21.4863320885643 |
| | 173 | 21.776133716775 |
| | 173.5 | 22.0743164938132 |
| | 174 | 22.3801070977065 |
| | 174.5 | 22.6927322064821 |
| | 175 | 23.0114184981674 |
| | 175.5 | 23.3353926507897 |

| | | |
|---|---|---|
| | 176 | 23.6638813423764 |
| | 176.5 | 23.996111250955 |
| | 177 | 24.3311925502323 |
| | 177.5 | 24.6675444958434 |
| | 178 | 25.0033326165696 |
| | 178.5 | 25.3367220524922 |
| | 179 | 25.6658779436929 |
| | 179.5 | 25.988965430253 |
| | 180 | 26.3041496522541 |
| | 180.5 | 26.6095957497777 |
| | 181 | 26.9034688629051 |
| | 181.5 | 27.1839340435464 |
| | 182 | 27.4485654670755 |
| | 182.5 | 27.6929382119229 |
| | 183 | 27.9122049991076 |
| | 183.5 | 28.1015185496484 |
| | 184 | 28.2560315845642 |
| | 184.5 | 28.3708968248742 |
| | 185 | 28.4412669915971 |
| | 185.5 | 28.4618195749619 |
| | 186 | 28.4253311420381 |
| | 186.5 | 28.324103029105 |
| | 187 | 28.1506899153086 |
| | 187.5 | 27.901437239728 |
| | 188 | 27.5756085759447 |
| | 188.5 | 27.173781913807 |
| | 189 | 26.7071089543274 |
| | 189.5 | 26.1920966507938 |
| | 190 | 25.6452936575293 |
| | 190.5 | 25.0838358505382 |
| | 191 | 24.5269428204549 |
| | 191.5 | 23.9942961288065 |
| | 192 | 23.5055755521565 |
| | 192.5 | 23.0770946057497 |
| | 193 | 22.714806046983 |
| | 193.5 | 22.4226757798693 |
| | 194 | 22.2046643955673 |
| | 194.5 | 22.0616574144814 |
| | 195 | 21.9863104738132 |
| | 195.5 | 21.969916974694 |
| | 196 | 22.0037705308497 |
| | 196.5 | 22.0791978445763 |
| | 197 | 22.1875942812203 |

| | | |
|---|---|---|
| | 197.5 | 22.3203637459707 |
| | 198 | 22.4689101440164 |
| | 198.5 | 22.6252004432081 |
| | 199 | 22.7850323760206 |
| | 199.5 | 22.9457992895612 |
| | 200 | 23.1048997943124 |
| | 200.5 | 23.2597325007567 |
| | 201 | 23.4077317004832 |
| | 201.5 | 23.5477187881055 |
| | 202 | 23.6803170541266 |
| | 202.5 | 23.8062702127852 |
| | 203 | 23.9263219783197 |
| | 203.5 | 24.0412160649685 |
| | 204 | 24.1516961869702 |
| | 204.5 | 24.2585060585633 |
| | 205 | 24.3623863741734 |
| | 205.5 | 24.4640643702727 |
| | 206 | 24.5642635357153 |
| | 206.5 | 24.6637073591985 |
| | 207 | 24.7631193294192 |
| | 207.5 | 24.8632229350744 |
| | 208 | 24.9647416648613 |
| | 208.5 | 25.068399007477 |
| | 209 | 25.1749171389007 |
| | 209.5 | 25.2847536124406 |
| | 210 | 25.3977821509792 |
| | 210.5 | 25.5137984907875 |
| | 211 | 25.6325983681363 |
| | 211.5 | 25.7539775192964 |
| | 212 | 25.8777316805388 |
| | 212.5 | 26.0036565881342 |
| | 213 | 26.1315479783537 |
| | 213.5 | 26.2612015874679 |
| | 214 | 26.3924131517479 |
| | 214.5 | 26.5249784074645 |
| | 215 | 26.6586930908884 |
| | 215.5 | 26.7933529382907 |
| | 216 | 26.9287536859422 |
| | 216.5 | 27.0646910701138 |
| | 217 | 27.2009608270762 |
| | 217.5 | 27.3373597404278 |
| | 218 | 27.4737913310867 |
| | 218.5 | 27.6103564539904 |

| | | |
|---|---|---|
| | 219 | 27.747177304906 |
| | 219.5 | 27.8843760796008 |
| | 220 | 28.0220749738418 |
| | 220.5 | 28.1603961833962 |
| | 221 | 28.2994619040313 |
| | 221.5 | 28.439394331514 |
| | 222 | 28.5803156616115 |
| | 222.5 | 28.7223480900911 |
| | 223 | 28.8656138127198 |
| | 223.5 | 29.0101849372963 |
| | 224 | 29.155855951718 |
| | 224.5 | 29.3023257023844 |
| | 225 | 29.4492929467932 |
| | 225.5 | 29.5964564424419 |
| | 226 | 29.7435149468281 |
| | 226.5 | 29.8901672174496 |
| | 227 | 30.0361283326099 |
| | 227.5 | 30.1812157150808 |
| | 228 | 30.3252863707704 |
| | 228.5 | 30.4681973911347 |
| | 229 | 30.6098058676297 |
| | 229.5 | 30.7499688917111 |
| | 230 | 30.8885435548349 |
| | 230.5 | 31.025386948457 |
| | 231 | 31.1603561640332 |
| | 231.5 | 31.2933082930194 |
| | 232 | 31.4241004268717 |
| | 232.5 | 31.5525896570458 |
| | 233 | 31.6786330749976 |
| | 233.5 | 31.802087772183 |
| | 234 | 31.922824733363 |
| | 234.5 | 32.0408150264009 |
| | 235 | 32.1560735802263 |
| | 235.5 | 32.2686155101811 |
| | 236 | 32.3784559316074 |
| | 236.5 | 32.4856099598471 |
| | 237 | 32.5900927102423 |
| | 237.5 | 32.6919192981351 |
| | 238 | 32.7911048388673 |
| | 238.5 | 32.8876748599706 |
| | 239 | 32.9818058549323 |
| | 239.5 | 33.0737881087213 |
| | 240 | 33.1639146968551 |

| | | |
|---|---|---|
| | 240.5 | 33.2524786948509 |
| | 241 | 33.339773178226 |
| | 241.5 | 33.4260912224978 |
| | 242 | 33.5117259031834 |
| | 242.5 | 33.5969702958003 |
| | 243 | 33.6821174758656 |
| | 243.5 | 33.7674605188968 |
| | 244 | 33.853292500411 |
| | 244.5 | 33.9399064959257 |
| | 245 | 34.027595580958 |
| | 245.5 | 34.1166528310254 |
| | 246 | 34.207371321645 |
| | 246.5 | 34.3000441283342 |
| | 247 | 34.3949643266103 |
| | 247.5 | 34.4924316589409 |
| | 248 | 34.5928857129183 |
| | 248.5 | 34.6968988112616 |
| | 249 | 34.8050485204274 |
| | 249.5 | 34.9179124068717 |
| | 250 | 35.0360521998028 |
| | 250.5 | 35.1597229772684 |
| | 251 | 35.2889021179699 |
| | 251.5 | 35.423556971581 |
| | 252 | 35.5636548877754 |
| | 252.5 | 35.7091632162267 |
| | 253 | 35.8600448874296 |
| | 253.5 | 36.0162319950151 |
| | 254 | 36.1776434941183 |
| | 254.5 | 36.3441982912003 |
| | 255 | 36.5158152926965 |
| | 255.5 | 36.6922920149394 |
| | 256 | 36.8729490753674 |
| | 256.5 | 37.0569899935857 |
| | 257 | 37.2436182891992 |
| | 257.5 | 37.4320374818132 |
| | 258 | 37.6214510910326 |
| | 258.5 | 37.8110626364628 |
| | 259 | 38.0000756377086 |
| | 259.5 | 38.1877691940923 |
| | 260 | 38.3737247238039 |
| | 260.5 | 38.5575992247505 |
| | 261 | 38.7390496948392 |
| | 261.5 | 38.9177331319769 |

| | | |
|---|---|---|
| | 262 | 39.0933065340709 |
| | 262.5 | 39.2654268990281 |
| | 263 | 39.4337512247557 |
| | 263.5 | 39.5979365091607 |
| | 264 | 39.7576397501502 |
| | 264.5 | 39.9125179456313 |
| | 265 | 40.0622280935109 |
| | 265.5 | 40.2064271916963 |
| | 266 | 40.3447722380945 |
| | 266.5 | 40.4769202306125 |
| | 267 | 40.6025281671574 |
| | 267.5 | 40.7212530456363 |
| | 268 | 40.8327518639563 |
| | 268.5 | 40.9366806137514 |
| | 269 | 41.0326031789405 |
| | 269.5 | 41.1199186288372 |
| | 270 | 41.198008923181 |
| | 270.5 | 41.2662560217114 |
| | 271 | 41.3240418841676 |
| | 271.5 | 41.3707484702892 |
| | 272 | 41.4057577398155 |
| | 272.5 | 41.4284516524859 |
| | 273 | 41.43821217 |
| | 273.5 | 41.434421246217 |
| | 274 | 41.4164608467564 |
| | 274.5 | 41.3837129293976 |
| | 275 | 41.33555945 |
| | 275.5 | 41.271382379943 |
| | 276 | 41.1905636673261 |
| | 276.5 | 41.0924852757687 |
| | 277 | 40.9765291650101 |
| | 277.5 | 40.8420772947898 |
| | 278 | 40.6885116248472 |
| | 278.5 | 40.5152141149217 |
| | 279 | 40.3215707570923 |
| | 279.5 | 40.1073240606673 |
| | 280 | 39.8728475866512 |
| | 280.5 | 39.6185795122081 |
| | 281 | 39.3449580145022 |
| | 281.5 | 39.0524212706978 |
| | 282 | 38.7414074579592 |
| | 282.5 | 38.4123547534504 |
| | 283 | 38.0657013343358 |

| | | | | |
|---|---|---|---|---|
| | 283.5 | 37.7018853777796 | | |
| | 284 | 37.321345060946 | | |
| | 284.5 | 36.9245185609992 | | |
| | 285 | 36.5118440551035 | | |
| | 285.5 | 36.0837597204231 | | |
| | 286 | 35.6407037341221 | | |
| | 286.5 | 35.1831142733649 | | |
| | 287 | 34.7115303803131 | | |
| | 287.5 | 34.2279535350452 | | |
| | 288 | 33.7354875388669 | | |
| | 288.5 | 33.237263225692 | | |
| | 289 | 32.7364114294344 | | |
| | 289.5 | 32.2360629840077 | | |
| | 290 | 31.7393487233258 | | |
| | 290.5 | 31.2493994813024 | | |
| | 291 | 30.7693460918513 | | |
| | 291.5 | 30.3023131714594 | | |
| | 292 | 29.8511419187009 | | |
| | 292.5 | 29.4182792282986 | | |
| | 293 | 29.0061428395829 | | |
| | 293.5 | 28.6171504918842 | | |
| | 294 | 28.2537199760995 | | |
| | 294.5 | 27.9183414030984 | | |
| | 295 | 27.6137210346822 | | |
| | 295.5 | 27.3426053402147 | | |
| | 296 | 27.1077407890593 | | |
| | 296.5 | 26.9118738069203 | | |
| | 297 | 26.7573466619271 | | |
| | 297.5 | 26.6451096209192 | | |
| | 298 | 26.5758134890526 | | |
| | 298.5 | 26.5501090714835 | | |
| | 299 | 26.5686471733679 | | |
| | 299.5 | 26.6320785998619 | | |
| | 300 | 26.7410541561216 | | |
| Selenium | 100 | 0.175454545454545 | 1 Ma | Large et al. (2015) |
| | 101 | 0.183636363636364 | | |
| | 102 | 0.191818181818182 | | |
| | 103 | 0.2 | | |
| | 104 | 0.208181818181818 | | |
| | 105 | 0.216363636363636 | | |
| | 106 | 0.224545454545455 | | |
| | 107 | 0.232727272727273 | | |
| | 108 | 0.240909090909091 | | |

| | | |
|---|---|---|
| | 109 | 0.249090909090909 |
| | 110 | 0.257272727272727 |
| | 111 | 0.265454545454545 |
| | 112 | 0.273636363636364 |
| | 113 | 0.281818181818182 |
| | 114 | 0.29 |
| | 115 | 0.298181818181818 |
| | 116 | 0.306363636363636 |
| | 117 | 0.314545454545455 |
| | 118 | 0.322727272727273 |
| | 119 | 0.330909090909091 |
| | 120 | 0.339090909090909 |
| | 121 | 0.347272727272727 |
| | 122 | 0.355454545454546 |
| | 123 | 0.363636363636364 |
| | 124 | 0.371818181818182 |
| | 125 | 0.38 |
| | 126 | 0.333333333333333 |
| | 127 | 0.286666666666667 |
| | 128 | 0.24 |
| | 129 | 0.193333333333333 |
| | 130 | 0.146666666666667 |
| | 131 | 0.1 |
| | 132 | 0.107272727272727 |
| | 133 | 0.114545454545455 |
| | 134 | 0.121818181818182 |
| | 135 | 0.129090909090909 |
| | 136 | 0.136363636363636 |
| | 137 | 0.143636363636364 |
| | 138 | 0.150909090909091 |
| | 139 | 0.158181818181818 |
| | 140 | 0.165454545454545 |
| | 141 | 0.172727272727273 |
| | 142 | 0.18 |
| | 143 | 0.17 |
| | 144 | 0.16 |
| | 145 | 0.15 |
| | 146 | 0.145416666666667 |
| | 147 | 0.140833333333333 |
| | 148 | 0.13625 |
| | 149 | 0.131666666666667 |
| | 150 | 0.127083333333333 |
| | 151 | 0.1225 |

| | | |
|---|---|---|
| | 152 | 0.117916666666667 |
| | 153 | 0.113333333333333 |
| | 154 | 0.10875 |
| | 155 | 0.104166666666667 |
| | 156 | 0.0995833333333333 |
| | 157 | 0.095 |
| | 158 | 0.0904166666666667 |
| | 159 | 0.0858333333333333 |
| | 160 | 0.08125 |
| | 161 | 0.0766666666666667 |
| | 162 | 0.0720833333333333 |
| | 163 | 0.0675 |
| | 164 | 0.0629166666666667 |
| | 165 | 0.0583333333333333 |
| | 166 | 0.05375 |
| | 167 | 0.0491666666666667 |
| | 168 | 0.0445833333333333 |
| | 169 | 0.04 |
| | 170 | 0.0609090909090909 |
| | 171 | 0.0818181818181818 |
| | 172 | 0.102727272727273 |
| | 173 | 0.123636363636364 |
| | 174 | 0.144545454545455 |
| | 175 | 0.165454545454545 |
| | 176 | 0.186363636363636 |
| | 177 | 0.207272727272727 |
| | 178 | 0.228181818181818 |
| | 179 | 0.249090909090909 |
| | 180 | 0.27 |
| | 181 | 0.243 |
| | 182 | 0.216 |
| | 183 | 0.189 |
| | 184 | 0.162 |
| | 185 | 0.135 |
| | 186 | 0.108 |
| | 187 | 0.081 |
| | 188 | 0.054 |
| | 189 | 0.027 |
| | 190 | 0 |
| | 191 | 0.00166666666666667 |
| | 192 | 0.00333333333333333 |
| | 193 | 0.005 |
| | 194 | 0.00666666666666667 |

| | | |
|---|---|---|
| | 195 | 0.00833333333333333 |
| | 196 | 0.01 |
| | 197 | 0.0116666666666667 |
| | 198 | 0.0133333333333333 |
| | 199 | 0.015 |
| | 200 | 0.0166666666666667 |
| | 201 | 0.0183333333333333 |
| | 202 | 0.02 |
| | 203 | 0.183333333333333 |
| | 204 | 0.346666666666667 |
| | 205 | 0.51 |
| | 206 | 0.4976 |
| | 207 | 0.4852 |
| | 208 | 0.4728 |
| | 209 | 0.4604 |
| | 210 | 0.448 |
| | 211 | 0.4356 |
| | 212 | 0.4232 |
| | 213 | 0.4108 |
| | 214 | 0.3984 |
| | 215 | 0.386 |
| | 216 | 0.3736 |
| | 217 | 0.3612 |
| | 218 | 0.3488 |
| | 219 | 0.3364 |
| | 220 | 0.324 |
| | 221 | 0.3116 |
| | 222 | 0.2992 |
| | 223 | 0.2868 |
| | 224 | 0.2744 |
| | 225 | 0.262 |
| | 226 | 0.2496 |
| | 227 | 0.2372 |
| | 228 | 0.2248 |
| | 229 | 0.2124 |
| | 230 | 0.2 |
| | 231 | 0.26 |
| | 232 | 0.32 |
| | 233 | 0.38 |
| | 234 | 0.44 |
| | 235 | 0.5 |
| | 236 | 0.482 |
| | 237 | 0.464 |

| | | |
|---|---|---|
| | 238 | 0.446 |
| | 239 | 0.428 |
| | 240 | 0.41 |
| | 241 | 0.392 |
| | 242 | 0.374 |
| | 243 | 0.356 |
| | 244 | 0.338 |
| | 245 | 0.32 |
| | 246 | 0.302 |
| | 247 | 0.284 |
| | 248 | 0.266 |
| | 249 | 0.248 |
| | 250 | 0.23 |
| | 251 | 0.26 |
| | 252 | 0.315 |
| | 253 | 0.37 |
| | 254 | 0.398421052631579 |
| | 255 | 0.426842105263158 |
| | 256 | 0.455263157894737 |
| | 257 | 0.483684210526316 |
| | 258 | 0.512105263157895 |
| | 259 | 0.540526315789474 |
| | 260 | 0.568947368421053 |
| | 261 | 0.597368421052632 |
| | 262 | 0.625789473684211 |
| | 263 | 0.654210526315789 |
| | 264 | 0.682631578947368 |
| | 265 | 0.711052631578947 |
| | 266 | 0.739473684210526 |
| | 267 | 0.767894736842105 |
| | 268 | 0.796315789473684 |
| | 269 | 0.824736842105263 |
| | 270 | 0.853157894736842 |
| | 271 | 0.881578947368421 |
| | 272 | 0.91 |
| | 273 | 0.866153846153846 |
| | 274 | 0.822307692307692 |
| | 275 | 0.778461538461539 |
| | 276 | 0.734615384615385 |
| | 277 | 0.690769230769231 |
| | 278 | 0.646923076923077 |
| | 279 | 0.603076923076923 |
| | 280 | 0.559230769230769 |

| | | |
|---|---|---|
| | 281 | 0.515384615384615 |
| | 282 | 0.471538461538461 |
| | 283 | 0.427692307692308 |
| | 284 | 0.383846153846154 |
| | 285 | 0.34 |
| | 286 | 0.34 |
| | 287 | 0.34 |
| | 288 | 0.34 |
| | 289 | 0.34 |
| | 290 | 0.34 |
| | 291 | 0.34 |
| | 292 | 0.34 |
| | 293 | 0.34 |
| | 294 | 0.34 |
| | 295 | 0.34 |
| | 296 | 0.34 |
| | 297 | 0.34 |
| | 298 | 0.38875 |
| | 299 | 0.4375 |
| | 300 | 0.48625 |
